# Immunogenicity of COVID-19 vaccines and their effect on the HIV reservoir in older people with HIV

**DOI:** 10.1101/2023.06.14.544834

**Authors:** Vitaliy A. Matveev, Erik Z. Mihelic, Erika Benko, Patrick Budylowski, Sebastian Grocott, Terry Lee, Chapin S. Korosec, Karen Colwill, Henry Stephenson, Ryan Law, Lesley A. Ward, Salma Sheikh-Mohamed, Geneviève Mailhot, Melanie Delgado-Brand, Adrian Pasculescu, Jenny H. Wang, Freda Qi, Tulunay Tursun, Lela Kardava, Serena Chau, Philip Samaan, Annam Imran, Dennis C. Copertino, Gary Chao, Yoojin Choi, Robert J. Reinhard, Rupert Kaul, Jane M. Heffernan, R. Brad Jones, Tae-Wook Chun, Susan Moir, Joel Singer, Jennifer Gommerman, Anne-Claude Gingras, Colin Kovacs, Mario Ostrowski

## Abstract

Older individuals and people with HIV (PWH) were prioritized for COVID-19 vaccination, yet comprehensive studies of the immunogenicity of these vaccines and their effects on HIV reservoirs are not available. We followed 68 PWH aged 55 and older and 23 age-matched HIV-negative individuals for 48 weeks from the first vaccine dose, after the total of three doses. All PWH were on antiretroviral therapy (cART) and had different immune status, including immune responders (IR), immune non-responders (INR), and PWH with low-level viremia (LLV). We measured total and neutralizing Ab responses to SARS-CoV-2 spike and RBD in sera, total anti-spike Abs in saliva, frequency of anti-RBD/NTD B cells, changes in frequency of anti-spike, HIV gag/nef-specific T cells, and HIV reservoirs in peripheral CD4^+^ T cells. The resulting datasets were used to create a mathematical model for within-host immunization. Various regimens of BNT162b2, mRNA-1273, and ChAdOx1 vaccines elicited equally strong anti-spike IgG responses in PWH and HIV^-^ participants in serum and saliva at all timepoints. These responses had similar kinetics in both cohorts and peaked at 4 weeks post-booster (third dose), while half-lives of plasma IgG also dramatically increased post-booster in both groups. Salivary spike IgA responses were low, especially in INRs. PWH had diminished live virus neutralizing titers after two vaccine doses which were ‘rescued’ after a booster. Anti-spike T cell immunity was enhanced in IRs even in comparison to HIV^-^ participants, suggesting Th1 imprinting from HIV, while in INRs it was the lowest. Increased frequency of viral ‘blips’ in PWH were seen post-vaccination, but vaccines did not affect the size of the intact HIV reservoir in CD4^+^ T cells in most PWH, except in LLVs. Thus, older PWH require three doses of COVID-19 vaccine to maximize neutralizing responses against SARS-CoV-2, although vaccines may increase HIV reservoirs in PWH with persistent viremia.

## INTRODUCTION

People with HIV infection (PWH) may be at a higher risk of severe symptoms and mortality related to COVID-19 (1-3). In-depth studies demonstrate a nuanced role of HIV infection in COVID-19 outcomes, arguing that co-morbidities and low CD4^+^ T cell count are better predictors of outcomes than the HIV diagnosis *per se* (4-6). Considering that not all PWH restore their immunity to pre-infection levels despite receiving combination antiretroviral therapy (cART), immunodeficiency may pose a serious risk in the context of COVID-19 disease. Thus, vaccine-mediated prevention of SARS-CoV-2 infection and/or severe disease is essential in this vulnerable population. There is a concern, however, that immunodeficiency may result in impaired responses to COVID-19 immunization, as has been reported for other vaccines in PWH (7-9) and for COVID-19 vaccines in other immunocompromised groups (10-14). The risk of inadequate responses may be even higher for older individuals due to their aging immune system. Since the introduction of cART in 1995, life expectancy of PWH has increased to ages beyond 50 (15). The incidence of HIV infection has also increased in older individuals (16). Given the concern of relative immunodeficiency in treated HIV and the effects of age on immunity, it is unclear whether living with HIV into older age affects responses to COVID-19 vaccines.

Despite cART effectiveness, HIV suppression is often insufficient to completely dampen the heightened immune activation, which itself has a negative impact on morbidity and mortality – but has also been linked to destabilized HIV reservoirs (17). A significant gap in our knowledge is whether COVID-19 vaccines may exacerbate ongoing immune dysregulations that can affect the dynamics of HIV reservoirs, as has been the case with some other vaccines (18-20). Previous work also demonstrated that influenza vaccines could disproportionately increase the HIV proviral burden in influenza-specific CD4^+^ T cells of cART-treated PWH with suppressed viral loads (VL), suggesting the ability of vaccination to affect HIV reservoirs (19). For this reason, understanding how COVID-19 vaccines affect immune activation and HIV reservoirs is paramount.

Given these concerns, we performed a longitudinal observational study to understand how PWH aged 55 and older respond to three COVID-19 vaccine doses across different arms of the immune system, compared to HIV-negative individuals. We also determined whether these vaccines destabilize HIV reservoirs, which has been a major concern for PWH.

## RESULTS

### Study protocol, timeline, participants

Baseline characteristics of study participants and clinical information are provided in Supplemental Tables S1-2, Table 1, and Fig. 1A-B. The study visits are defined in Fig. 1C and Supplemental Methods SM1. In total, we recruited 91 participants: 23 HIV-negative individuals and 68 PWH on cART, including 42 immunological responders, IR (for definitions see SM2), 20 immunological non-responders, INR, five PWH with low-level viremia, LLV, and one long-term non-progressor, LTNP. At the time of screening, all PWH, except the LLVs, had had viremia suppressed to undetectable levels (below 40 copies/mL) for 5-30 years.

**Figure 1.**
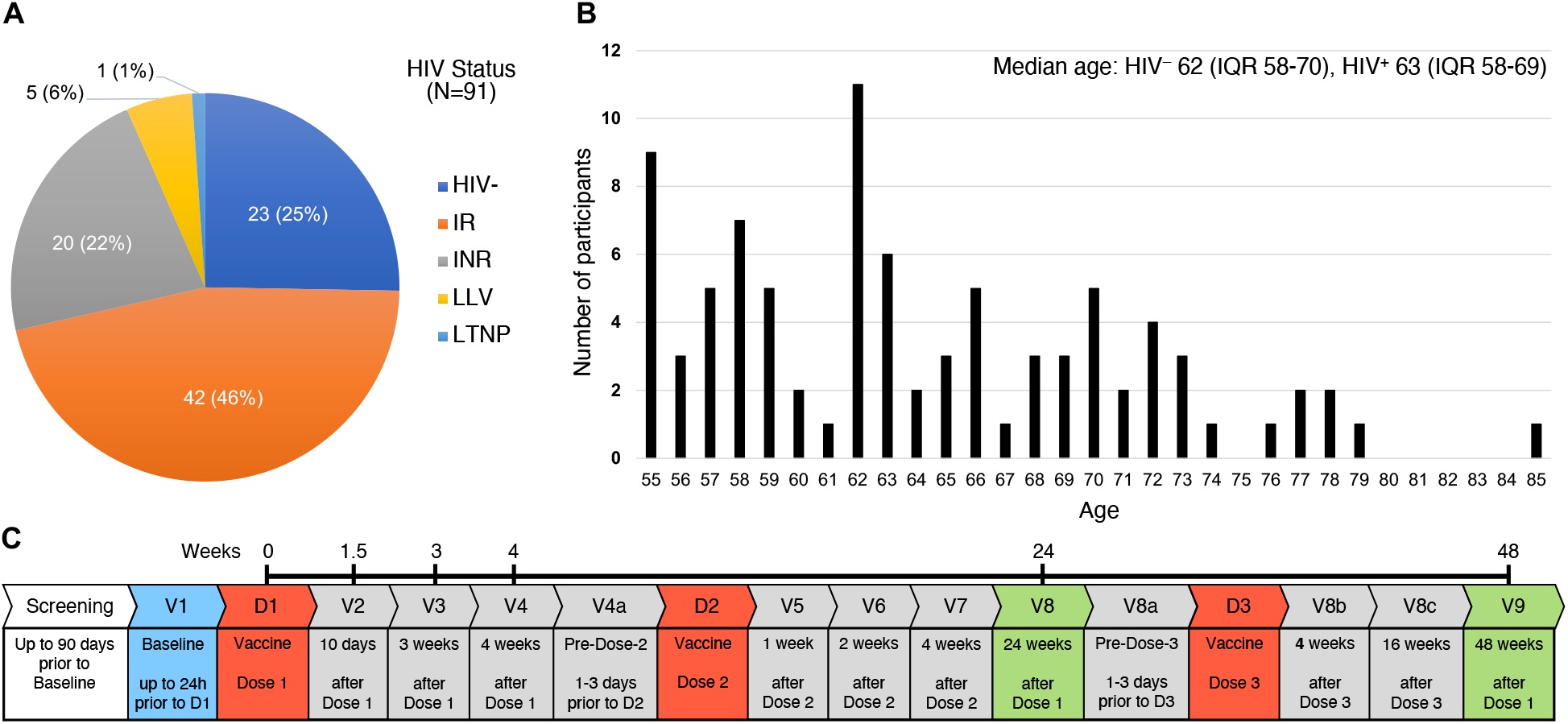
Study protocol, timeline, participants. **(A)** HIV status of study participants. PWH: IR – immunological responders, INR – immunological non-responders, LLV – PWH with low-level viremia, LTNP – long-term non-progressors. **(B)** Age distribution. **(C)** The timeline showing study visits (in blue, gray, or green) and clinically administered COVID-19 vaccine doses D1-3 (red).

**Table 1.** Baseline characteristics of study participants. *P* values are based on Fisher’s exact test for categorical variables and on Wilcoxon sum rank test for continuous variables.

| Variable | HIV- (n=23) | HIV+ (n=68) | *p* | IR<sup>b</sup> (n=42) | INR (n=20) | LLV (n=5) | LTNP (n=1) |
| --- | --- | --- | --- | --- | --- | --- | --- |
| **Age** |  |  | 0.982 |  |  |  |  |
| Median (IQR) | 62.0 (58.0, 70.0) | 63.0 (58.0, 69.0) |  | 62.0 (57.0, 68.0) | 63.0 (60.5, 67.5) | 66.0 (63.0, 70.0) | 76.0 (76.0, 76.0) |
| Range | (55.0, 77.0) | (54.0, 85.0) |  | (54.0, 85.0) | (55.0, 78.0) | (62.0, 73.0) | (76.0, 76.0) |
| **CD4 count<sup>a</sup>** |  |  | – |  |  |  |  |
| Median (IQR) |  | 527 (364, 665) |  | 635 (519, 833) | 316 (252, 410) | 511 (325, 565) | 597 (597, 597) |
| Range |  | (74.0, 1784) |  | (347, 1784) | (74.0, 502) | (322, 764) | (597, 597) |
| **CD4 nadir** |  |  | – |  |  |  |  |
| Median (IQR) |  | 148 (65.0, 240) |  | 210 (110, 290) | 31.0 (10.0, 130) | 91.0 (70.0, 120) | 470 (470, 470) |
| Range |  | (0.0, 1090) |  | (0.0, 1090) | (0.0, 200) | (10.0, 220) | (470, 470) |
| **CD4 percentage<sup>a</sup>** |  |  | – |  |  |  |  |
| Median (IQR) |  | 30.0 (22.0, 38.5) |  | 36.7 (29.8, 42.9) | 19.9 (17.6, 24.8) | 24.7 (17.4, 26.1) | 35.2 (35.2, 35.2) |
| Range |  | (12.9, 62.3) |  | (21.2, 62.3) | (12.9, 30.1) | (15.3, 48.2) | (35.2, 35.2) |
| **CD4/CD8 ratio<sup>a</sup>** |  |  | – |  |  |  |  |
| Median (IQR) |  | 0.9 (0.5, 1.2) |  | 1.1 (0.9, 1.5) | 0.5 (0.4, 0.6) | 0.5 (0.3, 0.5) | 1.0 (1.0, 1.0) |
| Range |  | (0.2, 6.1) |  | (0.4, 6.1) | (0.2, 1.1) | (0.3, 1.9) | (1.0, 1.0) |
| **HIV subgroup, n (%)** |  |  | – |  |  |  |  |
| Immune responders (IR) |  | 42 (61.8) |  |  |  |  |  |
| Immune non-responders (INR) |  | 20 (29.4) |  |  |  |  |  |
| Low-level vireemics (LLV) |  | 5 (7.4) |  |  |  |  |  |
| Long-term non-progressors (LTNP) |  | 1 (1.5) |  |  |  |  |  |
| **Vaccine D1/D2, n (%)** |  |  | 0.766 |  |  |  |  |
| mRNA/mRNA | 13 (56.5) | 35 (51.5) |  | 21 (50.0) | 11 (55.0) | 2 (40.0) | 1 (100) |
| ChAdOx1/ChAdOx1 | 9 (39.1) | 27 (39.7) |  | 17 (40.5) | 7 (35.0) | 3 (60.0) | 0 (0.0) |
| ChAdOx1/mRNA | 1 (4.3) | 6 (8.8) |  | 4 (9.5) | 2 (10.0) | 0 (0.0) | 0 (0.0) |
| **Vaccine D3, n (%)** |  |  | 1.000 |  |  |  |  |
| None | 0 (0.0) | 1 (1.5) |  | 1 (2.4) | 0 (0.0) | 0 (0.0) | 0 (0.0) |
| BNT162b2 | 19 (82.6) | 54 (79.4) |  | 29 (69.0) | 20 (100) | 5 (100) | 0 (0.0) |
| mRNA-1273 | 4 (17.4) | 13 (19.1) |  | 12 (28.6) | 0 (0.0) | 0 (0.0) | 1 (100) |
| **Time lapse between D1 and D2 (days)** |  |  | 0.819 |  |  |  |  |
| Median (IQR) | 63.0 (53.0, 65.0) | 60.0 (54.0, 71.0) |  | 61.0 (53.0, 74.0) | 58.0 (52.5, 67.5) | 65.0 (56.0, 75.0) | 59.0 (59.0, 59.0) |

| Variable | HIV- (n=23) | HIV+ (n=68) | p | IR<sup>b</sup> (n=42) | INR (n=20) | LLV (n=5) | LTNP (n=1) |
| --- | --- | --- | --- | --- | --- | --- | --- |
| Range | (30.0, 78.0) | (28.0, 98.0) |  | (28.0, 92.0) | (28.0, 98.0) | (56.0, 84.0) | (59.0, 59.0) |
| **Time lapse between D3 and V9 (days)** |  |  | 0.076 |  |  |  |  |
| Median (IQR) | 118 (107, 130) | 130 (112, 150) |  | 121 (110, 135) | 152 (138, 165) | 125 (112, 133) | 117 (117, 117) |
| Range | (97.0, 189) | (82.0, 180) |  | (82.0, 179) | (96.0, 179) | (110, 180) | (117, 117) |
| **N+ and/or post-COVID-19, n (%)** | 6 (26.1) | 10 (14.7) |  | 7 (16.7) | 1 (5.0) | 2 (40.0) | 0 (0.0) |
| **First visit number with N+/post-COVID-19, n (%)** |  |  |  |  |  |  |  |
| None | 17 (73.9) | 58 (85.3) |  | 35 (83.3) | 19 (95.0) | 3 (60.0) | 1 (100) |
| V1 | 1 (4.3) | 1 (1.5) |  | 1 (2.4) | 0 (0.0) | 0 (0.0) | 0 (0.0) |
| V4 | 0 (0.0) | 1 (1.5) |  | 0 (0.0) | 1 (5.0) | 0 (0.0) | 0 (0.0) |
| V8 | 0 (0.0) | 1 (1.5) |  | 1 (2.4) | 0 (0.0) | 0 (0.0) | 0 (0.0) |
| V8a | 1 (4.3) | 0 (0.0) |  | 0 (0.0) | 0 (0.0) | 0 (0.0) | 0 (0.0) |
| V8b | 0 (0.0) | 2 (2.9) |  | 1 (2.4) | 0 (0.0) | 1 (20.0) | 0 (0.0) |
| V9 | 4 (17.4) | 5 (7.4) |  | 4 (9.5) | 0 (0.0) | 1 (20.0) | 0 (0.0) |

<sup>a</sup>If baseline data were not available, the data at screening were used (n=24).
<sup>b</sup>IR = PWH who are immunological responders, INR = immunological non-responders, LLV = PWH with low-level viremia, LTNP = long-term non-progressors.

The age of participants ranged between 55 and 85 years, with median age in PWH being 63 (IQR 58-69), and 62 in the HIV^-^ cohort (IQR 58-70). One participant was a female (IR), the rest were males; 79 were Caucasian (86.8%), four were Latino (4.4%), and the rest included Black, Middle Eastern, South Asian (two individuals each, or 2.2%), one East Asian and one Native Brazilian (1.1% each).

As this was an observational study, participants had a variable vaccine regimen of one to three COVID-19 vaccines that had been approved in Ontario, Canada, by the time this study commenced (Tables 1, S1): an adenoviral vector vaccine ChAdOx1 and two mRNA vaccines, BNT162b2 and mRNA-1273 (21-26). We did not observe significant differences in D1 and D2 vaccine regimens between PWH and HIV^-^ (Table 1); all participants received an mRNA vaccine booster (D3). Importantly, the time interval between D1 and D2 was similar for PWH and HIV^-^, as was the one between D3 and V9 (Table 1).

Eighty-nine participants completed the protocol (Table S2). Volunteer OM5128 dropped out after the primary endpoint visit (V8), and CIRC0319 died shortly before the last study visit. This death was not related to the study or vaccination.

### Rates of breakthrough SARS-CoV-2 infection increased with the emergence of Omicron VOC, but were not higher in PWH

Assessment of vaccine efficiency was beyond the scope of this study. In order to discriminate between vaccine-induced changes and those from potential SARS-CoV-2 infection, we determined potentially convalescent samples based on clinically confirmed SARS-CoV-2 infection (by PCR or rapid antigen test) and tested all sera at six study visits with ELISA for anti-N (nucleocapsid) IgG. We did not observe higher rates of breakthrough infection in PWH than in HIV-negative participants – rather, the trend was the opposite (Table 1). One PWH and one HIV^-^ (2.2%) had anti-N IgG in sera at baseline, i.e. had possibly been infected before getting vaccinated. Fourteen more (15.7% of the remaining 89) had a breakthrough infection between D1 and V9, including nine PWH (13.4%) and five HIV^-^ individuals (22.7%). Interestingly, most breakthrough infections occurred after the third vaccine dose (11/16, or 68.8%), i.e. at the time of the spread of the more contagious Omicron variants from late 2021 through early 2022. The samples we presumed convalescent were further considered in sensitivity tests and excluded from some analyses.

### Three doses of COVID-19 vaccines elicit equally high levels of serum anti-spike/RBD IgG that increase with each dose

Neutralizing Abs (nAbs) against SARS-CoV-2 spike protein, particularly its receptor-binding domain (RBD), are a major effector in protection against this infection. We used ELISA assay to assess longitudinally anti-RBD (Fig. 2; Table S3A) and anti-spike (Supplemental Fig. S1; Table S3B) IgG responses in sera at eight timepoints before and after each of the three vaccine doses. At each timepoint, we analysed differences between total PWH and HIV^-^, IR and HIV^-^, INR and HIV^-^, and IR vs INR (Tables S4A, S4B). IgG levels were not normally distributed and were log-transformed to satisfy the required assumption for the regression analysis. Formal comparisons of log*_e_* IgG levels between the groups were based on mixed effects linear regression. We also analysed longitudinal changes in IgG levels within groups and subgroups between adjacent timepoints (Tables S5A, S5B).

**Figure 2.**
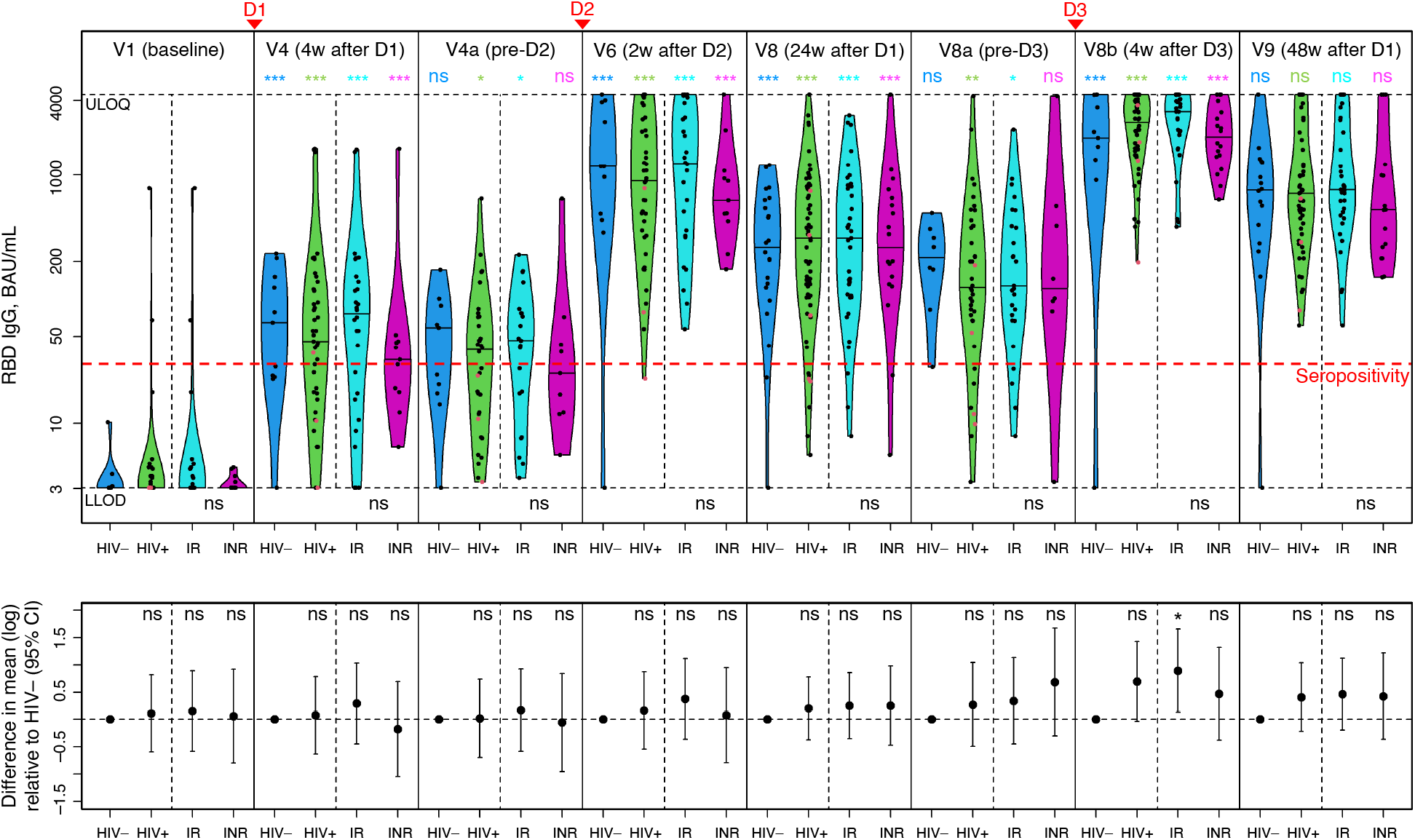
Three doses of COVID-19 vaccines elicit equally high levels of serum anti-RBD IgG that increase with each dose. The top panel shows a violin plot with medians for IgG concentrations (BAU/mL) in HIV^-^ (blue), total PWH (green), IRs (cyan), INRs (purple); N^+^ and/or post-COVID-19 samples are excluded. The bottom panel shows adjusted log*_e_* mean differences between PWH and HIV^-^ individuals, based on mixed effects linear regression. *P* values for log*_e_* mean differences between HIV^+^ and HIV^-^ samples at each timepoint are shown at the top of the bottom panel: *p*<0.001 (***), *p*<0.01 (**), *p*<0.05 (*), *p*≥0.05 (ns = ‘not significant’). *P* values for within-group/subgroup changes between neighbouring timepoints are color-coded accordingly. *P* values for IR vs INR differences are shown at the bottom of the top panel. The lower limit of detection, LLOD (3.02 BAU/mL), the upper limit of quantification, ULOQ (4,454 BAU/mL), and the seropositivity threshold (31 BAU/mL) are shown as dashed horizontal lines. LLV participants are shown as red dots. Vaccination timepoints (D1, D2, D3) are indicated with red arrowheads. See also Fig. S1.

We observed clear RBD and spike IgG responses in PWH and HIV^-^ participants that were boosted by each subsequent dose (Fig. 2, S1), with little difference in Ab levels between the two groups (Tables S4A-B). For RBD IgG, the median baseline levels were below the seropositivity threshold of 30.97 BAU/mL in all groups, while at 4 weeks after D1 (V4), the median value in PWH went up to 45.6 BAU/mL (V4 mean vs V1 *p*<0.001) and 64.5 in HIV^-^ (V4 mean vs V1 *p*<0.001). At 2 weeks post-D2, the median levels jumped to 893 in PWH (V6 mean vs V4a *p*<0.001) vs 1,169 in HIV^-^ (V6 mean vs V4a *p*<0.001). Finally, the RBD IgG peaked at 4 weeks after D3 (V8b) at 2,655 BAU/mL in PWH (V8b mean vs V8a *p*<0.001) vs 1,983 in HIV^-^ (V8b mean vs V8a *p*<0.001). The differences between PWH and HIV^-^ means at these three timepoints were not statistically significant (Fig. 2; Tables S3A, S4A, S5A). The same pattern was observed for spike IgG (Fig. S1; Tables S3B, S4B, S5B). Of note, at V8b, RBD IgG responses in IRs were significantly higher than in the HIV^-^ group, with adjusted log difference in mean being 0.89 (*p*=0.022, with N^+^/post-COVID-19 samples excluded), which was the only significant difference in serum IgG responses that we observed between PWH and HIV^-^ participants.

An increase in serum IgG levels post-vaccine was followed by decay, as seen at V4a and V8-8a (Fig. 2, S1). Interestingly, the change from the peak values at V8b to the last study visit (V9) suggest a slower decay of Abs following D3 – see the model below for more details.

No significant statistical differences were observed between IRs and INRs, although INRs showed a trend for lower IgG levels at most timepoints.

At 48 weeks (V9), after three vaccine doses, we registered the following median IgG concentrations in sera: for spike, 916 BAU/mL in PWH vs 919 in HIV^-^ (*p*=0.624), and for RBD, 706 BAU/mL in PWH vs 752 in HIV^-^ (*p*=0.198).

### PWH had lower SARS-CoV-2 nAb titers after two vaccine doses than HIV^-^ individuals, with responses ‘rescued’ by the booster

We performed live SARS-CoV-2 microneutralization (MN) to assess the neutralization capacity of sera following COVID-19 vaccination of PWH and HIV^-^ individuals at baseline, V8 (after two vaccine doses) and V9 (after three vaccine doses). Baseline 50% neutralization titers (NT_50_) were summarized as dichotomized variables (0 vs >0) given that most data were zero (Table S6). As expected, no differences were observed at V1. Post-baseline data were non-normal even after log-transformation, and the intergroup/subgroup comparisons were based on adjusted quantile regression (Tables S7-8).

In contrast to spike and RBD IgG responses, at 24 weeks, we observed markedly lower SARS-CoV-2 neutralization titers in PWH vs HIV^-^ participants after two COVID-19 vaccine doses (*p*<0.001). The median NT_50_ in PWH was 82.9 vs 535 in HIV^-^, with N^+^/post-COVID-19 samples excluded (Fig. 3A; Tables S7-8).

**Figure 3.**
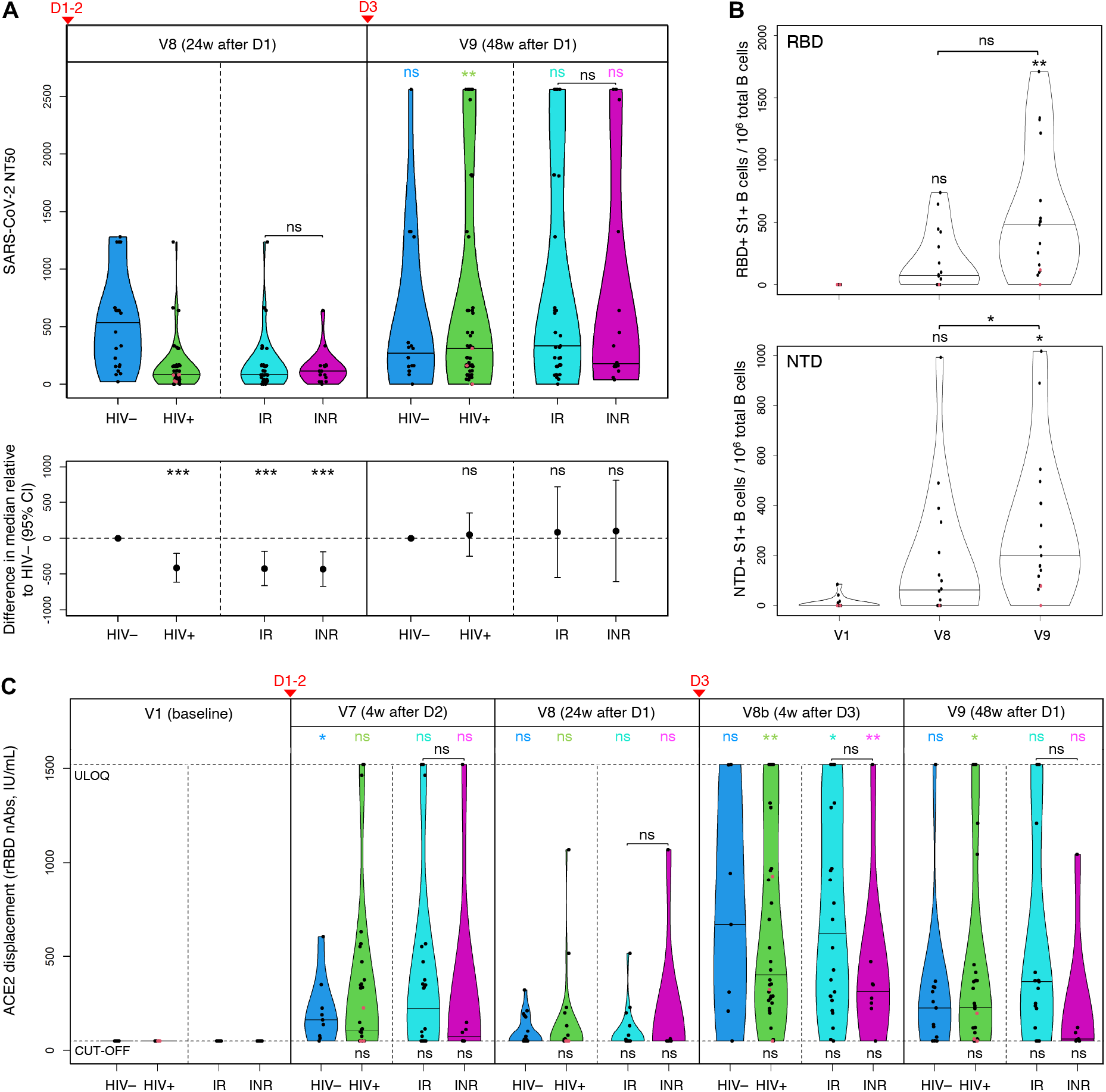
PWH had lower SARS-CoV-2 nAb titers after two vaccine doses than HIV^-^ individuals, with responses ‘rescued’ by the booster. **(A)** Live SARS-CoV-2 50% neutralization titers (NT_50_) in HIV^-^ individuals (blue), total PWH (green), IRs (cyan), INRs (purple); the horizontal bars show median titers. The bottom panel shows adjusted median differences between PWH and HIV^-^ participants, based on quantile regression, with *P* values at the top: *p*<0.001 (***), *p*<0.01 (**), *p*<0.05 (*), *p*≥0.05 (ns). *P* values for within-group/subgroup changes from V8 to V9 are color-coded accordingly. Vaccination timepoints (D1-2, D3) are indicated with red arrowheads. **(B)** Changes in frequency of RBD/NTD-specific B cells in selected PWH (n=17) following COVID-19 vaccination expressed as the number of RBD^+^S1^+^ and NTD^+^S1^+^ B cells per 10^6^ total B cells, based on spectral flow cytometry data. The horizontal bars show median frequencies. *P* values are based on quantile regression. See also Fig. S2A-B. **(C)** Changes in concentrations of anti-RBD nAbs in sera (IU/mL) following COVID-19 vaccination: in HIV^-^ participants (blue), total PWH (green), IRs (cyan), INRs (purple). The responses are measured with snELISA based on the ability of sera to displace rACE2 and analyzed by adjusted quantile regression. The horizontal bars show median concentrations; the neutralization cut-off (54.8 IU/mL) and ULOQ (1,520 IU/mL) are shown as dashed horizontal lines. *P* values for differences between PWH and HIV^-^ participants are shown at the bottom. *P* values for within-group/subgroup changes between neighbouring timepoints are color-coded accordingly. Vaccination timepoints (D1-2, D3) are indicated with red arrowheads. **(A-C)** LLV participants are shown as red dots. See also Fig. S2C.

At 48 weeks, the pattern changed considerably, with PWH responses having been ‘rescued’ by the D3 booster: the median NT_50_ value increased in PWH from 82.9 to 309 (*p*=0.009); the change in HIV^-^ was not statistically significant (*p*=0.858). The adjusted median difference between PWH and HIV^-^ individuals, as a result, decreased from -413 at V8 (*p*<0.001) to 49.1 at V9 (*p*=0.745).

The difference between IR and INR was not significant at V8. The V9 increase, however, was more pronounced in the former, albeit not significant. Therefore, IRs contributed the most to the ‘rescue’ of PWH responses post-D3 (Table S7): median 332 in IR vs 177 in INR. The adjusted median difference between IRs and INRs at V9 had a wide 95% CI and was not significant (Table S8). LLVs and our only LTNP showed approximately a two-fold increase in their NT_50_ titers from V8 to V9 (Table S7). However, formal comparisons involving these subgroups were not performed due to the small sample size.

### Third vaccine dose boosts spike-specific B cells

The discrepancy that we observed in PWH between total spike/RBD IgG responses and neutralizing titers in sera prompted us to evaluate vaccine-induced B cell responses. We focused on circulating B cells targeting RBD and NTD (N-terminal domain) – two domains within the S1 subunit of the SARS-CoV-2 spike that represent mutational hotspots and antigenic supersites for nAbs (27-32). The responses were measured by spectral flow cytometry at baseline, V8 and V9 in selected PWH with similar D3-V9 intervals (n=17) and reported as frequencies: the number of RBD^+^S1^+^ and NTD^+^S1^+^ B cells per 10^6^ total B cells (Table S9; Fig. 3B). HIV-negative participants were not included due to a limited number of specimens collected from this cohort.

Similarly to the NT_50_ titers, we only observed a moderate increase in frequencies of RBD^+^S1^+^ and NTD^+^S1^+^ B cells from V1 to V8 (Fig. 3B; Table S9). The third dose significantly boosted the RBD^+^S1^+^ and NTD^+^S1^+^ B cells, with median frequencies increasing from 73.6 at V8 to 480 at V9 for RBD, and from 62.6 to 200 for NTD. The change from baseline to V9 was significant for both domains (*p*=0.008 and *p=*0.034, respectively), whereas the change from V8 to V9 was only significant for the NTD^+^S1^+^ B cells (*p*=0.036).

Frequencies of RBD^+^S1^+^ and NTD^+^S1^+^ B cells showed moderate positive correlation with NT_50_ titers (Fig. S1A,B): ρ=0.46 and ρ=0.54, respectively (*p*<0.001). Interestingly, the only participant without either response (RBD or NTD) at V9 was the LLV individual OM5094 who also had negligible or undetectable NT_50_ titers at V8 and V9.

### RBD surrogate neutralization ELISA reaffirms the benefits of the D3 booster, points to weaker responses in INRs

We performed snELISA to estimate longitudinal changes in the concentration of anti-RBD nAbs based on the capacity of sera to prevent the binding of the labelled ACE2 receptor (the mediator of the SARS-CoV-2 cell entry) to immobilized RBD (33, 34).

All V1 data were below the neutralization cut-off of 54.8 IU/mL (Tables S10-11; Fig. 3C). In line with assays described above, the boosting effect from D3 was confirmed by the ACE2 displacement. With regards to responses in PWH at V8 and V9, the observed pattern was similar to that of NT_50_ titers, with much higher nAb concentrations at V9, which reaffirms the critical role of the booster. Likewise, we note little difference in median V9 values between PWH and HIV^-^ participants: 232 IU/mL vs 228, respectively. Spearman correlation test for snELISA and NT_50_ datasets for the two timepoints (Fig. S2C) showed a moderate positive correlation for the V8 dataset (ρ=0.56, *p*<0.001) and strong for V9 (ρ=0.79, *p*<0.001).

Higher responses at V7 vs V8, and V8b vs V9 demonstrate a relatively fast decay of RBD nAbs, particularly after D1-2. Neutralizing Abs targeting RBD peaked at 4 weeks post-D3 (V8b), and the difference between PWH and HIV^-^ was not statistically significant (Tables S10-11): the median value in PWH increased from undetectable at V8 to 403 IU/mL at V8b (*p*=0.007).

We did not observe statistically significant differences between IRs and INRs, but a clear trend for weaker median responses in the latter was evident: 225 IU/mL vs 73.0 at V7, 621 vs 315 at V8b, and 367 vs 59.9 at V9, respectively. Interestingly, the responses in the LLV participants were among the weakest, often undetectable (Fig. 3C; Table S10).

### Strong anti-spike IgG responses to three mRNA vaccine doses in saliva contrast with weak anti-spike IgA responses, with little effect from the booster on IgA

Mucosal immunity of the upper respiratory tract and oral cavity, particularly its IgA component, is a first-line defence against SARS-CoV-2, but one with many unknowns (35, 36). We performed ELISA to look at longitudinal changes in anti-spike IgG and IgA levels in saliva at several timepoints following three doses of mRNA COVID-19 vaccines in 31 participants (Tables S12-13). The results were expressed as %AUC (area under the curve) of the AUC of a pooled positive control of acute and convalescent saliva. Comparisons were based on Wilcoxon rank sum test (intergroup) or quantile regression (within-group/subgroup). Individuals who had at least one dose of adenoviral vaccine were not included, as such vaccines apparently elicit weaker mucosal anti-SARS-CoV-2 immunity when administered intramuscularly (37).

The median baseline levels of anti-spike IgG in the HIV^-^ group, 0.65%AUC, were just above the positive cut-off of 0.35%AUC (Table S12A), being significantly higher than in PWH (*p*=0.025). Of note, we only analysed saliva from seven HIV^-^ participants due to a limited number of mRNA-vaccinated HIV^-^ age-matched individuals. Post-baseline IgG responses to each vaccine dose were quite strong in PWH and HIV^-^ (Fig. 4, top), without significant differences between the two groups (Tables S12A, S13A). The IgG levels peaked at 4 weeks post-D3 (V8b) at 119%AUC in PWH and 136%AUC in HIV^-^, decreasing by V9 to median 48.1%AUC in PWH (*p*=0.023), whereas the HIV^-^ decrease to 95.9%AUC was not significant.

**Figure 4.**
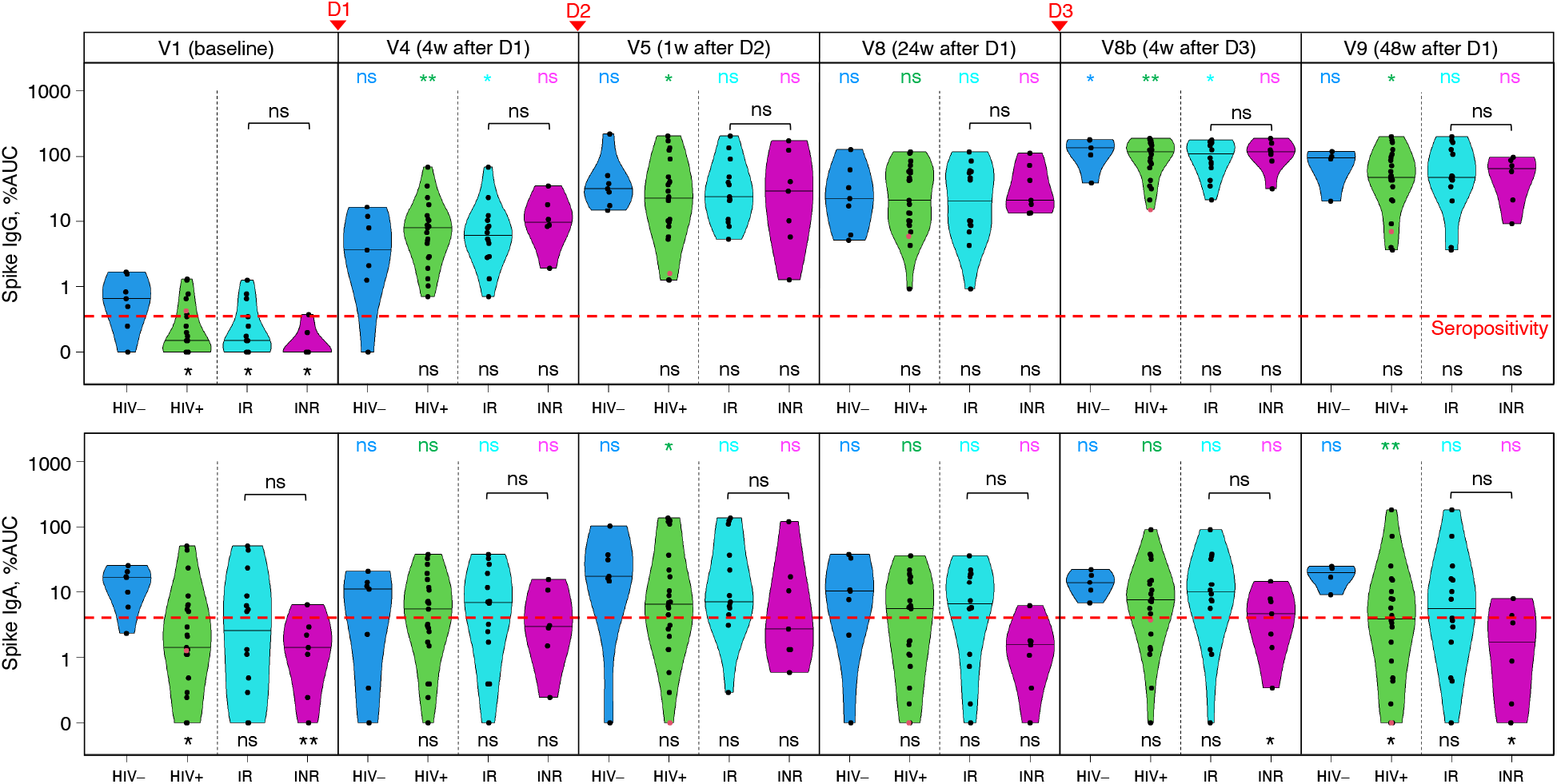
Strong anti-spike IgG responses to three mRNA vaccine doses in saliva contrast with weak anti-spike IgA responses, with little effect from the booster on IgA. Longitudinal changes in levels of spike-specific IgG (top) and IgA (bottom) Abs in saliva following COVID-19 vaccination in HIV^-^ participants (blue), total PWH (green), IRs (cyan), INRs (purple). The values are expressed as %AUC (percentage of the area under the curve of a positive control); the horizontal bars show median responses. *P* values for differences between PWH and HIV^-^ participants, and those between IRs and INRs, are based on Wilcoxon rank sum test and are shown below their respective violin plots or above each IR/INR pair: *p*<0.001 (***), *p*<0.01 (**), *p*<0.05 (*), *p*≥0.05 (ns). *P* values for within-group/subgroup changes between neighbouring timepoints are based on quantile regression and are color-coded. LLV participants are shown as red dots. Seropositivity thresholds, 0.35%AUC for IgG and 4.06%AUC for IgA, are shown as red dashed lines. Vaccination timepoints (D1, D2, D3) are indicated with red arrowheads.

Unexpectedly, we observed relatively high baseline anti-spike IgA responses in HIV^-^ participants, with a median of 17.1%AUC – well above the positive cut-off of 4.06%AUC (Fig. 4, bottom; Table S12B). In fact, IgA levels in this group remained at similarly high levels at all timepoints, with medians ranging between 10.3%AUC at V8 and 20.5%AUC at V9 (Table S13B). High baseline levels of anti-spike IgA (and to a lesser extent IgG) differed from previously studied younger cohorts (36, 38). In PWH, the V1 IgA responses did not exceed the cut-off. Consistent with previous findings (36), the IgA responses in PWH were modest relative to the positive control, with median values slightly exceeding the seropositivity threshold at V4, V5, and V8, peaking at 7.62%AUC at V8b. At the last study visit (V9), the IgA levels in PWH dropped below the cut-off to 3.83%AUC (*p*=0.007), compared to 20.5%AUC in HIV^-^ (*p*=0.039 for PWH vs HIV^-^).

We observed a trend for weaker spike IgA responses in INRs, compared to IRs, but these differences were not statistically significant (Fig. 4, bottom).

### IRs mount strong T cell responses to two and three vaccine doses, outperforming HIV^-^ individuals, while INRs show modest responses

We used cytokine enzyme-linked immune absorbent spot assay (ELISpot) to compare spike-specific T cell immunity in PWH and HIV^-^ participants following three doses of COVID-19 vaccines at baseline, 24 and 48 weeks post-D1 (V8, V9). Because T-cell spike-specific responses demonstrate a strong Th1 shift (39), we measured them as the frequency of cells that secrete IFN-γ, IL-2 or both – ‘dual’ (Fig. 5), following stimulation with spike peptides. The values were expressed as the number of spot-forming cells (SFC) per 10^6^ PBMCs.

**Figure 5.**
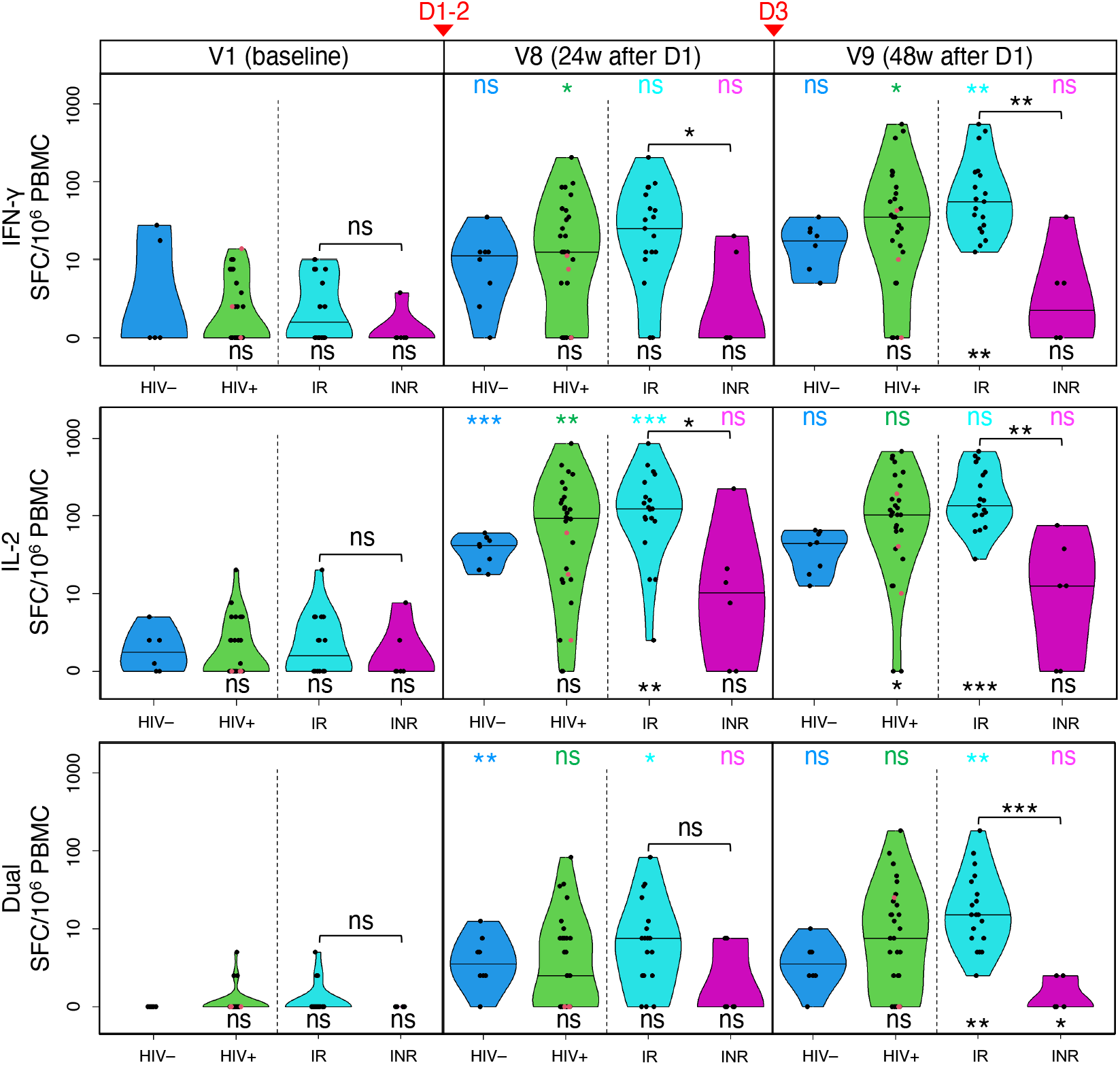
IRs mount strong T cell responses to two and three vaccine doses, outperforming HIV^-^ individuals, while INRs show modest responses. Changes in anti-spike T cell responses in HIV^-^ participants (blue), total PWH (green), IRs (cyan), and INRs (purple) following COVID-19 vaccination are determined with ELISpot as frequencies of cells secreting IFN-γ (top), IL-2 (middle) or both (‘Dual’, bottom) in response to stimulation with SARS-CoV-2 spike peptide pool and expressed as spot-forming cells (SFC) per 10^6^ PBMC. The horizontal bars show median frequencies. *P* values for differences between PWH and HIV^-^ participants, and those between IRs and INRs, are based on Wilcoxon rank sum test and are shown below their respective violin plots or above each IR/INR pair: *p*<0.001 (***), *p*<0.01 (**), *p*<0.05 (*), *p*≥0.05 (ns). *P* values for within-group/subgroup changes between neighbouring timepoints are based on quantile regression and are color-coded. LLV participants are shown as red dots. Vaccination timepoints (D1-2, D3) are indicated with red arrowheads.

The cytokine responses differed in magnitude: IL-2>IFN-γ>dual (Tables S14-15). For IL-2, we observed a dramatic shift in median SFC frequency from baseline to V8: zero to 92.5 in PWH (*p*=0.005), and 1.9 to 41.3 in HIV^-^ (*p*<0.001). For IFN-γ, the change was more subtle: zero to 12.5 in PWH (*p*=0.02), and zero to 11.3 in HIV^-^ (not significant). The dual responses at V8 only showed a minor change from baseline.

Dose 3 boosted T-cell responses further, but the change from V8 was moderate, compared to the change from V1 to V8 (Fig. 5). Perhaps, most notable were the stronger responses in PWH compared to HIV^-^ participants – at V8 and V9 for IL-2, and at V9 for IFN-γ and dual, with similar observations also made recently with flow cytometry(40). The difference between IRs and HIV^-^ was even more profound (Table S14; Fig. 5), as were the changes within IRs (Table S15; Fig. 5). The median IL-2 responses at V8 were 123 in IR vs 41.3 in HIV^-^ (*p*=0.009), vs 92.5 in total PWH (*p*=0.196). The V9 difference was even greater: 135 in IR vs 43.8 in HIV^-^ (*p*<0.001), vs 103 in total PWH (*p*=0.027). The same pattern was observed for IFN-γ and dual.

INRs had significantly lower responses than IRs at V8 and V9 for both cytokines. For IL-2, the means compared as 10.6 in INRs vs 92.5 in IRs at V8 (*p*=0.016); at V9, 12.5 in INRs vs 135 in IRs (*p*=0.001). INRs also showed a trend for weaker responses than in HIV^-^ individuals.

### The size of intact HIV reservoir in peripheral CD4^+^ T cells does not increase after three COVID-19 vaccine doses, with a possible exception of PWH with unsuppressed viremia

We used IPDA, or Intact Proviral DNA Assay (41), to assess the influence of two (V8) and three (V9) doses of COVID-19 vaccines on the size of HIV reservoir measured as the number of intact HIV proviruses per 10^6^ CD4^+^ T cells and analyzed by quantile regression. The pattern and magnitude of changes varied among PWH (Fig. 6, left). For total PWH, the median frequency of intact proviruses changed from 90.9 at baseline to 122 at V8, and to 95.0 at V9 (Table S16). Median changes between the three timepoints were subtle and statistically not significant.

**Figure 6.**
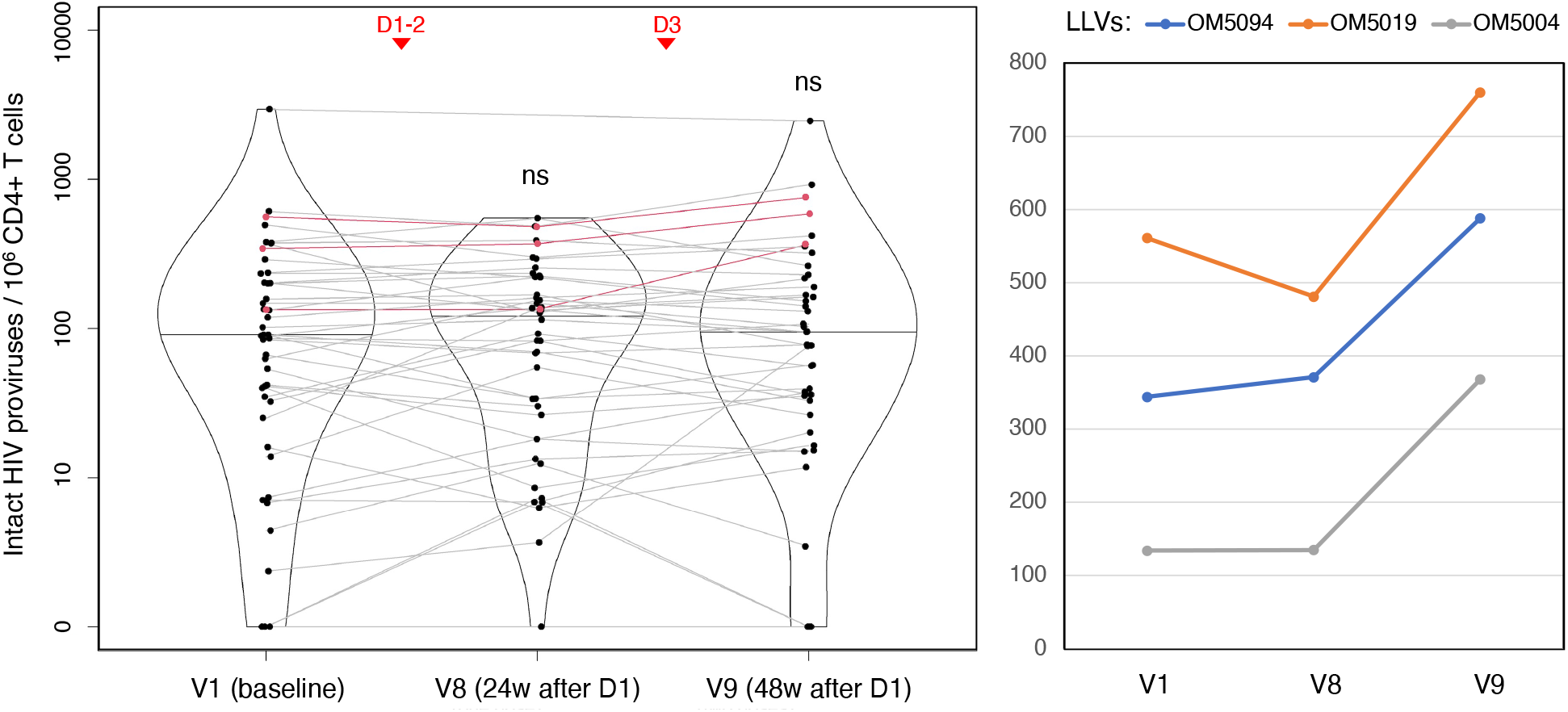
The size of intact HIV reservoir in peripheral CD4^+^ T cells does not increase after three COVID-19 vaccine doses, with a possible exception of PWH with unsuppressed viremia. The violin plot (left) shows IPDA-derived frequencies of intact HIV proviruses (with medians) per 10^6^ CD4^+^ T cells analyzed by quantile regression. The dots representing the same person are connected, and LLV participants are shown in red; ns = ‘not significant’ (*p*≥0.05). Vaccination timepoints (D1-2, D3) are indicated with red arrowheads. IPDA values for LLV participants on the original scale.

No significant changes or differences were observed within or between IRs and INRs. By contrast, we saw a dramatic increase in the frequency of intact HIV proviruses from baseline to V9 in all three LLV participants tested: by 35.5% in OM5019, 70.9% in OM5094, and 175% in OM5004, which is a mean increase of 93.7% (Fig. 6, right).

Destabilization of the intact HIV reservoir may trigger viral replication, which could then result in detectable VL. For most timepoints in this study, the number of VL ‘blips’ above 40 copies/mL was limited to one or two participants (Table S17; Fig. S3). After D2, however, we observed an increase from one such case at V5 (1/39, or 2.6%) to three at V6 (3/40, or 7.5%), and five at V7 (5/43, or 11.6%), with most cases being IRs. Only one of the latter five participants had a noticeable IPDA increase at V8. In total, we observed 24 such blips in 14 non-LLV PWH post-D1 over the course of the study. Only three of them (OM5085, OM5005, CIRC0041) had episodes of detectable HIV viremia over the period of two years preceding the study – one ‘blip’ per person. It should be noted that the frequency of tests was lower before the study and varied among participants.

### HIV gag/nef-specific T cells do not change in frequency following COVID-19 vaccination

As another measure of potential changes in the HIV reservoir, we assessed frequencies of gag- and nef-specific T cells by ELISpot, estimating a number of cells secreting IFN-γ, IL-2 or both (dual) following stimulation with gag or nef peptide pools and expressed as SFC per 10^6^ PBMC (Fig. S4). Contrary to the recent finding of increased frequencies of nef-specific CD8^+^ T cells in a cohort of 13 cART-treated PWH following two doses of the BNT162b2 mRNA vaccine (42), we did not observe such changes for nef- or gag-specific T cells after two or three vaccine doses, even after excluding non-mRNA vaccine data (not shown). None of the changes were significant, including those at the level of subgroups (Table S18).

### CD4^+^ T cell count in PWH increased transiently after the primary vaccine dose

The only significant change in CD4-related clinical parameters that we observed in PWH throughout the study and which is noteworthy was a transient increase in CD4^+^ T cell count at V2, or 10 days post-D1 (Fig. S5-7): the median count increased from 527 cells/μL at baseline to 597 at V2 (Table S19). This is consistent with an expectation of immune activation following the primary vaccine dose. The mean change from V1 to V2 was even more pronounced in IRs (Tables S20-21): 47.9 for total PWH (*p*=0.008) vs 73.0 for IR (*p*=0.001).

For CD4^+^ T cell percentage (Fig. S7), we observed a small but statistically significant transient increase in INRs at 4 weeks post-D3 (V8b), with estimated mean change of 1.7% from V8a (*p*=0.013). No changes in CD4^+^/CD8^+^ ratio were statistically significant (Tables S20-21).

### Longitudinal modelling of humoral spike/RBD-specific IgG responses

Figure 7 details longitudinal model-fit responses for serum anti-spike and anti-RBD IgG, as well as RBD^+^S1^+^ and NTD^+^S1^+^ B cells, sorted by HIV status. Overall, little difference in median responses was found (Table S22). For individual fit examples, see Figure S8.

**Figure 7.**
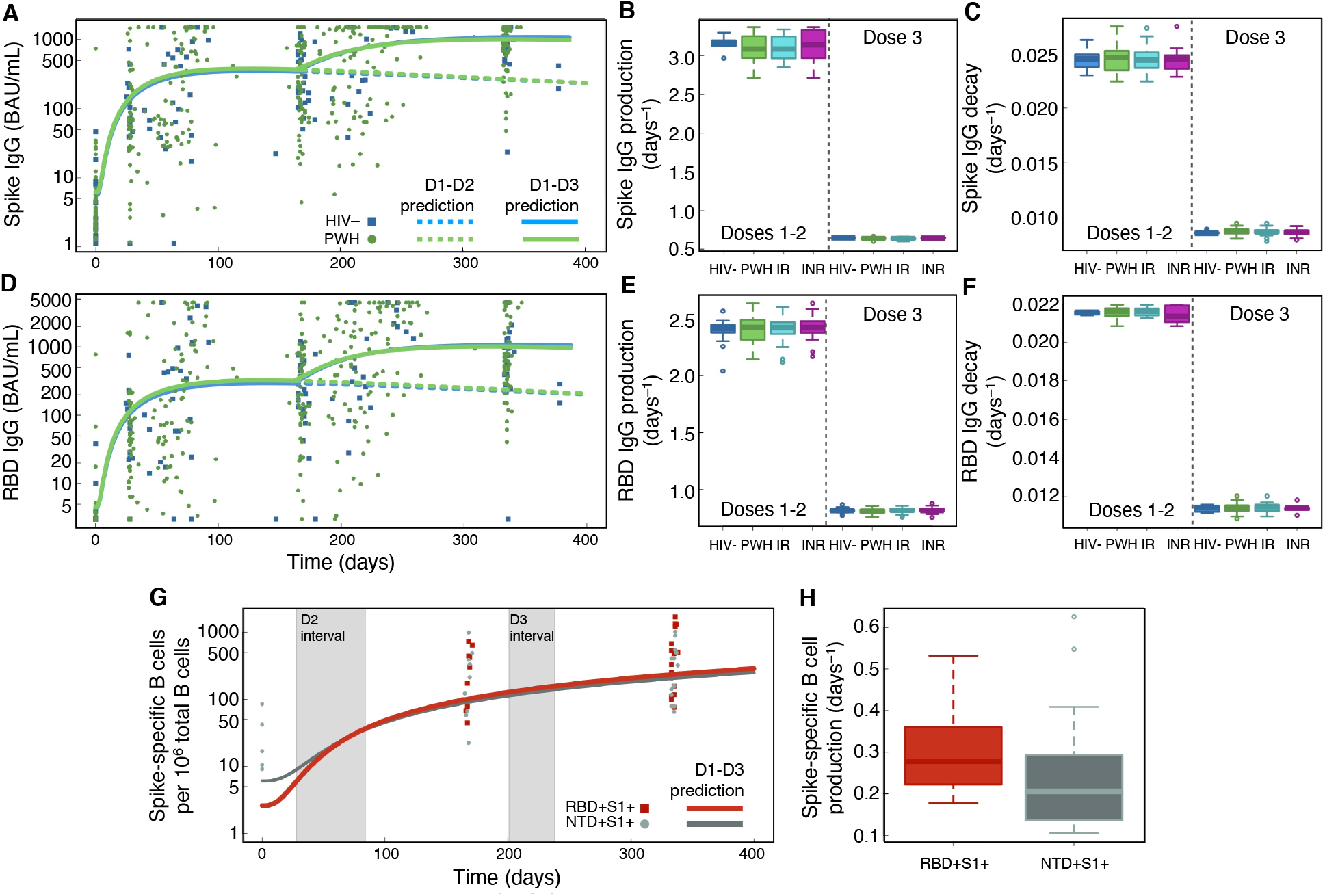
Longitudinal modelling of humoral spike/RBD-specific IgG responses. **(A)** Serum anti-spike IgG longitudinal model (Eq. 1d) mean fit prediction as a function of HIV status. Solid lines show IgG dynamics from the primary COVID-19 vaccine series (D1-2) and booster (D3). Dashed lines extend the trend in IgG decay from D1-2. **(B-C)** Model-predicted spike IgG production rates, *μA_spike_* **(B)**, and decay rates, *γA_spike_* **(C)**. **(D)** Serum anti-RBD IgG longitudinal model (Eq. 1e) mean fit prediction as a function of HIV status. Also see Fig. S8. **(E-F)** Model-predicted RBD IgG production rates, *μA_RBD_* **(E)**, and decay rates, *γA_RBD_* **(F)**. **(G)** Longitudinal model (Eqs. 1f,g) mean fit predictions for RBD^+^S1^+^ and NTD^+^S1^+^ B cells, respectively, in PWH. Shaded regions illustrate the range in dosage time intervals where the B cell data were measured. **(H)** A boxplot of individual model-predicted production rates for spike-specific B cells.

For PWH and HIV^-^ participants, spike IgG production for D1-D2 was almost identical, 3.18 and 3.17 BAU/mL·d, respectively, and decreased to 0.643 and 0.625 BAU/mL·d post-D3. RBD IgG production for D1-D2 is 2.4 BAU/mL·d and decreased post-D3 to 0.81 BAU/mL·d. IgG spike and RBD half-life values for D2 are 28.9 ± 7.2 and 32.1 ± 4.3 days, respectively, where errors are the relative standard error (Table S22). Estimated half-lives of spike and RBD IgG increased substantially post-D3 to 80 ± 22.4 and 63 ± 15 days, respectively. For distributions of serum IgG production and decay parameter estimates, see Fig. 7B-C (spike) and Fig. 7E-F (RBD).

Figure 7G shows mean longitudinal fit results for RBD^+^S1^+^ and NTD^+^ S1^+^ B cells in PWH: both are found to strictly increase over the study period of ∼340 days and display no decay in response. Thus, no estimate of half-life of B cells is possible in this study. The median production rate for RBD^+^S1^+^ B cells was higher, 0.28 d^-1^, vs 0.21 d^-1^ for NTD^+^S1^+^ (Fig. 7H). From equations Eqs. 1f,g we can also estimate the doubling time, *T_D_*, defined as the estimated time it takes for a population to double in size at a particular time. For NTD^+^S1^+^, we estimate *T_D_* to be 40 days post-D1, 97 days post-D2, and 377 days post-D3.

### Longitudinal modelling of cytokine responses in T cells

Figure 8 details longitudinal model-fit responses for IFN-γ and IL-2. Total PWH and IR IFN-γ mean responses are elevated relative to HIV^-^ (Fig. 8A). In contrast, INR IFN-γ mean responses are found to fall slightly below the HIV^-^ response. PWH and IR IL-2 mean responses are elevated, whereas INR IL-2 responses fall slightly below the HIV^-^ response (Fig. 8B).

**Figure 8.**
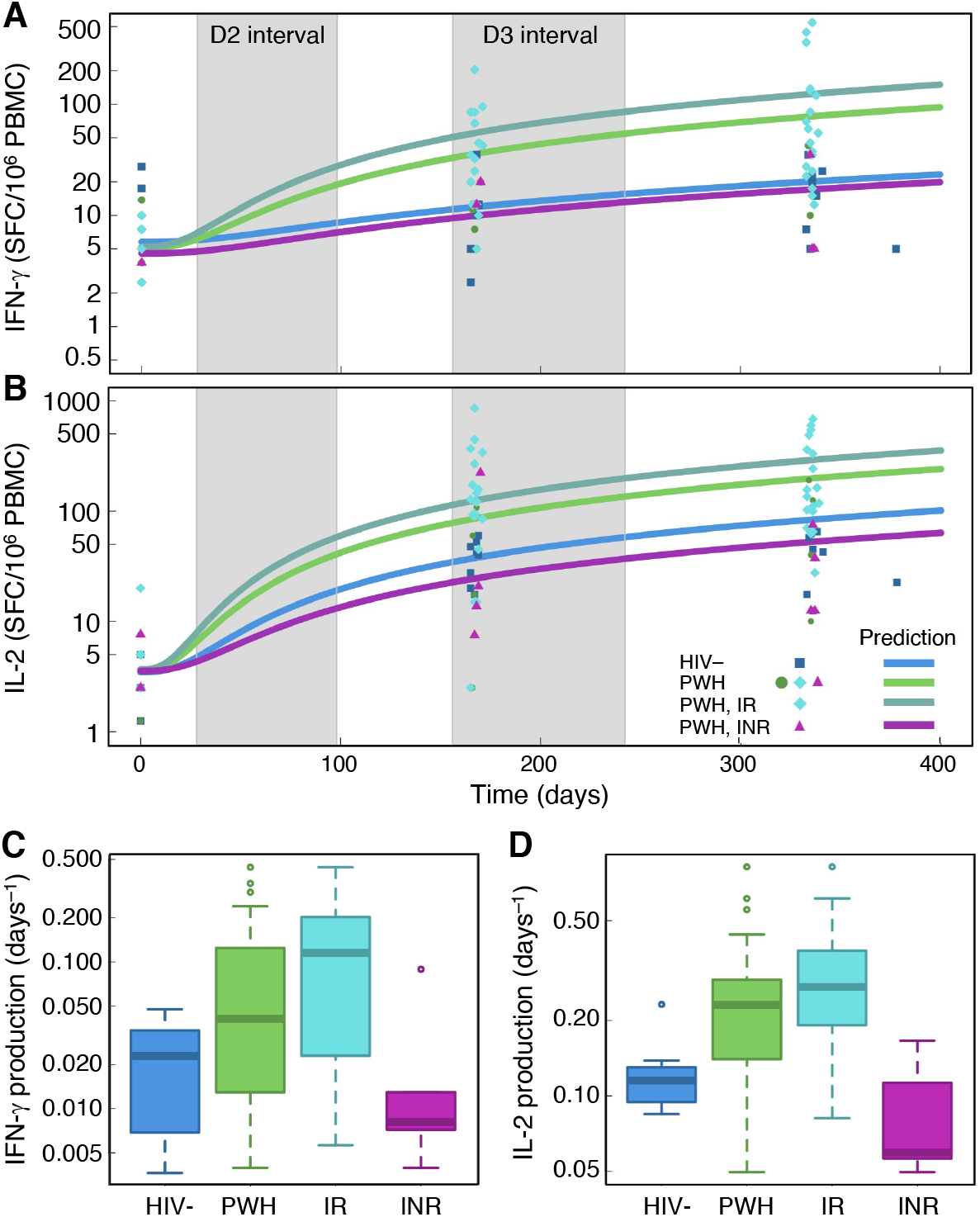
Longitudinal modelling of cytokine responses in T cells. **(A-B)** Longitudinal model (Eqs. 1g,h) mean fit predictions for anti-spike cytokine T cell responses as a function of HIV status. Shaded regions are the dosage time intervals across all individuals. Panels **(A)** and **(B)** are IFN-γ (Eq. 1g), and IL-2 (Eq. 1h) three-dose predictions as a function of days since COVID-19 vaccine D1, respectively. **(C-D)** Boxplots of production rate model estimates sorted by HIV status for IFN-γ **(C)** and IL-2 **(D)**. Also see Fig. S8.

Figure 8C,D displays boxplots of individual IFN-γ and IL-2 production rate estimates (summarized in Table S23). For IFN-γ production, the median response in PWH was 78% higher than in HIV^-^ individuals, and it was also significantly more heterogenous. IR and INR individuals had median IFN-γ production rates of 0.12 d^-1^ and 0.01 d^-1^, respectively, which suggests a 91.7% slower response in INR relative to IR. For IL-2 production, HIV^-^ individuals had a median production rate of 0.12 d^-1^, which was narrowly distributed with a SD of 0.05 d^-1^. The median IL-2 production rate amongst PWH was 0.23 d^-1^, which is a 92% higher rate relative to HIV^-^, but also with a higher SD of 0.18 d^-1^. IRs display a faster IL-2 response than HIV^-^, with a median production rate of 0.27 d^-1^, vs 0.06 d^-1^ in INRs, suggesting a 77.8% slower response in the latter.

Among all vaccine doses, the lowest *T_D_* for all participants were observed post-D1, suggesting rapid population growth. For IL-2, *T_D_* were approximately equal post-D1, 45-48 days for all PWH. IFN-γ *T_D_* was 162 days for HIV^-^ individuals, while 150 and 47 days for INR and IR, respectively. The doubling time suggests that IRs mounted the IL-2 response faster than HIV^-^ and INR individuals post-D1. For D2 and D3, the estimated *T_D_* increased substantially in PWH, suggesting saturation of IFN-γ and IL-2 responses. No decline was observed in IFN-γ and IL-2 longitudinal median responses across the ∼340-day study period.

## DISCUSSION

Older individuals and PWH were prioritized for COVID-19 vaccination in Ontario among other risk groups. In the current study, we characterize how PWH aged 55 and older, including individuals with immunodeficiency, responded to three doses of COVID-19 vaccines across different arms of the immune system. We also established whether concerns over adverse reactions from HIV reservoirs to these vaccines were justified.

In clinical trials, age had effect on immunogenicity after a single dose of BNT162b2(24); however, these differences became less apparent with two doses (25). In contrast, the mRNA-1273 phase 3 studies showed significantly lower vaccine efficacy in the older age group (23), while differences between binding and neutralizing Ab responses in different age groups were also reported (43). In the ChAdOx1 phase 2/3 trial, nAb titers were relatively similar after the second dose across all age groups, but still noticeably lower in older participants aged 56-69 (21). Older age was also associated with lower T cell responses for ChAdOx1, although the efficacy against infection did not appear to be affected (21, 22).

Despite a limited number of PWH in COVID-19 vaccine trials (23, 25), some reports suggested strong initial immune responses to two vaccine doses (44-49). It should be noted, however, that these studies typically excluded the most vulnerable subgroups with low CD4^+^ T cell count (<200 cells/µL) and/or unsuppressed viremia, older people, or had small sample size. Meanwhile, there is evidence for weaker responses in PWH with lower (11, 50-52) or even preserved CD4^+^ T cell count (44). In the latter case, age discrepancies between the study group and HIV-negative participants might have played a role. Individual case studies of PWH who are not on cART, have uncontrolled viremia and low CD4^+^ T cell count, reported a failure to seroconvert after two doses of COVID-19 vaccine (53).

Only a few studies have analysed immune responses in PWH following three doses of COVID-19 vaccines, although initial reports on individuals with preserved CD4^+^ T cell count are optimistic (54). In our study of cART-treated PWH in the older age group, we showed that each COVID-19 vaccine dose, regardless of its type, induces binding IgG Abs for spike and RBD in plasma and spike IgG in saliva at levels similar in PWH and age-matched HIV-negative individuals. We found that each subsequent dose boosted IgG Abs, which peaked at four weeks after the third dose. Importantly, production and decay rates of plasma IgG did not vary as a function of HIV status, while their half-lives increased post-D3 from 28.9 to 80 days for anti-spike IgG and from 32.1 to 63 days for anti-RBD IgG, indicating a beneficial role of the booster. The two-dose half-life results compare well to previous two-dose IgG half-life measurements for BNT162b2 and mRNA-1273 vaccines which estimate a half-life of 16 days (55), and are similar to IgG half-life of 21 days for mild SARS-CoV-2 cases (56).

Unlike RBD-binding Abs, serum neutralizing responses were significantly lower in PWH than in HIV^-^ participants after two vaccine doses, although a rapid decay was observed in both groups. The observed difference could have several explanations. First, not every Ab detected in the binding assays is a nAb. Second, neutralization capacity of sera may result from nAbs to other SARS-CoV-2 spike domains, such as NTD. Most studies focus on RBD, which is the primary target for nAbs. However, NTD, which facilitates the virus’s entry into the host cell, proved to be another mutational hotspot and antigenic supersite that represents an important target for nAbs (27-31). Indeed, the moderate positive correlation that we observed between the frequency of spike-specific B cells and neutralizing titers in PWH was stronger for NTD^+^S1^+^ B cells than for RBD^+^S1^+^.

Despite the suboptimal neutralization responses in PWH with two vaccine doses, the booster seems to have ‘rescued’ them, as both PWH and HIV^-^ participants had similar NT_50_ and RBD nAb concentrations at the last study visit. These changes were mirrored by frequencies of spike-specific B cells in PWH.

We also looked at mucosal immunity of the upper respiratory tract, which is the first line of defense against SARS-CoV-2, particularly its IgA component (35, 36), and arguably the least understood in the context of COVID-19 vaccination. We demonstrated that the pattern of longitudinal changes in anti-spike IgG responses in saliva of mRNA-vaccinated participants mirrors spike/RBD IgG responses in sera. Conversely, the amplitude of IgA responses was very low, and neither D2, nor D3 had a boosting effect on them, which echoes an earlier finding by co-authors of this study on HIV-negative individuals for two doses of mRNA COVID-19 vaccine (36). Interestingly, HIV^-^ individuals showed relatively high baseline spike-specific IgA levels which remained virtually unchanged throughout the study. One possible explanation could be a cross-reactivity with Abs to seasonal coronaviruses, after repeated exposure with older age. Alternatively, these Abs may, indeed, have arisen from exposure to SARS-CoV-2 that did not result in productive infection. Both scenarios would suggest weaker and/or less durable salivary IgA responses to coronaviruses in PWH, particularly in immunological non-responders. Such poor mucosal IgA response to current intramuscular COVID-19 vaccine regimens may be affecting protection against SARS-CoV-2, especially in PWH, which indicates the need for more advanced vaccine regimens. An intranasal adenoviral booster following the intramuscular mRNA priming could be potential alternative, which has already proven to induce high levels of mucosal IgA (57).

T cell responses to COVID-19 vaccines in PWH are less understood than humoral immunity, although CD4^+^ T cell responses demonstrate a strong Th1 shift to TNF, IFN-γ and IL-2 (39), with expression patterns varying between different cells (58). A recent report suggests that most PWH generate CD4^+^ T cell responses after two vaccine doses regardless of their CD4^+^ T cell count (58). Our study demonstrates enhanced spike-specific T cell immunity in PWH who are immunological responders, even when compared to HIV^-^ participants, which suggests Th1 imprinting from pre-existing HIV infection. In immune non-responders the trend is opposite, with significantly weaker IL-2, IFN-γ and dual responses than in IRs or HIV^-^, indicating a relative defect in spike-specific Th1 immunity.

We also demonstrated a heterogeneity of cytokine production rates in PWH, which were higher for IFN-γ and IL-2 in the IR individuals than in HIV^-^, but significantly lower for IFN-γ in INR participants. Mean doubling times for both cytokines increased post-D3 by a factor of 3-4, compared to D2 estimates, suggesting the cytokine dynamics were close to saturation post-booster. No decay in the mean or median IFN-γ or IL-2 responses was observed for PWH and HIV^-^ individuals over the study period. However, the frequency values for cytokine-producing T cells were increasing significantly slower in INRs and predicted to saturate at about an order of magnitude lower median value than in IRs and HIV^-^ individuals. Our observations suggest the importance of stratification of PWH in similar studies according to their immune status. The effect of Th1 imprinting on coronavirus infections will require further study.

A significant gap in our knowledge is whether COVID-19 vaccines may destabilize HIV reservoirs, as has been the case with some other vaccines (18-20). Indeed, there has been evidence of a transient increase in HIV VL and decrease in CD4^+^ T cell count in some PWH immediately after COVID-19 vaccination, which could potentially be deleterious for individuals with immunodeficiency (44). In our study, the only significant change with respect to CD4^+^ T cell count that we observed in PWH was a transient increase post-D1. Concerning HIV VL, however, we did observe a gradual transient increase in the proportion of PWH with detectable VL post-vaccination. For most timepoints, the number of participants with ‘blips’ above 40 copies/µL (but below 200 copies/µL) did not exceed one or two, except after D2, when we registered an increase from one such case one week post-D2 to three cases at two weeks, to five at four weeks (11.6%). Four of these five participants were IRs with controlled HIV viremia. In total, we observed 24 such blips in 14 non-LLV PWH over the 48 weeks of the study post-D1. Importantly, only three of them had episodes of detectable HIV viremia during two years preceding the study, one blip per person, or three in total. In other words, the frequency of viral blips in PWH increased post-vaccination. Of note, the frequency of tests was lower before the study and varied among participants. Still, we believe, that the changes we observed in a subset of participants may not be incidental in light of the recent finding of *ex vivo* activation of HIV transcription by mRNA COVID-19 vaccines through the RIG-I/TLR–TNF–NFκb pathway (42).

Influenza vaccines have been shown to cause an increase in the HIV proviral burden in influenza-specific CD4^+^ T cells of cART-treated PWH with suppressed VL (19). In our study, we assessed changes in the size of intact HIV reservoir in peripheral CD4^+^ T cells in PWH following three doses of COVID-19 vaccines. We found no significant changes relative to the baseline after two or three doses, except in three LLV participants who had a mean increase of 93.7% in the frequency of intact HIV proviruses after the booster. The small LLV sample size may not allow a generalization, but this finding could be pointing at potential risks for PWH with unsuppressed viremia, and therefore should not be discounted. The only other study (to the best of our knowledge) that addressed this issue reported a lack of significant changes in 11 PWH after two COVID-19 vaccine doses (42). The same study reported an increase in HIV-nef-specific T-cell responses in 13 cART-treated PWH with suppressed viremia following two doses of the BNT162b2 mRNA vaccine, and a significant increase in cell-associated (but not extracellular) HIV RNA post-D3 in another cohort of eight PWH, which suggests a degree of adverse reaction to vaccines from HIV reservoirs (42). Contrary to these findings, we did not observe changes in HIV nef or gag responses after two or three vaccine doses (n=28-29). The discrepancy might be due to the nature of responses measured. In our analysis, we looked at total T cell responses, whereas the only significant difference observed by Stevenson and co-authors (42) was in the frequency of granzyme-B-producing cells. Like us, this group did not report significant changes in nef-specific IFN-γ responses. Another factor is the timing of sampling: the median of 17 and 16 days post-D1 and D2, respectively (42), as opposed to 109 and 130 days post-D2 and D3, respectively, in this study. Therefore, our study timeline may have led to us missing a potential transient increase in the frequency of nef-specific T cells. Overall, we can conclude that a limited-scale destabilization of HIV reservoirs may, indeed, occur in response to COVID-19 vaccination, which may not necessarily affect the VL or the size of HIV reservoir. These negative outcomes may evidently be more likely in unsuppressed HIV viremia.

In summary, cART-treated PWH aged 55 and older show diminished SARS-CoV-2 neutralizing responses with two COVID-19 vaccine doses which are ‘rescued’ after a booster. They also have lower levels of spike-specific IgA in saliva post-vaccination which may affect protection. Enhanced anti-spike T cell immunity in this population, with the exception of INRs, suggests Th1 imprinting from pre-existing HIV infection. Finally, COVID-19 vaccines did not affect the size of intact HIV reservoir in most PWH but were associated with an increased frequency of viral ‘blips’, and may pose potential risk in unsuppressed HIV viremia.

## ACKNOWLEDGEMENTS

This study was supported by the Juan and Stefania Speck research fund and CITF grant 253488 (MO), CTN grant CTNPT045 (VAM, MO), and OHTN grant EFP-1032-IP (CK). A-CG, KC, JG are thanking Jim Rini (University of Toronto) for supplying recombinant proteins. A-CG, KC are grateful to Yves Durocher (NRCC) for ELISA reagents. The robotics equipment at the Network Biology Collaborative Centre, Lunenfeld-Tanenbaum Research Institute, is supported by Canada Foundation for Innovation, the Ontarian Government, Genome Canada and Ontario Genomics (OGI-139). CSK and JMH acknowledge funding from NSERC, CIHR, NSERC EIDM, and the York Research Chair Program. RK acknowledges a CIHR grant MM1-174919. SM would like to acknowledge funding from Intramural Research Program of the Division of Intramural Research, NIAID, NIH. RBJ acknowledges grant UM1AI164565 from NIAID, NIH. The authors are grateful to all study participants for their time, dedication and samples provided.

## AUTHOR CONTRIBUTIONS

Study design and conceptualization: VAM, EB, YC, RR, RK, TWC, CK, MO. Funding acquisition: VAM, CK, MO. Project administration and/or data curation: VAM, EZM, EB, GC, SC, AI, CK. Methodology: VAM, GC, YC, RR, RK, JMH, SM, JS, JG, A-CG, TWC, CK, MO. Sample collection and/or processing: EB, EZM, SG, PS, SC, AI. Experimentation: VAM, EZM, SG, PS, HS, RL, PB, LW, SSM, GM, MD-B, TT, LK, FQ, AP. Data analysis and/or review: VAM, KC, JHW, FQ, AP, DCCJr, RBJ, JMH, SM, JS, JG, A-CG. Statistical analysis: TL, CSK. Mathematical modelling: CSK. Supervision: VAM, KC, JHW, JMH, SM, JS, JG, A-CG. Drafting the manuscript: VAM, CSK, TL. Critical revision of the manuscript: MO, KC, JMH, SM, JS, RR, JG, A-CG.

## MATERIALS AND METHODS

### Study design

This was a single-site prospective non-randomized observational study (CTNPT 045) where PWH and HIV-negative individuals were followed over 48 weeks with blood draws and saliva sampling, with evaluations taken at screening, baseline, and 12 additional timepoints (Fig. 1) preceding or following three doses of one (or a combined regimen) of the following COVID-19 vaccines: an adenoviral vector vaccine ChAdOx1 (Oxford University, AstraZeneca) and two mRNA vaccines, BNT162b2 (Pfizer-BioNTech) and mRNA1273 (Moderna).

### Participants

A total of 91 participants aged 55 and over were recruited for the study through Maple Leaf Medical Clinic in Toronto, Canada, including 68 PWH and 23 HIV-negative individuals (Fig. 1A-B, Supplemental Table S1). We initially approached 73 eligible PWH, five of whom declined to participate. Next, we approached 28 HIV-negative individuals in the matching age range. Five of them decline to participate. In total, 91 out of 101 approached candidates (90.1%) volunteered and signed an informed consent form. Due to the rapid vaccine rollout in Ontario for the older demographic, baseline samples could only be obtained from 57 participants, including 45 PWH and 12 HIV-negative individuals. An additional 34 participants joined the study at Visit 8, including 23 PWH and 11 HIV^-^. All PWH had been on cART for at least one year. For PWH subgroup definitions, see SM2.

### Study visits and sampling

Vaccination was not provided as a part of this study. The timeline of the study protocol is shown on Fig. 1C and detailed in SM1. At the screening visit, we performed routine blood work (hematology, biochemistry) for all participants and HIV test for HIV-negative individuals. PWH were assayed for CD4^+^ T-cell count, VL, and CD4^+^/CD8^+^ ratio, which were also recorded retrospectively, along with CD4^+^ T-cell nadir. CD4^+^ and CD8^+^ T-cell count and percentage, CD4^+^/CD8^+^ ratio and VL were assessed at each study visit. For each participant, blood was collected in a single SST tube (8.5 mL/tube), and in up to 10 ACD tubes (at V1, V4a, and V8-9; 8.5 mL/tube). Saliva was collected in a Salivette (Sarstedt, Germany) at V1, V4-5, and V8-9. Specimens were processed immediately upon sampling.

### Serum isolation

SST tubes with whole clotted blood were centrifuged at 1,200×*g* for 10 minutes. Sera were then aspirated, aliquoted in 0.5 mL increments and frozen at –80°C.

### PBMC isolation

PBMC were isolated by centrifugation using Ficoll-Paque PLUS (GE Healthcare) as previously described (59). The cell suspension was diluted 1:1 with a freezing medium and aliquoted for storage at –150°C.

### Saliva processing

Salivette tubes were centrifuged at 1,200×*g* for 10 minutes at room temperature (RT). Saliva was collected from the bottom section, aliquoted in cryovials up to 1 mL per vial, and frozen at –80°C.

### Antibody detection in serum (ELISA)

An automated ELISA assay was used to detect total IgG Ab levels to full-length spike trimer, RBD and nucleocapsid as previously described (33, 38). Briefly, 384-well microplates were pre-coated with spike (SmT1), RBD or N, supplied by the National Research Council of Canada (NRC). The main steps were blocking in Blocker BLOTTO (ThermoFisher Scientific), incubation with 1:160 or 1:2,560 sera dilutions, incubation with human anti-IgG fused to HRP (IgG#5 by NRC), and incubation with ELISA Pico Chemiluminescent Substrate (ThermoFisher Scientific). Chemiluminescence was read on an EnVision 2105 Multimode Plate Reader (Perkin Elmer). Raw values were normalized to a synthetic standard on each plate (VHH72-Fc by NRC for spike/RBD or an anti-N IgG from Genscript, #A02039). The relative ratios were further converted to BAU/mL using the WHO International Standard 20/136 as a calibrant (33, 60). Seropositivity thresholds were determined for the 1:160 dilution using 3×SD from the mean of controls (33).

### Live SARS-CoV-2 microneutralization (MN)

A MN assay to measure 50% neutralizing titers (NT_50_) in the sera was performed as described previously (61). Briefly, heat-inactivated sera were serially diluted and incubated with 100 TCID_50_ of the wild type SARS-CoV-2 virus in serum-free DMEM, then added onto the VeroE6 cells in 96-well plates. After 1h, inoculums were removed and DMEM with 2% FBS was added to the cells. The cells were incubated for 5 days and checked daily for cytopathic effects (CPE). NT_50_ titer was defined as the highest dilution factor of the serum that protected 50% of cells from CPEs and calculated by the four-parameter logistic regression in GraphPad Prism 8.

### Frequency of spike-specific B cells (spectral flow cytometry)

We measured frequencies of RBD- and NTD-specific B cells (RBD^+^S1^+^ and NTD^+^S1^+^, respectively) among total B cells by spectral flow cytometry using a 19-color subset (Table S24) of a previously described 21-color panel (62). Briefly, the biotinylated spike proteins were tetramerized with fluorescently labeled streptavidin at a molar ratio of 4:1 as described previously (63), with minor modifications (62). Cryopreserved PBMCs were thawed and stained with Zombie NIR Fixable Viability Dye, followed by staining with a cocktail of mAbs and four fluorochrome-conjugated spike proteins in staining buffer (2% FBS/PBS) supplemented with Brilliant Stain Buffer Plus (BD Biosciences). The stained cells were fixed (Lysing Solution, BD Biosciences) and acquired on an Aurora spectral cytometer using SpectroFlo Software v3.0.1 (Cytek Biosciences). The analysis was performed with FlowJo v10 (BD Biosciences). The CIRC0041 baseline plot is shown as an example of our gating strategy (Fig. S9).

### ACE2 displacement assay (snELISA)

The rACE2 displacement assay was performed as previously described (33, 34), except assay plates were Maxisorp (ThermoFisher Scientific). Briefly, the 384-well plates were coated with RBD (supplied by NRC), blocked with BSA in PBS-T, incubated with 1:10 or 1:40 sera dilutions, then with biotinylated ACE2 (supplied by Jim Rini, University of Toronto), and treated with Streptavidin-Peroxidase Polymer Ultrasensitive (Sigma-Aldrich). Incubation with ELISA Pico Chemiluminescent Substrate and reading on the EnVision were performed as in direct detection. All values were normalized to sample-free blanks on the same plate. The resulting relative ratios (RR) were converted to IU/mL using the WHO International Standard 20/136 as a calibrant (33) and the following formula: log_2_(IU/mL at sample dilution d)=RR/(-0.2308)+5.7898+log_2_(d). Neutralization threshold was determined for the 1:10 dilution as 2×MAD (median absolute deviation) from the mean of blanks.

### Antibody detection in saliva (ELISA)

ELISA assays for anti-spike IgA and IgG in saliva were performed in 96-well plates pre-coated with streptavidin as described earlier (36, 38). Briefly, the plates were coated with biotinylated SARS-CoV-2 spike or PBS (controls) and incubated with saliva samples diluted with a BLOTTO solution (BioShop) at 1:5, 1:10, 1:20. After washing, the plates were incubated with HRP-conjugated goat anti-human IgG and IgA secondary Abs (Southern Biotech, IgG: 2044-05, IgA: 2053-05) and developed by the TMB substrate (ThermoFisher). The reaction was stopped with 1N H_2_SO_4_, and the plates were read on the Thermo Multiskan FC spectrophotometer. Raw OD450 values for the PBS controls were subtracted from the sample values for each dilution. Each adjusted value was used to calculate the area under the curve (AUC) which was then normalized to the AUC of the positive control of pooled saliva from COVID-19 acute and convalescent participants. The normalized AUC was also converted to %AUC of the positive control. Pooled early pandemic saliva from 10 individuals with a median bin age range of 30-39 years was used as a negative control whose integrated scores were calculated as described previously (36). Seropositivity values for each isotype (0.35% for IgG, 4.06% for IgA) were calculated as average integrated scores of pre-pandemic samples (0.086% for IgG, 0.85% for IgA) plus 2×SD. Only samples from mRNA-vaccinated individuals were used (n=31).

### T cell cytokine responses (dual-color ELISpot)

*Ex vivo* ELISpot assay was performed as previously described (59). Briefly, 96-well MultiScreen plates (Millipore) were coated with capture mAbs to IFN-γ (Mabtech #1-D1K) and IL-2 (Mabtech #MT2A91/2C95) and blocked with R-10 medium. PBMCs were plated at 200,000 cells/well and stimulated with one of the following peptide pools: SARS-CoV-2 spike (JPT #PM-WCPV-S-1), HIV-1 nef or HIV-1 gag (NIH HIV Reagent Program #ARP-12822, #ARP-12437). Positive controls included SEB (Sigma-Aldrich), CEF (NIH HIV Reagent Program), and CEFTA (Mabtech). Washed plates were incubated with alkaline-phosphatase-conjugated mAb for IFN-γ (Mabtech #7-B6-1) and with biotinylated mAb for IL-2 (Mabtech #MT8G10), then incubated with streptavidin-HRP and developed with Vector Blue for ALP and Vector NovaRed for HRP (Vector Laboratories). The mean number of spots in the negative-control (DMSO) wells were subtracted from peptide-stimulated wells, and the results were expressed as spot-forming cells (SFC) per 10^6^ PBMC.

### Intact Proviral DNA Assay (IPDA)

The assay was run in 96-well plates (Bio-Rad) as described previously (41, 64), with minor adjustments to a 22-µL reaction volume and four technical replicates (wells) per HIV and RPP30 reaction per sample. CD4^+^ T cells were isolated from PBMCs with the EasySep Human CD4^+^ T cell Enrichment Kit (STEMCELL Technologies). Genomic DNA was isolated from a median of 3.65 million CD4^+^ T cells (IQR 2.48-4.98) with the DNeasy Blood & Tissue Kit (QIAGEN). Each HIV reaction contained 825 ng of gDNA, and each RPP30 reaction contained 8.25 ng of gDNA. Genomic DNA was combined with ddPCR Supermix for Probes (no dUTPs, Bio-Rad), primers (900 nM final, IDT), probes (250 nM final, ThermoFisher Scientific), and XhoI restriction enzyme (0.5 U/µL, ThermoFisher Scientific). For sequences, see Table S25. For participants OM5365, CIRC0324, OM5005, and OM5016 we used an alternative primer/probe mix (64) as the IPDA Ψ failed (Alt Ψ in Table S25). Droplets were generated with Automated Droplet Generator (Bio-Rad), and the PCR was run as described earlier (41, 64). The plates were read on a QX200 Droplet Reader (Bio-Rad) and analysed with QuantaSoft v.1.7.4.0917 and QuantaSoft Analysis Pro v.1.0.596 (Bio-Rad). Replicate wells were merged prior to analysis. DNA from JLat-8.4 cells, which harbour a single HIV provirus per cell, was used as a positive control. HIV^-^ DNA from Huh-7 cells and no-template controls were also used.

### Vaccination model for within-host immunization

To model the vaccine inoculation and subsequent within-host immunization, we used a simplified model previously used to model immunological outcomes from mRNA-based vaccines (55), determine significant immunological timescales from mRNA-based vaccines (65), and explore differing prime-booster strategies of AstraZeneca vaccines in humans (66). All model terms, parameter assumptions, and details of sensitivity analysis, are described in SM3-4. A schematic of the model is shown in Fig. 9. The model equations are provided in Equations S1a-S1i.

**Figure 9.**
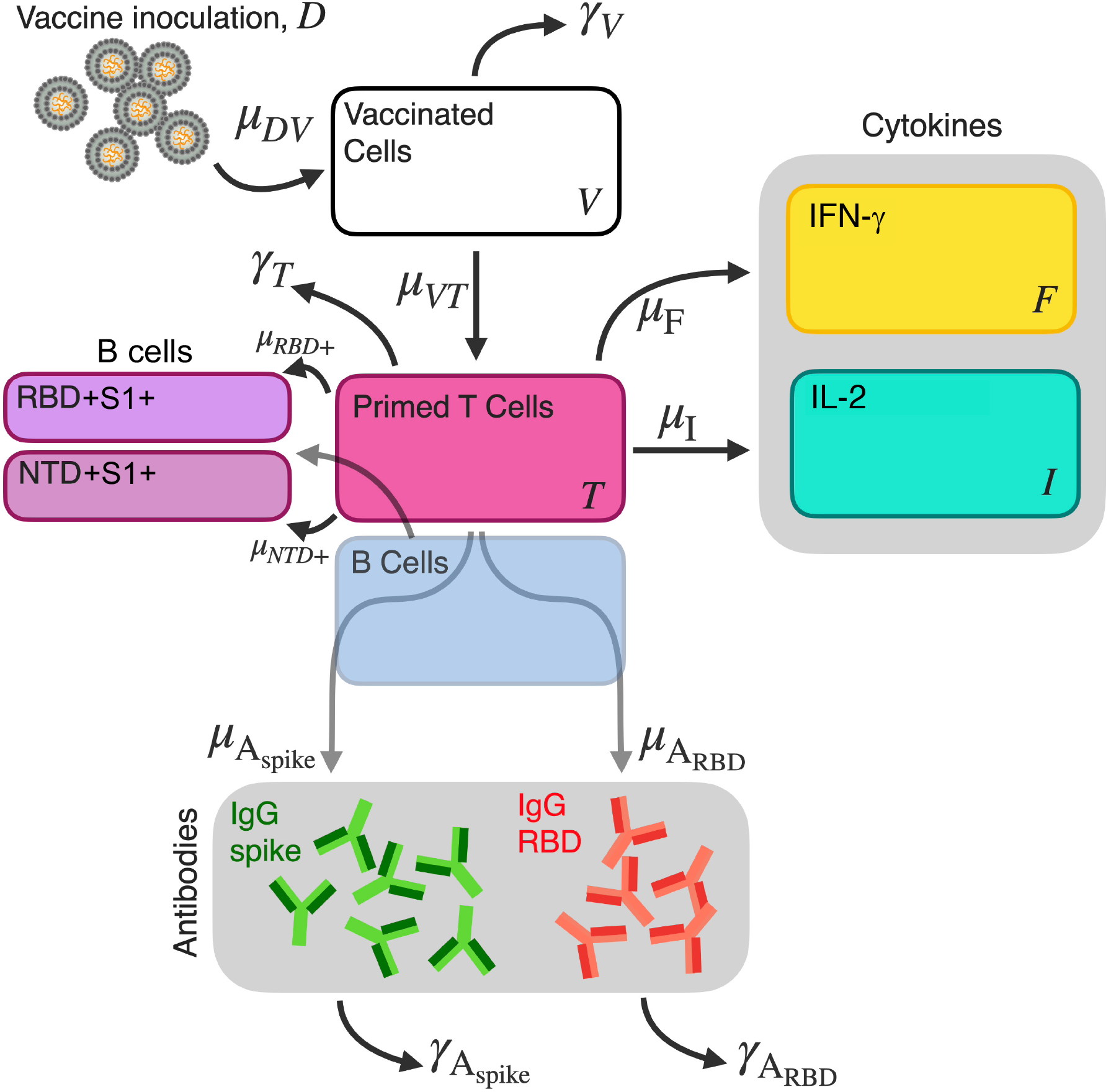
Vaccination model for within-host immunization. A schematic diagram of COVID-19 vaccine inoculation, T cell stimulation, cytokine, IgG, and spike-specific B cell production model (Eq. 1). For a detailed description of the model, see SM3.

### Statistical analysis

Comparison of serum IgG levels was done by mixed effects linear regression adjusted for the same set of variables. Serum IgG levels were not normally distributed and were log*_e_*- transformed for analysis. Baseline live SARS-CoV-2 neutralization titers were summarized as dichotomized variable given that most values were zero. Comparison between groups was based on Chi-square test or Fisher’s exact test as appropriate. Post-baseline comparison between groups was performed with quantile regression adjusted for age, vaccine regimen for D1 and D2 (mRNA/mRNA, ChAdOx1/ChAdOx1 and ChAdOx1/mRNA), and time between D1 and D2. For comparison at V9, we further adjusted for time between D3 and V9. For saliva ELISA and T cell ELISpot, the sample sizes were too small for a proper adjusted comparison between groups. Unadjusted comparison was based on Wilcoxon rank sum test. Analyses were conducted using SAS 9.4 (SAS Institute Inc., Cary, NC).

### Study approval

All study participants provided informed written consent. The study protocol and consent form were approved by the University of Toronto Research Ethics Board (RIS #40713) and Sinai Health REB (21-0223-E). The study was conducted in accordance with the protocol, applicable regulations and guidelines for Good Clinical Practice (GCP), Health Canada’s regulations, and the Tri-Council Policy Statement: Ethical Conduct for Research Involving Humans (TCPS 2.0).

