## Supplemental Data for "Immunogenicity of COVID-19 vaccines and their effect on the HIV reservoir in older people with HIV"

**SUPPLEMENTAL METHODS**

**SM1. Study timeline**

The screening visit took place within 90 days prior to the baseline visit. The baseline visit (V1) occurred up to 24 hours prior to the first COVID-19 vaccine dose (D1). The second, third and fourth study visits (V2, V3, V4) are 10, 20 and 28 days following D1, respectively. As extended time intervals between the first and the second (D2) vaccine doses had been adopted in Ontario, we introduced additional study visit up to three days prior to D2 – Visit 4a (V4a). Visits 5, 6 and 7 (V5, V6, V7) took place one, two and four weeks after D2, respectively. Our primary endpoint, Visit 8 (V8), occurred at a 24-week mark (six months) following D1. Visit 8a (V8a) took place within three days preceding the third vaccine dose (D3), which was then followed by Visits 8b and 8c (V8b, V8c) at four and 16 weeks post-D3. The final study visit, Visit 9 (V9), corresponds to 48 weeks (one year) after D1.

Sixty out of 91 participants had either completed V8a, or it was not scheduled for them because their V8 fell within seven days preceding D3. For most participants in the additional cohort recruited at V8, the V8a and V8b visits were not scheduled. For those participants for whom V8c and V9 fell within two weeks of each other, these two timepoints were combined into one study visit (Table S2). We allowed a three-day window for each study visit on each side of the target date. For the datasets appearing as ‘Out-of-window data excluded’, we excluded samples outside of the three-day window from the target date (±7 days for V8 and V9). Samples deviating by more than 14 days from the target date were excluded from all analyses. Three participants received a fourth dose before the final study visit, so their V9 samples were excluded from the datasets.

**SM2. PWH subgroup definitions**

Immunological responders (IR): undetectable viral load (VL) for at least one year, CD4^+^ T-cell count > 500 cells/µL and any CD4^+^/CD8^+^ ratio, or CD4^+^ T-cell count = 350-500 cells/µL and CD4^+^/CD8^+^ ratio ≥ 0.7. Immunological non-responders (INR): undetectable VL for at least one year, CD4^+^ T-cell count = 350-500 cells/µL and CD4^+^/CD8^+^ ratio < 0.7, or CD4^+^ T-cell count < 350 cells/µL and any CD4^+^/CD8^+^ ratio. PWH with low-level viremia (LLV): cART-adherent, VL > 40 copies/mL at least three times a year for at least three consecutive years, any CD4^+^ T-cell count, any CD4^+^/CD8^+^ ratio. Long-term non-progressors (LTNP): no decline for at least five years without cART, VL < 5,000 copies/mL (our participant had low or undetectable VL from 1987 – when he was diagnosed with HIV, to 2011 – when he elected to start cART), CD4^+^ T cell count > 500 cells/μL.

**SM3. Vaccination model for within-host immunization**

To model the vaccine inoculation and subsequent within-host immunization, we used a simplified compartmental model previously shown to accurately determine long-term IgG decay and capture cytokine dynamics of BNT162b2 and mRNA-1273^1^, determine significant immunological timescales from mRNA-LNP-based vaccines^2^, and explore differing prime-booster strategies of AstraZeneca vaccines in humans^3^. The model is briefly described as follows. The initial concentration of the vaccine (BNT162b2, mRNA-1273, or AstraZeneca) in the host following the inoculation is *D_0_*. Upon inoculation, the host begins to produce vaccinated cells, *V*, through the interaction term *μ_VD_*. The amount of vaccine concentration, *D*, is strictly decreasing within the host for all *t* > *t_0_*. Vaccinated cells produce antigen, and are not modelled as cell-type specific. The priming dynamics of naïve T cells into antigen-primed T cells is captured by the term *μ_TV_*, loss of primed T-cell secretion activity is *γ_T_*. Increase in IFN-γ and IL-2 cytokine-producing T cells is modelled by the proliferation terms *μ_F_* and *μ_I_*, respectively. Plasma B cells are primed by antigen-specific T cells^4^. However, in this work longitudinal B cell trajectories are absent. In the absence of plasma B cell longitudinal data, we model the production of IgG values as linearly proportional to primed antigen-specific T cells and thus implicitly consider the plasma B cell response. Generation of memory B cells is linked to a T-dependent response. We further consider the development of anti-RBD (RBD^+^S1^+^) and anti-NTD (NTD^+^S1^+^) memory B cells through proliferation terms *μ_RBD+_* and *μ_NTD+_*, respectively. The full model used in this work, shown in Eq. 1, is described by nine equations, whereby Eqs. S1d-i are fit to longitudinal IgG spike, IgG RBD, memory B cell RBD^+^ and NTD^+^, IFN-γ, and IL-2 data sets, respectively. A schematic of the model is shown in Fig. 9.

Equation 1:

Vaccine inoculation:	$\frac{dD}{dt}$ *= –μ_VD_ D*	(S1a)

Vaccinated cells:	$\frac{dV}{dt}$ *=* *μ_VD_ D – γ_V_ V*	(S1b)

Primed T cells:	$\frac{dT}{dt}$ *= μ_VT_ V – γ_T_ T*	(S1c)

IgG spike:	$\frac{dAspike}{dt}$ *= μ_Aspike_ T – γ_Aspike_ A_spike_*	(S1d)

IgG RBD:	$\frac{dARBD}{dt}$ *= μ_ARBD_ T – γ_ARBD_ A_RBD_*	(S1e)

RBD^+^ memory B cell response:	$\frac{dMRBD+}{dt}$ *= μ_RBD+_ T*	(S1f)

NTD^+^ memory B cell response:	$\frac{dMNTD+}{dt}$ *= μ_NTD+_ T*	(S1g)

IFN-γ:	$\frac{dF}{dt}$ *= μ_F_ T*	(S1h)

IL-2:	$\frac{dI}{dt}$ *= μ_I_ T*	(S1i)

**SM4. Parameter assumptions, estimations, initial conditions, and sensitivity analysis for the model**

Previous findings in mice have shown that mRNA-LNP vaccines are translated locally at an intramuscular injection site for approximately 10 days^1^. In this work we, therefore, fix the translation half-life to approximately 10 days upon receiving a vaccine dose, leading to a fixed γ_v_ of 0.07 d^–1^. Initial inoculation (Dose 1, D1), and subsequent dosages (Dose 2 and 3, D2-3) are assumed to be of value D_0_=2 and are in arbitrary units of concentration. Thus, we do not consider differences in dosage concentrations between vaccine types.

All fits to clinical data using Eq. 1 were performed in Monolix v2020R1 using non-linear mixed effects. All individuals and all individual data points for three doses are used in the longitudinal modelling assay; data points acquiring post-Dose-4 are removed from the analysis. IgG best-fit parameters are obtained in a dose-dependent two-stage fitting process whereby the data is separated into two parts: baseline to pre-Dose-3, and pre-Dose-3 through to end of study timepoint V9. A set of IgG fit parameters are therefore obtained for each of the two fits. Cytokine data is included in the IgG fits. However, these are not reported on as the errors and uncertainties in the cytokine-respective two-stage fitted parameters were unable to be estimated due to the lack of data resolution within the time window for the two-stage fits. We note that the production and decay rates estimated from the longitudinal modelling approach will be affected by the IgG saturation values which limit the dynamical range in observed BAU/mL. Therefore, our production and decay rates likely underestimate and overestimate the true value, respectively.

A single-stage fitting process is used to estimate robust longitudinal cytokine and memory B cell dynamics: these fitted parameters are estimated through a fit to all Dose-3 clinical data present for all time points, V1 through to V9, resulting in a single set of cytokine model parameters. The cytokine and memory B cell data resolution do not support dose-dependent parameter estimate as done with the IgG. All random effects, and relative standard error on random effects from the model results are shown in Table S26.

We performed structural and practical sensitivity analysis to characterize the response of our model outputs to variation in the fitted parameters^5^. Latin hypercube sampling (LHC) and partial rank correlation coefficient (PRCC)^6^ are employed to study the structural identifiability of the longitudinal model outcomes. For a particular model parameter, PRCC values close to the maximum value of 1.0 indicate model output is highly sensitive to variation in that parameter, with values greater than 0.5 considered significant^7^. LHC PRCC analysis results are shown in Fig. S10. We also report random effects and relative standard error on random effects to estimate practical data-driven identifiability in model outcomes (Table S26). Practical identifiability analysis quantifies how well-constrained the estimated model parameters are by the data sets to which they are being fit^5^. Both structural and practical demonstrate no identifiability issues with any parameters reported in this work for IgG, memory B cells, or cytokines.

**Supplemental table S1**. Demographics, HIV status and COVID-19 vaccination profiles of study participants.

| **#** | **Participant**  **ID** | **Age** | **Sex** | **Race^a^** | **HIV Status, Subgroup^b^** | **HIV**  **since** | **Years VL^c^ suppressed** | **Type of COVID-19 Vaccine^d^** | | | **Joined at**  **Baseline vs. V8** |
| --- | --- | --- | --- | --- | --- | --- | --- | --- | --- | --- | --- |
|  |  |  |  |  |  |  |  | **D1^e^** | **D2** | **D3** |  |
| 1 | OM 5402 | 59 | M | C | N | NA^f^ | NA | AZO | AZO | PB | Baseline |
| 2 | OM 5399 | 77 | M | C | N | NA | NA | AZO | AZO | PB | Baseline |
| 3 | OM 5405 | 69 | M | C | N | NA | NA | PB | PB | PB | Baseline |
| 4 | OM 5398 | 71 | M | C | N | NA | NA | PB | PB | PB | Baseline |
| 5 | OM 5164 | 60 | M | C | N | NA | NA | AZO | AZO | PB | Baseline |
| 6 | OM 5382 | 57 | M | C | N | NA | NA | AZO | AZO | PB | Baseline |
| 7 | OM 5406 | 68 | M | C | N | NA | NA | PB | PB | PB | Baseline |
| 8 | OM 5401 | 65 | M | C | N | NA | NA | PB | PB | PB | Baseline |
| 9 | OM 5397 | 73 | M | C | N | NA | NA | AZO | AZO | PB | Baseline |
| 10 | OM 5033 | 55 | M | C | N | NA | NA | PB | PB | PB | Baseline |
| 11 | OM 5408 | 58 | M | C | N | NA | NA | PB | PB | PB | Baseline |
| 12 | OM 5370 | 59 | M | C | N | NA | NA | PB | M | PB | Baseline |
| 13 | OM 5413 | 62 | M | C | N | NA | NA | AZO | AZO | PB | V8 |
| 14 | OM 5410 | 73 | M | C | N | NA | NA | AZO | AZO | PB | V8 |
| 15 | OM 5414 | 70 | M | C | N | NA | NA | PB | PB | PB | V8 |
| 16 | OM 5415 | 70 | M | EA | N | NA | NA | PB | PB | PB | V8 |
| 17 | OM 5411 | 63 | M | C | N | NA | NA | AZO | AZO | M | V8 |
| 18 | OM 5416 | 57 | M | C | N | NA | NA | PB | PB | PB | V8 |
| 19 | OM 5403 | 67 | M | C | N | NA | NA | PB | PB | PB | V8 |
| 20 | OM 5064 | 55 | M | C | N | NA | NA | PB | PB | PB | V8 |
| 21 | OM 5412 | 62 | M | B | N | NA | NA | M | M | M | V8 |
| 22 | OM 5392 | 58 | M | C | N | NA | NA | AZO | M | M | V8 |
| 23 | OM 5417 | 62 | M | C | N | NA | NA | AZO | AZO | M | V8 |
| 24 | CIRC 0054 | 76 | M | C | LTNP | 1998 | 24 | M | M | M | Baseline |
| 25 | OM 215 | 62 | M | C | IR | 1993 | 28 | M | M | PB | Baseline |
| 26 | OM 5085 | 64 | M | C | IR | 1985 | 20 | AZO | AZO | PB | Baseline |
| 27 | OM 5076 | 64 | M | L | IR | 1989 | 21 | AZO | AZO | PB | Baseline |
| 28 | CIRC 0022 | 79 | M | C | IR | 1995 | 23 | PB | PB | PB | Baseline |
| 29 | OM 5056 | 66 | M | C | IR | 1987 | 24 | AZO | AZO | PB | Baseline |
| 30 | CIRC 0116 | 65 | M | C | IR | 2000 | 19 | PB | PB | PB | Baseline |
| 31 | OM 5030 | 62 | M | L | IR | 1985 | 20 | AZO | AZO | PB | Baseline |
| 32 | CIRC 0050 | 68 | M | C | IR | 1989 | 23 | AZO | AZO | M | Baseline |
| 33 | OM 5135 | 62 | M | C | IR | 1988 | 17 | AZO | AZO | M | Baseline |
| 34 | OM 5208 | 65 | M | C | IR | 1989 | 17 | AZO | AZO | PB | Baseline |
| 35 | CIRC 0120 | 78 | M | C | IR | 1989 | 16 | PB | M | M | Baseline |
| 36 | CIRC 0313 | 72 | M | C | IR | 1984 | 20 | AZO | AZO | PB | Baseline |
| 37 | OM 5051 | 74 | M | C | IR | 1990 | 23 | PB | M | M | Baseline |
| 38 | CIRC 0041 | 72 | M | C | IR | 1999 | 22 | PB | PB | PB | Baseline |
| 39 | OM 5202 | 69 | M | C | IR | 2008 | 12 | PB | PB | PB | Baseline |
| 40 | OM 5055 | 55 | M | C | IR | 2002 | 25 | M | M | M | Baseline |
| 41 | OM 5013 | 56 | M | C | IR | 1989 | 25 | AZO | AZO | PB | Baseline |
| 42 | CIRC 0066 | 59 | M | C | IR | 1989 | 24 | AZO | AZO | PB | Baseline |
| 43 | OM 5128 | 55 | M | C | IR | 2002 | 17 | PB | PB | ND^6^ | Baseline |
| 44 | OM 5130 | 56 | M | C | IR | 1990 | 16 | PB | M | M | Baseline |
| 45 | OM 5265 | 57 | M | ME | IR | 2011 | 10 | AZO | AZO | PB | Baseline |
| 46 | OM 5232 | 58 | M | C | IR | 2000 | 21 | PB | PB | PB | Baseline |
| 47 | OM 5368 | 57 | M | C | IR | 2015 | 5 | AZO | AZO | PB | Baseline |
| 48 | OM 5407 | 59 | M | SA | IR | 1992 | 7 | AZO | AZO | PB | Baseline |
| 49 | OM 5248 | 55 | M | C | IR | 1998 | 20 | AZO | AZO | PB | Baseline |
| 50 | OM 5365 | 61 | M | C | IR | 1990 | 18 | PB | PB | PB | Baseline |
| 51 | OM 5225 | 68 | M | C | IR | 1999 | 21 | PB | PB | PB | Baseline |
| 52 | OM 5200 | 72 | M | C | IR | 1990 | 24 | PB | PB | PB | Baseline |
| 53 | CIRC 0280 | 70 | F | C | IR | 1992 | 11 | PB | PB | PB | Baseline |
| 54 | CIRC 0196 | 58 | M | C | IR | 2007 | 8 | AZO | AZO | PB | V8 |
| 55 | CIRC 0281 | 60 | M | L | IR | 1997 | 13 | PB | PB | PB | V8 |
| 56 | CIRC 0146 | 77 | M | C | IR | 1990 | 15 | PB | PB | PB | V8 |
| 57 | CIRC 0113 | 55 | M | C | IR | 1993 | 19 | AZO | AZO | MB | V8 |
| 58 | CIRC 0323 | 58 | M | C | IR | 1990 | 19 | AZO | M | PB | V8 |
| 59 | OM 5213 | 65 | M | IB | IR | 1993 | 20 | PB | PB | PB | V8 |
| 60 | CIRC 0273 | 85 | M | C | IR | 2000 | 16 | PB | M | PB | V8 |
| 61 | CIRC 0164 | 58 | M | C | IR | 2006 | 11 | AZO | AZO | M | V8 |
| 62 | OM 5122 | 54 | M | C | IR | 2007 | 13 | AZO | M | M | V8 |
| 63 | OM 5409 | 55 | M | ME | IR | 2007 | 12 | AZO | M | M | V8 |
| 64 | CIRC 0028 | 62 | M | C | IR | 1991 | 30 | PB | M | M | V8 |
| 65 | CIRC 0188 | 66 | M | C | IR | 1996 | 20 | PB | M | PB | V8 |
| 66 | CIRC 0302 | 55 | M | C | IR | 2000 | 17 | AZO | M | M | V8 |
| 67 | CIRC 0324 | 63 | M | C | INR | 1987 | 16 | PB | PB | PB | Baseline |
| 68 | OM 5400 | 63 | M | C | INR | 1989 | 15 | AZO | AZO | PB | Baseline |
| 69 | OM 5005 | 63 | M | C | INR | 1987 | 15 | AZO | AZO | PB | Baseline |
| 70 | CIRC 0216 | 62 | M | L | INR | 2000 | 17 | AZO | AZO | PB | Baseline |
| 71 | OM 5016 | 62 | M | C | INR | 1992 | 14 | AZO | AZO | PB | Baseline |
| 72 | CIRC 0322 | 71 | M | C | INR | 1990 | 14 | PB | PB | PB | Baseline |
| 73 | CIRC 0266 | 66 | M | C | INR | 1988 | 16 | PB | PB | PB | Baseline |
| 74 | CIRC 0060 | 63 | M | C | INR | 1994 | 24 | PB | PB | PB | Baseline |
| 75 | OM 5168 | 59 | M | C | INR | 1985 | 20 | PB | PB | PB | Baseline |
| 76 | OM 5156 | 55 | M | C | INR | 1985 | 14 | AZO | AZO | PB | Baseline |
| 77 | CIRC 0270 | 66 | M | C | INR | 1985 | 22 | PB | PB | PB | Baseline |
| 78 | CIRC 0274 | 56 | M | B | INR | 2000 | 20 | PB | M | PB | Baseline |
| 79 | OM 5131 | 62 | M | C | INR | 1982 | 15 | AZO | M | PB | V8 |
| 80 | CIRC 0319 | 78 | M | C | INR | 1990 | 30 | AZO | AZO | PB | V8 |
| 81 | OM 5244 | 72 | M | C | INR | 1986 | 20 | M | M | PB | V8 |
| 82 | CIRC 0058 | 70 | M | C | INR | 1993 | 23 | AZO | PB | PB | V8 |
| 83 | OM 5226 | 58 | M | C | INR | 2005 | 16 | AZO | AZO | PB | V8 |
| 84 | CIRC 0174 | 66 | M | C | INR | 1994 | 20 | PB | PB | PB | V8 |
| 85 | CIRC 0036 | 69 | M | C | INR | 1991 | 30 | PB | PB | PB | V8 |
| 86 | CIRC 0308 | 57 | M | C | INR | 1988 | 19 | PB | PB | PB | V8 |
| 87 | OM 5094 | 73 | M | C | LLV | 1985 | NA | PB | PB | PB | Baseline |
| 88 | OM 5019 | 63 | M | C | LLV | 2005 | NA | AZO | AZO | PB | Baseline |
| 89 | OM 5004 | 66 | M | SA | LLV | 1981 | NA | AZO | AZO | PB | Baseline |
| 90 | CIRC 0106 | 70 | M | С | LLV | 1990 | NA | PB | PB | PB | V8 |
| 91 | OM 5211 | 62 | M | C | LLV | 2000 | NA | AZO | AZO | PB | V8 |

^a^C – Caucasian, B – black, L – Latino, ME – Middle Eastern, EA – East Asian, SA – South Asian, IB – Indigenous Brazilian.

^b^N – HIV-negative, IR – PWH who are immunological responders, INR – PWH who are immunological non-responders, LLV – PWH with low-level viremia, LTNP – PWH who are long-term non-progressors.

^c^VL – viral load.

^d^AZO – ChAdOx1 adenoviral vector vaccine (AstraZeneca, Oxford University), PB – BNT162b2 mRNA vaccine (Pfizer, BioNTech), M – mRNA-1273 mRNA vaccine (Moderna), ND – not determined.

^e^D1-D3 – COVID-19 vaccine doses 1-3.

^f^NA – not applicable.

**Supplemental Table S2.** Study progress and adherence to the protocol among participants.

| Visit # | V1 | V2 | V3 | V4 | V4a^a^ | V5 | V6 | V7 | V8^b^ | V8a^c^ | V8b^d^ | V8c^e^ | V9 |
| --- | --- | --- | --- | --- | --- | --- | --- | --- | --- | --- | --- | --- | --- |
| Participants  completed | 57/57 | 56/57 | 56/57 | 56/57 | 50/57 | 52/57 | 55/57 | 55/57 | 91/91 | 60/91 | 67/91 | 82/91 | 89/91 |

^a^For V4a, two participants failed to complete, for five more V4a was not scheduled because D2 was administered without a delay.

^b^Additional 34 participants were recruited at this point, increasing the total number from 57 to 91.

^c^60 out of 91 participants have either completed V8a or it was not scheduled because V8 fell within 7 days preceding D3. For most participants in the additional cohort recruited at V8, the V8a visit was not scheduled.

^d^For most participants in the additional cohort recruited at V8, the V8b visit was not scheduled.

^e^82 out of 91 participants have either completed V8c, or it was not scheduled because it fell within 14 days of V9.

**Supplemental table S3A**. RBD IgG levels in serum (BAU/mL) in PWH and HIV^–^ participants. N^+^/post-COVID-19 samples (n=16) and V9 samples preceded by D4 (n=3) were excluded.

|  | **HIV–** | **HIV+** | **IRs** | **INRs** | **LLVs** | **LTNPs** |
| --- | --- | --- | --- | --- | --- | --- |
| **V1** |  |  |  |  |  |  |
| n | 11 | 44 | 28 | 12 | 3 | 1 |
| Median (IQR) | 3.02^a^ (3.02, 3.11) | 3.02 (3.02, 3.70) | 3.02 (3.02, 4.03) | 3.02 (3.02, 3.57) | 3.02 (3.02, 3.02) | 3.02 (3.02, 3.02) |
| Range | (3.02, 10.1) | (3.02, 779) | (3.02, 779) | (3.02, 4.44) | (3.02, 3.02) | (3.02, 3.02) |
| **V4** |  |  |  |  |  |  |
| n | 11 | 43 | 28 | 11 | 3 | 1 |
| Median (IQR) | 64.5 (23.4, 150) | 45.6 (15.4, 120) | 76.2 (16.4, 145) | 32.5 (17.7, 45.6) | 10.5 (3.02, 36.7) | 161 (161, 161) |
| Range | (3.02, 232) | (3.02, 1625) | (3.02, 1602) | (6.46, 1625) | (3.02, 36.7) | (161, 161) |
| **V4a** |  |  |  |  |  |  |
| n | 11 | 34 | 22 | 9 | 3 | 0 |
| Median (IQR) | 58.6 (17.3, 88.3) | 39.2 (11.8, 72.3) | 46.3 (17.3, 77.2) | 25.2 (12.2, 43.0) | 10.8 (3.37, 24.0) | – |
| Range | (3.02, 172) | (3.37, 643) | (3.63, 226) | (5.57, 643) | (3.37, 24.0) | – |
| **V6** |  |  |  |  |  |  |
| n | 11 | 41 | 26 | 11 | 3 | 1 |
| Median (IQR) | 1169 (431, 3859) | 893 (323, 2291) | 1231 (323, 2906) | 621 (420, 1072) | 78.6 (22.8, 779) | 3647 (3647, 3647) |
| Range | (3.02, 4454^b^) | (22.8, 4454) | (57.31, 4454) | (174, 4454) | (22.8, 779) | (3647, 3647) |
| **V8** |  |  |  |  |  |  |
| n | 22 | 63 | 39 | 18 | 5 | 1 |
| Median (IQR) | 260 (123, 624) | 307 (104, 767) | 307 (104, 818) | 267 (132, 650) | 74.3 (22.6, 326) | 1027 (1027, 1027) |
| Range | (3.02, 1199) | (5.57, 4454) | (7.85, 3022) | (5.57, 4454) | (21.6, 741) | (1027, 1027) |
| **V8a** |  |  |  |  |  |  |
| n | 8 | 40 | 27 | 8 | 4 | 1 |
| Median (IQR) | 220 (127, 337) | 124 (67.1, 386) | 128 (67.2, 384) | 123 (88.0, 473) | 32.7 (10.8, 120) | 427 (427, 427) |
| Range | (28.2, 492) | (3.37, 4321) | (7.85, 2321) | (3.37, 4321) | (9.79, 187) | (427, 427) |
| **V8b** |  |  |  |  |  |  |
| n | 9 | 52 | 29 | 18 | 4 | 1 |
| Median (IQR) | 1983 (1320, 3498) | 2655 (1498, 3986) | 3236 (2104, 4086) | 2027 (1395, 3204) | 1567 (750, 2736) | 4454 (4454, 4454) |
| Range | (3.02, 4454) | (196, 4454) | (381, 4454) | (632, 4454) | (196, 3641) | (4454, 4454) |
| **V9** |  |  |  |  |  |  |
| n | 16 | 53 | 33 | 16 | 3 | 1 |
| Median (IQR) | 752 (337, 1275) | 706 (332, 1593) | 756 (468, 2197) | 522 (237, 1099) | 286 (80.9, 643) | 1911 (1911, 1911) |
| Range | (3.02, 4454) | (61.4, 4454) | (61.4, 4454) | (150, 4454) | (80.9, 643) | (1911, 1911) |

^a^3.02 BAU/mL is the LLOD. The seropositivity threshold is 31 BAU/mL.

^b^The values are capped at 4,454 BAU/mL which is the ULOQ indicating saturation for the range of dilutions tested.

**Supplemental table S3B**. Spike IgG levels in serum (BAU/mL) in PWH and HIV^–^ participants. N^+^/post-COVID-19 samples (n=16) and V9 samples preceded by the 4^th^ vaccine dose, D4 (n=3), were excluded.

|  | **HIV–** | **HIV+** | **IRs** | **INRs** | **LLVs** | **LTNPs** |
| --- | --- | --- | --- | --- | --- | --- |
| **V1** |  |  |  |  |  |  |
| n | 11 | 44 | 28 | 12 | 3 | 1 |
| Median (IQR) | 2.16 (1.45, 7.92) | 3.27 (2.32, 5.78) | 3.48 (2.62, 9.72) | 2.87 (2.06, 4.91) | 4.41 (1.26, 5.75) | 2.05 (2.05, 2.05) |
| Range | (1.13^a^, 18.3) | (1.26, 746) | (1.35, 746) | (1.64, 10.2) | (1.26, 5.75) | (2.05, 2.05) |
| **V4** |  |  |  |  |  |  |
| n | 11 | 43 | 28 | 11 | 3 | 1 |
| Median (IQR) | 73.5 (46.6, 204) | 64.7 (26.9, 184) | 77.5 (38.5, 217) | 48.1 (25.3, 68.4) | 25.4 (3.45, 42.0) | 135 (135, 135) |
| Range | (1.13, 437) | (1.16, 1449) | (1.16, 1220) | (8.91, 1449) | (3.45, 42.0) | (135, 135) |
| **V4a** |  |  |  |  |  |  |
| n | 11 | 34 | 22 | 9 | 3 | 0 |
| Median (IQR) | 63.9 (38.6, 97.2) | 43.4 (29.0, 78.6) | 59.1 (30.5, 95.1) | 38.2 (33.3, 59.9) | 28.6 (3.56, 29.5) | – |
| Range | (1.13, 158) | (2.87, 726) | (2.87, 401) | (3.68, 727) | (3.56, 29.5) | – |
| **V6** |  |  |  |  |  |  |
| n | 11 | 41 | 26 | 11 | 3 | 1 |
| Median (IQR) | 935 (560, 1501^b^) | 677 (330, 1501) | 933 (351, 1501) | 437 (330, 1045) | 64.9 (30.0, 636) | 1501 (1501, 1501) |
| Range | (17.2, 1501) | (30.0, 1501) | (49.7, 1501) | (92.0, 1501) | (30.0, 636) | (1501, 1501) |
| **V8** |  |  |  |  |  |  |
| n | 22 | 63 | 39 | 18 | 5 | 1 |
| Median (IQR) | 384 (170, 652) | 244 (98.3, 776) | 289 (98.3, 811) | 301 (123, 684) | 74.6 (34.6, 203) | 1108 (1108, 1108) |
| Range | (22.2, 1211) | (15.8, 1501) | (15.8, 1501) | (22.1, 1501) | (23.9, 757) | (1108, 1108) |
| **V8a** |  |  |  |  |  |  |
| n | 8 | 40 | 27 | 8 | 4 | 1 |
| Median (IQR) | 216 (127, 279) | 117 (64.6, 522) | 118 (64.0, 496) | 106 (70.9, 586) | 44.6 (18.8, 99.7) | 663 (663, 663) |
| Range | (38.3, 686) | (14.9, 1501) | (14.9, 1501) | (62.3, 1501) | (15.5, 132) | (663, 663) |
| **V8b** |  |  |  |  |  |  |
| n | 9 | 52 | 29 | 18 | 4 | 1 |
| Median (IQR) | 1470 (1312, 1501) | 1501 (1337, 1501) | 1501 (1501, 1501) | 1490 (1209, 1501) | 1251 (622, 1436) | 1501 (1501, 1501) |
| Range | (69.0, 1501) | (113, 1501) | (519, 1501) | (925, 1501) | (113, 1501) | (1501, 1501) |
| **V9** |  |  |  |  |  |  |
| n | 16 | 53 | 33 | 16 | 3 | 1 |
| Median (IQR) | 919 (619, 1479) | 916 (636, 1501) | 1138 (690, 1501) | 707 (462, 1501) | 361 (71.6, 807) | 1501 (1501, 1501) |
| Range | (23.6, 1501) | (71.6, 1501) | (88.8, 1501) | (167, 1501) | (71.6, 807) | (1501, 1501) |

^a^1.13 BAU/mL is the LLOD. The seropositivity threshold is 11.3 BAU/mL.

^b^The values are capped at 1,501 BAU/mL which is the ULOQ indicating saturation for the range of dilutions tested.

**Supplemental table S4A**. Intergroup/subgroup comparisons of anti-RBD IgG levels in sera (difference in mean, log*_e_* BAU/mL) for different time points by adjusted mixed effects linear regression. For comparison at baseline (V1), the analysis was adjusted for age. For comparisons at time points between D1 and D2 (V4 and V4a), we adjusted for age and D1 vaccine type (mRNA vs ChAdOx1). For comparisons post-D2 (V6 and onward), we adjusted for age, vaccine regimen for D1 and D2 (mRNA/mRNA, mRNA/ChAdOx1, and ChAdOx1/ChAdOx1), and time lapse between D1 and D2. For comparison at V9, we adjusted for time lapse between D3 and V9. V9 samples preceded by D4 were excluded (n=3).

|  | **All data** | | **Out-of-window data excluded** | | **N+/post-COVID-19 data excluded** | | **N+/post-COVID-19 and out-of-window data excluded** | |
| --- | --- | --- | --- | --- | --- | --- | --- | --- |
|  | **Difference in mean, log (95% CI)** | ***p*** | **Difference in mean, log (95% CI)** | ***p*** | **Difference in mean, log (95% CI)** | ***p*** | **Difference in mean, log (95% CI)** | ***p*** |
| **V1** |  |  |  |  |  |  |  |  |
| HIV+ vs HIV– | -0.08 (-0.83, 0.67) | 0.832 | -0.10 (-0.85, 0.65) | 0.800 | 0.11 (-0.59, 0.82) | 0.755 | 0.11 (-0.60, 0.81) | 0.767 |
| IR vs HIV– | -0.05 (-0.84, 0.74) | 0.903 | -0.05 (-0.83, 0.74) | 0.907 | 0.15 (-0.58, 0.89) | 0.683 | 0.16 (-0.58, 0.89) | 0.677 |
| INR vs HIV– | -0.18 (-1.12, 0.75) | 0.699 | -0.20 (-1.13, 0.73) | 0.675 | 0.06 (-0.80, 0.92) | 0.891 | 0.05 (-0.81, 0.91) | 0.908 |
| INR vs IR | -0.13 (-0.93, 0.66) | 0.740 | -0.15 (-0.94, 0.64) | 0.707 | -0.09 (-0.81, 0.63) | 0.800 | -0.11 (-0.83, 0.61) | 0.772 |
| **V4** |  |  |  |  |  |  |  |  |
| HIV+ vs HIV– | -0.13 (-0.88, 0.63) | 0.737 | -0.18 (-0.93, 0.58) | 0.642 | 0.07 (-0.63, 0.78) | 0.835 | 0.09 (-0.62, 0.80) | 0.808 |
| IR vs HIV– | -0.07 (-0.86, 0.72) | 0.863 | -0.06 (-0.85, 0.73) | 0.878 | 0.29 (-0.45, 1.03) | 0.436 | 0.30 (-0.44, 1.04) | 0.429 |
| INR vs HIV– | -0.07 (-1.00, 0.87) | 0.888 | -0.24 (-1.20, 0.72) | 0.619 | -0.18 (-1.05, 0.69) | 0.691 | -0.13 (-1.02, 0.75) | 0.768 |
| INR vs IR | 0.00 (-0.80, 0.80) | 0.995 | -0.18 (-1.01, 0.65) | 0.667 | -0.47 (-1.21, 0.27) | 0.213 | -0.43 (-1.18, 0.32) | 0.262 |
| **V4a** |  |  |  |  |  |  |  |  |
| HIV+ vs HIV– | -0.12 (-0.89, 0.64) | 0.753 | -0.13 (-0.90, 0.63) | 0.736 | 0.02 (-0.70, 0.74) | 0.956 | 0.02 (-0.70, 0.74) | 0.962 |
| IR vs HIV– | -0.14 (-0.95, 0.67) | 0.731 | -0.13 (-0.94, 0.67) | 0.746 | 0.17 (-0.58, 0.92) | 0.654 | 0.18 (-0.58, 0.93) | 0.643 |
| INR vs HIV– | 0.15 (-0.81, 1.11) | 0.761 | 0.15 (-0.81, 1.11) | 0.762 | -0.06 (-0.96, 0.84) | 0.901 | -0.06 (-0.96, 0.84) | 0.896 |
| INR vs IR | 0.29 (-0.56, 1.14) | 0.501 | 0.28 (-0.57, 1.13) | 0.514 | -0.23 (-1.01, 0.56) | 0.568 | -0.24 (-1.02, 0.55) | 0.554 |
| **V6** |  |  |  |  |  |  |  |  |
| HIV+ vs HIV– | 0.12 (-0.63, 0.88) | 0.745 | 0.12 (-0.64, 0.88) | 0.757 | 0.17 (-0.54, 0.88) | 0.642 | 0.19 (-0.53, 0.90) | 0.606 |
| IR vs HIV– | 0.22 (-0.58, 1.01) | 0.593 | 0.22 (-0.58, 1.02) | 0.584 | 0.38 (-0.36, 1.12) | 0.317 | 0.40 (-0.34, 1.15) | 0.288 |
| INR vs HIV– | 0.21 (-0.72, 1.14) | 0.657 | 0.11 (-0.87, 1.09) | 0.826 | 0.08 (-0.79, 0.95) | 0.862 | -0.00 (-0.90, 0.90) | 0.998 |
| INR vs IR | -0.00 (-0.81, 0.80) | 0.990 | -0.11 (-0.98, 0.75) | 0.796 | -0.30 (-1.04, 0.44) | 0.426 | -0.41 (-1.19, 0.38) | 0.308 |
| **V8** |  |  |  |  |  |  |  |  |
| HIV+ vs HIV– | 0.19 (-0.43, 0.81) | 0.547 | 0.12 (-0.51, 0.75) | 0.708 | 0.20 (-0.37, 0.78) | 0.487 | 0.10 (-0.49, 0.69) | 0.731 |
| IR vs HIV– | 0.15 (-0.50, 0.81) | 0.642 | 0.07 (-0.59, 0.73) | 0.836 | 0.25 (-0.35, 0.86) | 0.409 | 0.15 (-0.46, 0.77) | 0.624 |
| INR vs HIV– | 0.42 (-0.36, 1.21) | 0.286 | 0.45 (-0.35, 1.25) | 0.267 | 0.26 (-0.47, 0.98) | 0.485 | 0.26 (-0.48, 1.00) | 0.494 |
| INR vs IR | 0.27 (-0.43, 0.97) | 0.450 | 0.38 (-0.33, 1.10) | 0.295 | 0.00 (-0.65, 0.65) | 0.993 | 0.11 (-0.56, 0.77) | 0.755 |
| **V8a** |  |  |  |  |  |  |  |  |
| HIV+ vs HIV– | -0.13 (-0.93, 0.66) | 0.739 | -0.15 (-0.94, 0.64) | 0.708 | 0.27 (-0.49, 1.04) | 0.483 | 0.25 (-0.52, 1.02) | 0.528 |
| IR vs HIV– | -0.18 (-1.01, 0.64) | 0.664 | -0.20 (-1.02, 0.62) | 0.633 | 0.34 (-0.45, 1.13) | 0.396 | 0.32 (-0.47, 1.11) | 0.428 |
| INR vs HIV– | 0.54 (-0.49, 1.58) | 0.303 | 0.56 (-0.47, 1.60) | 0.282 | 0.69 (-0.30, 1.67) | 0.174 | 0.69 (-0.31, 1.68) | 0.175 |
| INR vs IR | 0.72 (-0.16, 1.60) | 0.106 | 0.76 (-0.11, 1.64) | 0.087 | 0.34 (-0.48, 1.17) | 0.411 | 0.37 (-0.46, 1.19) | 0.382 |
| **V8b** |  |  |  |  |  |  |  |  |
| HIV+ vs HIV– | 0.53 (-0.23, 1.29) | 0.171 | 0.52 (-0.23, 1.28) | 0.174 | 0.70 (-0.04, 1.43) | 0.063 | 0.70 (-0.04, 1.43) | 0.063 |
| IR vs HIV– | 0.70 (-0.09, 1.50) | 0.084 | 0.70 (-0.09, 1.50) | 0.082 | 0.89 (0.13, 1.66) | 0.022 | 0.89 (0.13, 1.66) | 0.022 |
| INR vs HIV– | 0.34 (-0.55, 1.23) | 0.454 | 0.34 (-0.54, 1.23) | 0.448 | 0.47 (-0.38, 1.32) | 0.276 | 0.47 (-0.37, 1.32) | 0.271 |
| INR vs IR | -0.36 (-1.09, 0.36) | 0.323 | -0.36 (-1.08, 0.35) | 0.320 | -0.42 (-1.09, 0.25) | 0.215 | -0.42 (-1.09, 0.25) | 0.216 |
| **V9** |  |  |  |  |  |  |  |  |
| HIV+ vs HIV– | 0.39 (-0.24, 1.02) | 0.228 | 0.39 (-0.24, 1.02) | 0.228 | 0.41 (-0.22, 1.04) | 0.198 | 0.47 (-0.16, 1.11) | 0.144 |
| IR vs HIV– | 0.41 (-0.25, 1.08) | 0.217 | 0.46 (-0.21, 1.12) | 0.176 | 0.46 (-0.19, 1.12) | 0.165 | 0.52 (-0.15, 1.19) | 0.125 |
| INR vs HIV– | 0.29 (-0.55, 1.12) | 0.500 | 0.37 (-0.46, 1.21) | 0.378 | 0.43 (-0.37, 1.22) | 0.290 | 0.52 (-0.27, 1.32) | 0.195 |
| INR vs IR | -0.13 (-0.88, 0.63) | 0.740 | -0.08 (-0.83, 0.67) | 0.832 | -0.04 (-0.75, 0.67) | 0.915 | 0.00 (-0.71, 0.72) | 0.989 |

**Supplemental table S4B**. Intergroup/subgroup comparisons of anti-spike IgG levels in sera (difference in mean, log*_e_* BAU/mL) for different time points by adjusted mixed effects linear regression. For comparison at baseline (V1), the analysis was adjusted for age. For comparisons at time points between D1 and D2 (V4 and V4a), we adjusted for age and D1 vaccine type (mRNA vs ChAdOx1). For comparisons post-D2 (V6 and onward), we adjusted for age, vaccine regimen for D1 and D2 (mRNA/mRNA, mRNA/ChAdOx1, and ChAdOx1/ChAdOx1), and time lapse between D1 and D2. For comparison at V9, we adjusted for time lapse between D3 and V9. V9 samples preceded by D4 were excluded (n=3).

|  | **All data** | | **Out-of-window data excluded** | | **N+/post-COVID-19 data excluded** | | **N+/post-COVID-19 and out-of-window data excluded** | |
| --- | --- | --- | --- | --- | --- | --- | --- | --- |
|  | **Difference in mean, log (95% CI)** | ***p*** | **Difference in mean, log (95% CI)** | ***p*** | **Difference in mean, log (95% CI)** | ***p*** | **Difference in mean, log (95% CI)** | ***p*** |
| **V1** |  |  |  |  |  |  |  |  |
| HIV+ vs HIV– | 0.04 (-0.57, 0.66) | 0.889 | 0.03 (-0.59, 0.64) | 0.935 | 0.28 (-0.32, 0.87) | 0.359 | 0.27 (-0.33, 0.87) | 0.379 |
| IR vs HIV– | 0.17 (-0.47, 0.82) | 0.597 | 0.17 (-0.48, 0.81) | 0.612 | 0.43 (-0.19, 1.04) | 0.171 | 0.42 (-0.19, 1.04) | 0.178 |
| INR vs HIV– | -0.17 (-0.93, 0.59) | 0.653 | -0.19 (-0.95, 0.57) | 0.625 | 0.08 (-0.64, 0.79) | 0.831 | 0.07 (-0.65, 0.78) | 0.855 |
| INR vs IR | -0.35 (-0.99, 0.30) | 0.293 | -0.36 (-1.00, 0.29) | 0.280 | -0.35 (-0.95, 0.25) | 0.251 | -0.36 (-0.96, 0.25) | 0.244 |
| **V4** |  |  |  |  |  |  |  |  |
| HIV+ vs HIV– | -0.14 (-0.76, 0.47) | 0.648 | -0.17 (-0.79, 0.45) | 0.594 | 0.09 (-0.51, 0.69) | 0.773 | 0.11 (-0.49, 0.71) | 0.719 |
| IR vs HIV– | 0.01 (-0.64, 0.65) | 0.982 | 0.01 (-0.64, 0.65) | 0.983 | 0.36 (-0.25, 0.98) | 0.247 | 0.36 (-0.26, 0.98) | 0.251 |
| INR vs HIV– | -0.20 (-0.96, 0.56) | 0.601 | -0.29 (-1.07, 0.50) | 0.473 | -0.20 (-0.92, 0.52) | 0.584 | -0.12 (-0.86, 0.63) | 0.760 |
| INR vs IR | -0.21 (-0.86, 0.44) | 0.525 | -0.29 (-0.97, 0.39) | 0.395 | -0.56 (-1.18, 0.05) | 0.071 | -0.48 (-1.11, 0.15) | 0.138 |
| **V4a** |  |  |  |  |  |  |  |  |
| HIV+ vs HIV– | -0.09 (-0.71, 0.54) | 0.786 | -0.10 (-0.73, 0.53) | 0.761 | 0.09 (-0.51, 0.70) | 0.760 | 0.09 (-0.52, 0.70) | 0.771 |
| IR vs HIV– | -0.04 (-0.71, 0.62) | 0.896 | -0.04 (-0.71, 0.62) | 0.896 | 0.26 (-0.36, 0.89) | 0.406 | 0.27 (-0.37, 0.90) | 0.409 |
| INR vs HIV– | 0.05 (-0.73, 0.84) | 0.893 | 0.06 (-0.73, 0.84) | 0.889 | -0.02 (-0.77, 0.73) | 0.961 | -0.01 (-0.77, 0.74) | 0.970 |
| INR vs IR | 0.10 (-0.60, 0.79) | 0.782 | 0.10 (-0.59, 0.79) | 0.778 | -0.28 (-0.94, 0.37) | 0.397 | -0.28 (-0.94, 0.38) | 0.405 |
| **V6** |  |  |  |  |  |  |  |  |
| HIV+ vs HIV– | -0.18 (-0.80, 0.44) | 0.573 | -0.17 (-0.80, 0.46) | 0.590 | -0.12 (-0.72, 0.47) | 0.682 | -0.10 (-0.71, 0.50) | 0.738 |
| IR vs HIV– | -0.10 (-0.74, 0.55) | 0.768 | -0.10 (-0.76, 0.55) | 0.754 | 0.03 (-0.58, 0.65) | 0.919 | 0.04 (-0.58, 0.67) | 0.894 |
| INR vs HIV– | -0.07 (-0.83, 0.69) | 0.862 | -0.10 (-0.90, 0.71) | 0.813 | -0.11 (-0.83, 0.62) | 0.770 | -0.14 (-0.89, 0.62) | 0.720 |
| INR vs IR | 0.03 (-0.62, 0.68) | 0.929 | 0.01 (-0.70, 0.72) | 0.983 | -0.14 (-0.75, 0.47) | 0.656 | -0.18 (-0.84, 0.48) | 0.589 |
| **V8** |  |  |  |  |  |  |  |  |
| HIV+ vs HIV– | -0.10 (-0.60, 0.41) | 0.699 | -0.13 (-0.65, 0.39) | 0.617 | -0.10 (-0.59, 0.38) | 0.672 | -0.16 (-0.66, 0.34) | 0.525 |
| IR vs HIV– | -0.10 (-0.62, 0.43) | 0.721 | -0.11 (-0.65, 0.43) | 0.687 | -0.05 (-0.54, 0.45) | 0.856 | -0.08 (-0.59, 0.43) | 0.763 |
| INR vs HIV– | 0.07 (-0.56, 0.71) | 0.819 | 0.09 (-0.56, 0.74) | 0.785 | -0.03 (-0.62, 0.56) | 0.920 | -0.04 (-0.65, 0.58) | 0.904 |
| INR vs IR | 0.17 (-0.40, 0.74) | 0.557 | 0.20 (-0.38, 0.78) | 0.498 | 0.02 (-0.52, 0.55) | 0.955 | 0.04 (-0.51, 0.59) | 0.886 |
| **V8a** |  |  |  |  |  |  |  |  |
| HIV+ vs HIV– | -0.17 (-0.82, 0.48) | 0.617 | -0.18 (-0.83, 0.47) | 0.592 | 0.09 (-0.56, 0.74) | 0.783 | 0.08 (-0.58, 0.73) | 0.818 |
| IR vs HIV– | -0.18 (-0.85, 0.49) | 0.603 | -0.19 (-0.86, 0.49) | 0.585 | 0.15 (-0.51, 0.82) | 0.645 | 0.15 (-0.52, 0.81) | 0.668 |
| INR vs HIV– | 0.24 (-0.60, 1.09) | 0.570 | 0.25 (-0.59, 1.10) | 0.557 | 0.32 (-0.51, 1.15) | 0.445 | 0.32 (-0.52, 1.16) | 0.455 |
| INR vs IR | 0.42 (-0.30, 1.14) | 0.248 | 0.44 (-0.28, 1.16) | 0.230 | 0.17 (-0.52, 0.86) | 0.632 | 0.17 (-0.52, 0.87) | 0.624 |
| **V8b** |  |  |  |  |  |  |  |  |
| HIV+ vs HIV– | 0.04 (-0.58, 0.66) | 0.908 | 0.02 (-0.60, 0.65) | 0.940 | 0.14 (-0.48, 0.76) | 0.652 | 0.14 (-0.49, 0.76) | 0.669 |
| IR vs HIV– | 0.13 (-0.52, 0.78) | 0.685 | 0.13 (-0.52, 0.78) | 0.700 | 0.26 (-0.38, 0.90) | 0.422 | 0.26 (-0.39, 0.90) | 0.434 |
| INR vs HIV– | -0.02 (-0.74, 0.71) | 0.966 | -0.03 (-0.75, 0.70) | 0.941 | 0.12 (-0.59, 0.82) | 0.741 | 0.11 (-0.60, 0.82) | 0.764 |
| INR vs IR | -0.15 (-0.73, 0.44) | 0.614 | -0.16 (-0.74, 0.43) | 0.601 | -0.14 (-0.69, 0.41) | 0.612 | -0.15 (-0.70, 0.41) | 0.600 |
| **V9** |  |  |  |  |  |  |  |  |
| HIV+ vs HIV– | 0.10 (-0.42, 0.61) | 0.716 | 0.10 (-0.42, 0.63) | 0.692 | 0.13 (-0.40, 0.66) | 0.624 | 0.16 (-0.37, 0.70) | 0.547 |
| IR vs HIV– | 0.13 (-0.40, 0.66) | 0.634 | 0.16 (-0.38, 0.70) | 0.550 | 0.16 (-0.38, 0.71) | 0.551 | 0.20 (-0.35, 0.76) | 0.475 |
| INR vs HIV– | 0.10 (-0.58, 0.78) | 0.775 | 0.14 (-0.55, 0.82) | 0.693 | 0.22 (-0.43, 0.88) | 0.501 | 0.26 (-0.40, 0.92) | 0.441 |
| INR vs IR | -0.03 (-0.64, 0.58) | 0.921 | -0.03 (-0.64, 0.59) | 0.932 | 0.06 (-0.53, 0.65) | 0.843 | 0.06 (-0.54, 0.65) | 0.846 |

**Supplemental table S5A**. Within-group/subgroup changes in anti-RBD IgG levels in sera (difference in mean, log*_e_* BAU/mL) between the neighbouring time points by regression analysis. V9 samples preceded by D4 were excluded (n=3).

|  | **All data** | | **Out-of-window data excluded** | | **N+/post-COVID-19 data excluded** | | **N+/post-COVID-19 and out-of-window data excluded** | |
| --- | --- | --- | --- | --- | --- | --- | --- | --- |
|  | **Difference in mean, log (95% CI)** | ***p*** | **Difference in mean, log (95% CI)** | ***p*** | **Difference in mean, log (95% CI)** | ***p*** | **Difference in mean, log (95% CI)** | ***p*** |
| **V4 vs V1** |  |  |  |  |  |  |  |  |
| HIV– | 2.54 (1.79, 3.29) | <0.001 | 2.50 (1.75, 3.25) | <0.001 | 2.38 (1.69, 3.07) | <0.001 | 2.37 (1.67, 3.07) | <0.001 |
| HIV+ | 2.49 (2.05, 2.93) | <0.001 | 2.42 (1.97, 2.86) | <0.001 | 2.34 (1.94, 2.74) | <0.001 | 2.35 (1.95, 2.76) | <0.001 |
| IR | 2.54 (2.02, 3.05) | <0.001 | 2.50 (1.98, 3.02) | <0.001 | 2.51 (2.04, 2.98) | <0.001 | 2.50 (2.03, 2.97) | <0.001 |
| INR | 2.67 (1.92, 3.42) | <0.001 | 2.47 (1.68, 3.26) | <0.001 | 2.14 (1.45, 2.83) | <0.001 | 2.18 (1.47, 2.89) | <0.001 |
| **V4a vs V4** |  |  |  |  |  |  |  |  |
| HIV– | -0.36 (-1.06, 0.34) | 0.310 | -0.36 (-1.06, 0.34) | 0.312 | -0.34 (-0.98, 0.30) | 0.295 | -0.34 (-0.98, 0.30) | 0.299 |
| HIV+ | -0.35 (-0.75, 0.04) | 0.076 | -0.31 (-0.71, 0.08) | 0.122 | -0.39 (-0.74, -0.05) | 0.027 | -0.41 (-0.76, -0.06) | 0.023 |
| IR | -0.43 (-0.92, 0.06) | 0.084 | -0.43 (-0.93, 0.06) | 0.086 | -0.46 (-0.89, -0.03) | 0.037 | -0.46 (-0.90, -0.02) | 0.039 |
| INR | -0.14 (-0.88, 0.59) | 0.699 | 0.03 (-0.74, 0.80) | 0.939 | -0.22 (-0.90, 0.45) | 0.520 | -0.27 (-0.96, 0.43) | 0.450 |
| **V6 vs V4a** |  |  |  |  |  |  |  |  |
| HIV– | 2.03 (0.69, 3.37) | 0.003 | 2.08 (0.74, 3.41) | 0.002 | 2.58 (1.34, 3.82) | <0.001 | 2.61 (1.37, 3.84) | <0.001 |
| HIV+ | 2.28 (1.06, 3.50) | <0.001 | 2.33 (1.11, 3.55) | <0.001 | 2.73 (1.60, 3.85) | <0.001 | 2.78 (1.65, 3.91) | <0.001 |
| IR | 2.18 (0.92, 3.43) | <0.001 | 2.23 (0.98, 3.48) | <0.001 | 2.56 (1.41, 3.70) | <0.001 | 2.61 (1.46, 3.76) | <0.001 |
| INR | 1.88 (0.51, 3.25) | 0.007 | 1.84 (0.44, 3.24) | 0.010 | 2.48 (1.22, 3.75) | <0.001 | 2.44 (1.16, 3.72) | <0.001 |
| **V8 vs V6** |  |  |  |  |  |  |  |  |
| HIV– | -1.27 (-1.91, -0.63) | <0.001 | -1.25 (-1.92, -0.59) | <0.001 | -1.29 (-1.88, -0.70) | <0.001 | -1.25 (-1.87, -0.64) | <0.001 |
| HIV+ | -1.21 (-1.55, -0.86) | <0.001 | -1.25 (-1.63, -0.87) | <0.001 | -1.26 (-1.57, -0.95) | <0.001 | -1.34 (-1.68, -1.00) | <0.001 |
| IR | -1.33 (-1.76, -0.89) | <0.001 | -1.39 (-1.85, -0.93) | <0.001 | -1.41 (-1.80, -1.02) | <0.001 | -1.49 (-1.91, -1.08) | <0.001 |
| INR | -1.05 (-1.71, -0.40) | 0.002 | -0.89 (-1.64, -0.15) | 0.019 | -1.11 (-1.70, -0.51) | <0.001 | -0.98 (-1.64, -0.32) | 0.004 |
| **V8a vs V8** |  |  |  |  |  |  |  |  |
| HIV– | -0.11 (-0.80, 0.57) | 0.747 | -0.12 (-0.82, 0.59) | 0.742 | -0.51 (-1.17, 0.15) | 0.130 | -0.53 (-1.21, 0.15) | 0.126 |
| HIV+ | -0.44 (-0.79, -0.08) | 0.015 | -0.39 (-0.75, -0.03) | 0.035 | -0.44 (-0.75, -0.12) | 0.007 | -0.39 (-0.71, -0.06) | 0.020 |
| IR | -0.45 (-0.88, -0.02) | 0.039 | -0.40 (-0.84, 0.04) | 0.074 | -0.43 (-0.81, -0.04) | 0.029 | -0.37 (-0.77, 0.02) | 0.061 |
| INR | 0.00 (-0.76, 0.76) | 0.999 | -0.02 (-0.79, 0.76) | 0.966 | -0.09 (-0.79, 0.62) | 0.811 | -0.11 (-0.84, 0.61) | 0.756 |
| **V8b vs V8a** |  |  |  |  |  |  |  |  |
| HIV– | 1.96 (1.20, 2.71) | <0.001 | 1.94 (1.19, 2.70) | <0.001 | 2.16 (1.43, 2.90) | <0.001 | 2.14 (1.40, 2.88) | <0.001 |
| HIV+ | 2.62 (2.26, 2.98) | <0.001 | 2.62 (2.25, 2.98) | <0.001 | 2.59 (2.26, 2.91) | <0.001 | 2.59 (2.26, 2.92) | <0.001 |
| IR | 2.85 (2.40, 3.30) | <0.001 | 2.85 (2.39, 3.30) | <0.001 | 2.72 (2.31, 3.12) | <0.001 | 2.72 (2.31, 3.13) | <0.001 |
| INR | 1.76 (1.00, 2.51) | <0.001 | 1.72 (0.96, 2.48) | <0.001 | 1.95 (1.25, 2.65) | <0.001 | 1.93 (1.22, 2.64) | <0.001 |
| **V9 vs V8b** |  |  |  |  |  |  |  |  |
| HIV– | 1.27 (-0.13, 2.68) | 0.075 | 1.16 (-0.27, 2.59) | 0.110 | 1.15 (-0.24, 2.55) | 0.106 | 1.15 (-0.27, 2.58) | 0.112 |
| HIV+ | 1.13 (-0.19, 2.45) | 0.092 | 1.03 (-0.32, 2.37) | 0.135 | 0.87 (-0.39, 2.12) | 0.176 | 0.93 (-0.35, 2.20) | 0.153 |
| IR | 1.08 (-0.37, 2.53) | 0.145 | 1.24 (-0.23, 2.70) | 0.099 | 0.91 (-0.45, 2.27) | 0.187 | 1.01 (-0.36, 2.39) | 0.149 |
| INR | 1.31 (-0.43, 3.06) | 0.139 | 1.52 (-0.25, 3.28) | 0.092 | 1.30 (-0.35, 2.94) | 0.122 | 1.44 (-0.23, 3.10) | 0.091 |

**Supplemental table S5B**. Within-group/subgroup changes in anti-spike IgG levels in sera (difference in mean, log*_e_* BAU/mL) between the neighbouring time points by regression analysis. V9 samples preceded by D4 were excluded (n=3).

|  | **All data** | | **Out-of-window data excluded** | | **N+/post-COVID-19 data excluded** | | **N+/post-COVID-19 and out-of-window data excluded** | |
| --- | --- | --- | --- | --- | --- | --- | --- | --- |
|  | **Difference in mean, log (95% CI)** | ***p*** | **Difference in mean, log (95% CI)** | ***p*** | **Difference in mean, log (95% CI)** | ***p*** | **Difference in mean, log (95% CI)** | ***p*** |
| **V4 vs V1** |  |  |  |  |  |  |  |  |
| HIV– | 2.82 (2.20, 3.44) | <0.001 | 2.78 (2.16, 3.40) | <0.001 | 2.75 (2.15, 3.35) | <0.001 | 2.73 (2.13, 3.33) | <0.001 |
| HIV+ | 2.63 (2.27, 2.99) | <0.001 | 2.58 (2.22, 2.95) | <0.001 | 2.56 (2.21, 2.91) | <0.001 | 2.57 (2.22, 2.92) | <0.001 |
| IR | 2.65 (2.23, 3.08) | <0.001 | 2.62 (2.19, 3.05) | <0.001 | 2.66 (2.26, 3.07) | <0.001 | 2.65 (2.24, 3.06) | <0.001 |
| INR | 2.79 (2.17, 3.41) | <0.001 | 2.68 (2.02, 3.34) | <0.001 | 2.45 (1.86, 3.05) | <0.001 | 2.53 (1.92, 3.14) | <0.001 |
| **V4a vs V4** |  |  |  |  |  |  |  |  |
| HIV– | -0.42 (-1.00, 0.15) | 0.150 | -0.42 (-1.00, 0.16) | 0.152 | -0.41 (-0.97, 0.14) | 0.140 | -0.41 (-0.97, 0.14) | 0.142 |
| HIV+ | -0.37 (-0.69, -0.04) | 0.027 | -0.35 (-0.68, -0.02) | 0.036 | -0.41 (-0.71, -0.11) | 0.008 | -0.43 (-0.74, -0.13) | 0.005 |
| IR | -0.47 (-0.88, -0.07) | 0.023 | -0.47 (-0.89, -0.06) | 0.024 | -0.51 (-0.88, -0.14) | 0.007 | -0.51 (-0.89, -0.13) | 0.008 |
| INR | -0.17 (-0.78, 0.44) | 0.591 | -0.08 (-0.72, 0.56) | 0.806 | -0.23 (-0.81, 0.35) | 0.435 | -0.31 (-0.91, 0.29) | 0.305 |
| **V6 vs V4a** |  |  |  |  |  |  |  |  |
| HIV– | 2.01 (0.91, 3.10) | <0.001 | 2.03 (0.93, 3.13) | <0.001 | 2.37 (1.32, 3.42) | <0.001 | 2.38 (1.33, 3.43) | <0.001 |
| HIV+ | 1.91 (0.92, 2.91) | <0.001 | 1.96 (0.95, 2.96) | <0.001 | 2.15 (1.20, 3.10) | <0.001 | 2.19 (1.23, 3.14) | <0.001 |
| IR | 1.86 (0.84, 2.88) | <0.001 | 1.89 (0.86, 2.91) | <0.001 | 2.04 (1.09, 2.99) | <0.001 | 2.06 (1.10, 3.02) | <0.001 |
| INR | 1.79 (0.67, 2.91) | 0.002 | 1.80 (0.65, 2.94) | 0.002 | 2.18 (1.13, 3.23) | <0.001 | 2.16 (1.08, 3.24) | <0.001 |
| **V8 vs V6** |  |  |  |  |  |  |  |  |
| HIV– | -0.94 (-1.47, -0.41) | <0.001 | -0.98 (-1.53, -0.43) | <0.001 | -0.91 (-1.42, -0.40) | <0.001 | -0.94 (-1.47, -0.41) | <0.001 |
| HIV+ | -0.86 (-1.15, -0.57) | <0.001 | -0.94 (-1.26, -0.63) | <0.001 | -0.89 (-1.16, -0.62) | <0.001 | -1.00 (-1.29, -0.71) | <0.001 |
| IR | -0.93 (-1.29, -0.57) | <0.001 | -0.97 (-1.36, -0.59) | <0.001 | -0.98 (-1.32, -0.64) | <0.001 | -1.04 (-1.40, -0.68) | <0.001 |
| INR | -0.79 (-1.33, -0.24) | 0.005 | -0.78 (-1.40, -0.16) | 0.014 | -0.83 (-1.34, -0.31) | 0.002 | -0.82 (-1.40, -0.25) | 0.005 |
| **V8a vs V8** |  |  |  |  |  |  |  |  |
| HIV– | -0.38 (-0.95, 0.18) | 0.184 | -0.34 (-0.92, 0.24) | 0.253 | -0.66 (-1.23, -0.09) | 0.024 | -0.63 (-1.22, -0.04) | 0.035 |
| HIV+ | -0.45 (-0.74, -0.16) | 0.003 | -0.38 (-0.68, -0.09) | 0.012 | -0.46 (-0.73, -0.19) | <0.001 | -0.39 (-0.67, -0.11) | 0.006 |
| IR | -0.47 (-0.83, -0.11) | 0.010 | -0.43 (-0.79, -0.06) | 0.022 | -0.46 (-0.79, -0.13) | 0.006 | -0.42 (-0.76, -0.08) | 0.016 |
| INR | -0.22 (-0.85, 0.41) | 0.498 | -0.19 (-0.83, 0.46) | 0.567 | -0.31 (-0.92, 0.30) | 0.316 | -0.29 (-0.91, 0.34) | 0.368 |
| **V8b vs V8a** |  |  |  |  |  |  |  |  |
| HIV– | 1.86 (1.24, 2.49) | <0.001 | 1.87 (1.24, 2.49) | <0.001 | 1.99 (1.35, 2.63) | <0.001 | 1.99 (1.35, 2.63) | <0.001 |
| HIV+ | 2.07 (1.77, 2.36) | <0.001 | 2.07 (1.77, 2.37) | <0.001 | 2.04 (1.76, 2.32) | <0.001 | 2.05 (1.76, 2.33) | <0.001 |
| IR | 2.17 (1.80, 2.55) | <0.001 | 2.18 (1.80, 2.55) | <0.001 | 2.09 (1.74, 2.44) | <0.001 | 2.09 (1.74, 2.44) | <0.001 |
| INR | 1.60 (0.97, 2.23) | <0.001 | 1.58 (0.95, 2.22) | <0.001 | 1.78 (1.17, 2.39) | <0.001 | 1.77 (1.16, 2.39) | <0.001 |
| **V9 vs V8b** |  |  |  |  |  |  |  |  |
| HIV– | 0.73 (-0.42, 1.89) | 0.214 | 0.63 (-0.55, 1.80) | 0.297 | 0.76 (-0.44, 1.97) | 0.214 | 0.75 (-0.47, 1.98) | 0.227 |
| HIV+ | 0.79 (-0.30, 1.88) | 0.153 | 0.71 (-0.40, 1.82) | 0.212 | 0.75 (-0.33, 1.84) | 0.173 | 0.78 (-0.32, 1.88) | 0.162 |
| IR | 0.81 (-0.40, 2.01) | 0.189 | 0.87 (-0.36, 2.09) | 0.165 | 0.95 (-0.22, 2.12) | 0.111 | 0.98 (-0.21, 2.18) | 0.105 |
| INR | 0.93 (-0.52, 2.37) | 0.210 | 0.99 (-0.47, 2.46) | 0.184 | 1.15 (-0.27, 2.57) | 0.111 | 1.19 (-0.25, 2.63) | 0.105 |

**Supplemental table S6**. Live SARS-CoV-2 virus 50% neutralization titers (NT50) at baseline (V1) assessed as a dichotomized variable (0 vs. above 0). *P* value is based on Chi-square test or Fisher’s exact test as appropriate. Formal comparisons involving the LLV and LNTP subgroups were not performed as the sample size was too small.

| **Neutralization titers** | **HIV–** | **HIV+** | **IRs** | **INRs** | **LLVs** | **LTNPs** |
| --- | --- | --- | --- | --- | --- | --- |
| **V1** |  |  |  |  |  |  |
| Above 0, n (%) | 2/12 (16.7) | 6/45 (13.3) | 1/29 (3.4) | 3/12 (25.0) | 2/3 (66.7) | 0/1 (0.0) |
| *p* value |  |  |  |  |  |  |
| Compared to HIV– | – | 0.768 | 0.200 | 1.000 | – | – |
| Compared to IR | – | – | – | 0.068 | – | – |
| **V1** (N^+^ data excluded) |  |  |  |  |  |  |
| Above 0, n (%) | 2/11 (18.2) | 6/44 (13.6) | 1/28 (3.6) | 3/12 (25.0) | 2/3 (66.7) | 0/1 (0.0) |
| *p* value |  |  |  |  |  |  |
| Compared to HIV– | – | 0.702 | 0.187 | 1.000 | – | – |
| Compared to IR | – | – | – | 0.073 | – | – |

**Supplemental table S7**. Live SARS-CoV-2 50% neutralization titers (NT50) at 24 weeks (V8) and 48 weeks (V9) post-D1 by adjusted quantile regression. Adjusted for age, vaccine regimen for D1 and D2 (mRNA/mRNA, mRNA/ChAdOx1, and ChAdOx1/ChAdOx1), and time lapse between D1 and D2. Formal comparisons involving the LLV and LTNP subgroups were not performed as the sample size was too small.

|  | **HIV–** | **HIV+** | **IRs** | **INRs** | **LLVs** | **LTNPs** |
| --- | --- | --- | --- | --- | --- | --- |
| **V8** |  |  |  |  |  |  |
| **n** | 23 | 66 | 41 | 19 | 5 | 1 |
| **Median (IQR)** | 618 (154, 640) | 82.9 (38.6, 160) | 82.9 (38.6, 165.8) | 113.1 (57.7, 160.0) | 77.2 (27.9, 80.0) | 56.6 (56.6, 56.6) |
| **Range** | (20.0, 1280) | (0.0, 1235) | (0.0, 1235.0) | (0.0, 1235.0) | (20.0, 82.9) | (56.6, 56.6) |
| **V9** |  |  |  |  |  |  |
| **N** | 22 | 62 | 39 | 17 | 5 | 1 |
| **Median (IQR)** | 314 (160, 1327) | 332 (154, 2560) | 450 (154, 2560) | 188 (154, 1327) | 309 (160, 2560) | 116 (116, 116) |
| **Range** | (0.0, 2560) | (0.0, 2560) | (0.0, 2560) | (38.6, 2560) | (0.0, 2560) | (116, 116) |
| **Change from V8 to V9** |  |  |  |  |  |  |
| **n** | 22 | 60 | 38 | 16 |  |  |
| **Median (IQR)** | -21.4 (-458, 828) | 200 (61.3, 1863) | 345 (63.6, 1920) | 137 (61.7, 1285) |  |  |
| **Range** | (-1122, 2394) | (-903, 2560) | (-903, 2560) | (-121, 2400) |  |  |
| ***p*** | 0.913 | 0.310 | 0.250 | 0.572 |  |  |
| **V8 (Out-of-window data excluded)** |  |  |  |  |  |  |
| **n** | 21 | 60 | 39 | 17 | 4 | 0 |
| **Median (IQR)** | 640 (226, 640) | 82.9 (27.9, 160) | 82.9 (27.9, 166) | 154 (57.7, 160) | 52.5 (23.9, 78.6) | – |
| **Range** | (82.9, 1280) | (0.0, 1235) | (0.0, 1235) | (0.0, 1235) | (20.0, 80.0) | – |
| **V9 (Out-of-window data excluded)** |  |  |  |  |  |  |
| **n** | 21 | 60 | 38 | 17 | 4 | 1 |
| **Median (IQR)** | 320 (160, 1327) | 326 (154, 2144) | 435 (154, 2560) | 188 (154, 1327) | 234 (80.0, 1434) | 116 (116, 116) |
| **Range** | (0.0, 2560) | (0.0, 2560) | (0.0, 2560) | (38.6, 2560) | (0.0, 2560) | (116, 116) |
| **Change from V8 to V9 (Out-of-window data excluded)** |  |  |  |  |  |  |
| **n** | 19 | 52 | 35 | 14 |  |  |
| **Median (IQR)** | -22.9 (-480, 828) | 169 (47.7, 1766) | 290 (57.7, 1920) | 137 (37.7, 1325) |  |  |
| **Range** | (-1122, 2334) | (-903, 2560) | (-903, 2560) | (-121, 2400) |  |  |
| ***p*** | 0.920 | 0.158 | 0.235 | 0.608 |  |  |
| **V8 (N+/post-COVID-19 data excluded)** |  |  |  |  |  |  |
| **n** | 22 | 63 | 39 | 18 | 5 | 1 |
| **Median (IQR)** | 535 (154, 640) | 82.9 (38.6, 160) | 82.9 (38.6, 166) | 113 (57.7, 160) | 77.2 (27.9, 80.0) | 56.6 (56.6, 56.6) |
| **Range** | (20.0, 1280) | (0.0, 1235) | (0.0, 1235) | (0.0, 640) | (20.0, 82.9) | (56.6, 56.6) |
| **V9 (N+/post-COVID-19 data excluded)** |  |  |  |  |  |  |
| **n** | 16 | 53 | 33 | 16 | 3 | 1 |
| **Median (IQR)** | 269 (134, 820) | 309 (154, 1280) | 332 (154, 1808) | 177 (154, 984) | 160 (0.0, 309) | 116 (116, 116) |
| **Range** | (0.0, 2560) | (0.0, 2560) | (0.0, 2560) | (38.6, 2560) | (0.0, 309) | (116, 116) |
| **Change from V8 to V9 (N+/post-COVID-19 data excluded)** |  |  |  |  |  |  |
| **n** | 16 | 51 | 32 | 15 |  |  |
| **Median (IQR)** | -89.5 (-504, 46.0) | 149 (57.7, 948) | 223 (60.6, 1229) | 134 (37.7, 1244) |  |  |
| **Range** | (-1122, 2394) | (-903, 2480) | (-903, 2480) | (-121, 2400) |  |  |
| ***p*** | 0.858 | 0.009 | 0.249 | 0.514 |  |  |
| **V8 (N+/post-COVID-19, out-of-window data excluded)** |  |  |  |  |  |  |
| **n** | 20 | 57 | 37 | 16 | 4 | 0 |
| **Median (IQR)** | 629 (190, 652) | 82.9 (27.9, 160) | 82.9 (38.6, 160) | 134 (39.2, 160) | 52.5 (23.9, 78.6) | – |
| **Range** | (82.9, 1280) | (0.0, 1235) | (0.0, 1235) | (0.0, 640) | (20.0, 80.0) | – |
| **V9 (N+/post-COVID-19, out-of-window data excluded)** |  |  |  |  |  |  |
| **n** | 15 | 52 | 32 | 16 | 3 | 1 |
| **Median (IQR)** | 309 (113, 1280) | 269 (154, 972) | 326 (154, 1544) | 177 (154, 984) | 160 (0.0, 309) | 116 (116, 116) |
| **Range** | (0.0, 2560) | (0.0, 2560) | (0.0, 2560) | (38.6, 2560) | (0.0, 309) | (116, 116) |
| **Change from V8 to V9 (N+/post-COVID-19, out-of-window data excluded)** |  |  |  |  |  |  |
| **n** | 13 | 45 | 29 | 13 |  |  |
| **Median (IQR)** | -149 (-550, 0.0) | 140 (37.7, 612) | 166 (57.7, 643) | 134 (37.7, 582) |  |  |
| **Range** | (-1122, 1237) | (-903, 2480) | (-903, 2480) | (-121, 2400) |  |  |
| ***p*** | 0.552 | 0.017 | 0.187 | 0.590 |  |  |

**Supplemental table S8**. Intergroup/subgroup comparisons of live SARS-CoV-2 50% neutralization titers (difference in median NT50) at 24 weeks (V8) and 48 weeks (V9) post-D1 by quantile regression. Adjusted for age, vaccine regimen for D1 and D2 (mRNA/mRNA, mRNA/ChAdOx1, and ChAdOx1/ChAdOx1), and time lapse between D1 and D2. For comparison at V9, we adjusted for time lapse between D3 and V9.

|  | **All data** | | **Out-of-window data excluded** | | **N+/post-COVID-19 data excluded** | | **N+/post-COVID-19, out-of-window data excluded** | |
| --- | --- | --- | --- | --- | --- | --- | --- | --- |
|  | **Difference in median (95% CI)** | ***p*** | **Difference in median (95% CI)** | ***p*** | **Difference in median (95% CI)** | ***p*** | **Difference in median (95% CI)** | ***p*** |
| **V8** |  |  |  |  |  |  |  |  |
| HIV+ vs HIV– | -515 (-725, -304) | <0.001 | -488 (-704, -271) | <0.001 | -413 (-615, -212) | <0.001 | -488 (-699, -277) | <0.001 |
| IR vs HIV– | -505 (-714, -297) | <0.001 | -494 (-701, -286) | <0.001 | -423 (-662, -183) | <0.001 | -484 (-693, -275) | <0.001 |
| INR vs HIV– | -512 (-723, -302) | <0.001 | -482 (-692, -273) | <0.001 | -430 (-670, -189) | <0.001 | -484 (-695, -274) | <0.001 |
| INR vs IR | -6.9 (-65.4, 51.5) | 0.813 | 11.4 (-50.2, 72.9) | 0.714 | -7.2 (-58.1, 43.8) | 0.780 | -0.6 (-48.0, 46.8) | 0.980 |
| **V9** |  |  |  |  |  |  |  |  |
| HIV+ vs HIV– | 63.0 (-487, 613) | 0.820 | 64.5 (-586, 715) | 0.844 | 49.1 (-252, 350) | 0.745 | 50.5 (-324, 425) | 0.788 |
| IR vs HIV– | 111 (-729, 951) | 0.793 | 103 (-714, 920) | 0.801 | 83.3 (-549, 715) | 0.793 | 40.7 (-506, 587) | 0.882 |
| INR vs HIV– | 12.8 (-956, 981) | 0.979 | 16.5 (-925, 958) | 0.972 | 101 (-605, 806) | 0.776 | 52.3 (-612, 716) | 0.875 |
| INR vs IR | -98.2 (-930, 733) | 0.814 | -86.8 (-889, 715) | 0.830 | 17.2 (-713, 747) | 0.962 | 11.6 (-570, 594) | 0.968 |

**Supplemental table S9**. Changes in frequencies of Spike-specific B cells (per 10^6^ total B cells) in PWH – from baseline (V1) to V8, to V9. No data were post-infection. No data at V1 and V9 were out-of-window. *P* values are based on quantile regression.

| **Among HIV+** | **RBD+S1+ B cells** | **NTD+S1+ B Cells** |
| --- | --- | --- |
| **V1** |  |  |
| n | 17 | 17 |
| Median (IQR) | 0.0 (0.0, 0.0) | 0.0 (0.0, 9.2) |
| Range | (0.0, 0.0) | (0.0, 85.3) |
| **V8** |  |  |
| n | 16 | 16 |
| Median (IQR) | 73.6 (0.0, 363) | 62.6 (0.0, 273) |
| Range | (0.0, 738) | (0.0, 993) |
| **V9** |  |  |
| n | 17 | 17 |
| Median (IQR) | 480 (118, 676) | 200 (118, 409) |
| Range | (0.0, 1710) | (0.0, 1020) |
| **Change from V1 to V8** |  |  |
| n | 16 | 16 |
| Median (IQR) | 73.6 (0.0, 363) | 62.6 (0.0, 258) |
| Range | (0.0, 738) | (-42.2, 984) |
| *p* | 0.415 | 0.356 |
| **Change from V1 to V9** |  |  |
| n | 17 | 17 |
| Median (IQR) | 480 (118, 676) | 200 (115, 409) |
| Range | (0.0, 1710) | (0.0, 1011) |
| *p* | 0.008 | 0.034 |
| **Change from V8 to V9** |  |  |
| n | 16 | 16 |
| Median (IQR) | 266 (92.7, 505) | 125 (50.8, 199) |
| Range | (-15.4, 1339) | (-81.4, 546) |
| *p* | 0.170 | 0.036 |
| **V8** (Out-of-window data excluded) |  |  |
| n | 15 | 15 |
| Median (IQR) | 79.2 (0.0, 423) | 66.8 (0.0, 334) |
| Range | (0.0, 738) | (0.0, 993) |
| **Change from V1 to V8** (Out-of-window data excluded) |  |  |
| n | 15 | 15 |
| Median (IQR) | 79.2 (0.0, 423) | 66.8 (0.0, 304) |
| Range | (0.0, 738) | (-42.2, 984) |
| *p* | 0.371 | 0.303 |
| **Change from V8 to V9** (Out-of-window data excluded) |  |  |
| n | 15 | 15 |
| Median (IQR) | 203 (75.8, 478) | 109 (42.5, 163) |
| Range | (-15.4, 1265) | (-81.4, 501) |
| *p* | 0.104 | 0.011 |

**Supplemental table S10**. Within-group/subgroup changes in concentrations of RBD-specific nAbs (IU/mL) based on rACE2 displacement (rRBD surrogate neutralisation assay, snELISA). All V1 data were below LLOD (lower limit of detection). *P* values are based on quantile regression. N^+^/post-COVID-19 samples (n=16) and V9 samples preceded by D4 (n=3) were excluded.

|  | **HIV–** | **HIV+** | **IRs** | **INRs** | **LLVs** | **LTNPs** |
| --- | --- | --- | --- | --- | --- | --- |
| **V7** |  |  |  |  |  |  |
| n | 9 | 34 | 22 | 8 | 3 | 1 |
| Median (IQR) | 165 (76.5, 226) | 106 (50.1, 377) | 225 (50.1, 472) | 73.0 (50.1, 132) | 50.1 (50.1, 227) | 632 (632, 632) |
| Range | (50.1^a^, 606) | (50.1, 1520^b^) | (50.1, 1520) | (50.1, 1520) | (50.1, 227) | (632, 632) |
| **V8** |  |  |  |  |  |  |
| n | 22 | 37 | 23 | 10 | 3 | 1 |
| Median (IQR) | 50.1 (50.1, 181) | 50.1 (50.1, 65.3) | 50.1 (50.1, 80.8) | 50.1 (50.1, 65.3) | 50.1 (50.1, 50.1) | 314 (314, 314) |
| Range | (50.1, 323) | (50.1, 1069) | (50.1, 517) | (50.1, 1069) | (50.1, 50.1) | (314, 314) |
| **V8b** |  |  |  |  |  |  |
| n | 7 | 36 | 24 | 8 | 3 | 1 |
| Median (IQR) | 670 (211, 1519) | 403 (260, 1130) | 621 (278, 1303) | 315 (239, 414) | 325 (50.1, 926) | 1520 (1520, 1520) |
| Range | (50.1, 1520) | (50.1, 1520) | (50.1, 1520) | (50.1, 1520) | (50.1, 926) | (1520, 1520) |
| **V9** |  |  |  |  |  |  |
| n | 13 | 30 | 19 | 7 | 3 | 1 |
| Median (IQR) | 228 (70.6, 336) | 232 (57.5, 416) | 367 (122, 1209) | 59.9 (50.1, 124) | 60.5 (50.1, 199) | 456 (456, 456) |
| Range | (50.1, 1520) | (50.1, 1520) | (50.1, 1520) | (50.1, 1044) | (50.1, 199) | (456, 456) |
| **V1 to V7 change** |  |  |  |  |  |  |
| n | 9 | 34 | 22 | 8 | 3 | 1 |
| Median (IQR) | 115 (26.4, 176) | 56.0 (0.0, 327) | 174 (0.0, 422) | 22.9 (0.0, 81.5) | 0.0 (0.0, 177) | 582 (582, 582) |
| Range | (0.0, 555) | (0.0, 1470) | (0.0, 1470) | (0.0, 1470) | (0.0, 177) | (582, 582) |
| *p* | 0.046 | 0.493 | 0.610 | 0.989 |  |  |
| **V7 to V8 change** |  |  |  |  |  |  |
| n | 9 | 30 | 19 | 7 | 3 | 1 |
| Median (IQR) | -89.9 (-142, -15.0) | -37.0 (-255, 0.0) | -49.4 (-282, 0.0) | -1.0 (-101, 0.0) | 0.0 (-177, 0.0) | -318 (-318, -318) |
| Range | (-419, 0.0) | (-1003, 0.0) | (-1003, 0.0) | (-451, 0.0) | (-177, 0.0) | (-318, -318) |
| *p* | 0.064 | 0.554 | 0.430 | 0.988 | - | - |
| **V8 to V8b change** |  |  |  |  |  |  |
| n | 7 | 31 | 20 | 7 | 3 | 1 |
| Median (IQR) | 457 (30.4, 1334) | 301 (160, 876) | 437 (157, 910) | 277 (160, 423) | 275 (0.0, 876) | 1206 (1206, 1206) |
| Range | (0.0, 1469) | (0.0, 1467) | (0.0, 1467) | (0.0, 451) | (0.0, 876) | (1206, 1206) |
| *p* | 0.239 | 0.007 | 0.043 | 0.005 | - | - |
| **V8b to V9 change** |  |  |  |  |  |  |
| n | 5 | 28 | 18 | 6 | 3 | 1 |
| Median (IQR) | 0.0 (-715, 126) | -191 (-510, 0.0) | -97.5 (-544, 0.0) | -230 (-349, -165) | -265 (-727, 0.0) | -1064 (-1064, -1064) |
| Range | (-1151, 1208) | (-1104, 1208) | (-1104, 1208) | (-476, 0.0) | (-727, 0.0) | (-1064, -1064) |
| *p* | 1.000 | 0.043 | 0.628 | 0.051 | - | - |
| **V8 to V9 change** |  |  |  |  |  |  |
| n | 13 | 26 | 15 | 7 | 3 | 1 |
| Median (IQR) | 78.9 (0.0, 183) | 58.9 (0.0, 317) | 281 (0.0, 324) | 0.0 (-5.4, 43.5) | 10.4 (0.0, 149) | 142 (142, 142) |
| Range | (-23.6, 1330) | (-247, 1387) | (-247, 1387) | (-25.2, 74.3) | (0.0, 149) | (142, 142) |
| *p* | 0.123 | 0.527 | 0.023 | 1.000 | - | - |

^a^50.1 IU/mL is the LLOD (as the lower limit of the linear range). The positivity threshold is 54.8 IU/mL.

^b^The values are capped at 1,520 IU/mL which is the ULOQ (the upper limit of the linear range).

**Supplemental table S11**. Changes in concentrations of RBD-specific nAbs (IU/mL) between different groups and subgroups based on rACE2 displacement (rRBD surrogate neutralisation assay, snELISA), by quantile regression. Adjusted for age, vaccine regimen for D1 and D2 (mRNA/mRNA, mRNA/ChAdOx1, and ChAdOx1/ChAdOx1) and time lapse between D1 and D2. We further adjusted for time lapse between D3 and V9 for comparison at V9.

|  | **All data** | | **Out-of-window data excluded** | | **N+/post-COVID-19 data excluded** | | **N+/post-COVID-19 and out-of-window data excluded** | |
| --- | --- | --- | --- | --- | --- | --- | --- | --- |
|  | **Estimated difference in median (95% CI)** | ***p*** | **Estimated difference in median (95% CI)** | ***p*** | **Estimated difference in median (95% CI)** | ***p*** | **Estimated difference in median (95% CI)** | ***p*** |
| **Comparison at V7** |  |  |  |  |  |  |  |  |
| HIV+ vs HIV– | 21.0 (-182, 224) | 0.836 | 21.0 (-182, 224) | 0.836 | 61.7 (-140, 263) | 0.539 | 61.7 (-140, 263) | 0.539 |
| IR vs HIV– | 53.3 (-171, 277) | 0.632 | 53.3 (-171, 277) | 0.632 | 113 (-147, 373) | 0.382 | 113 (-147, 373) | 0.382 |
| INR vs HIV– | -4.3 (-273, 265) | 0.974 | -4.3 (-273, 265) | 0.974 | 26.3 (-278, 330) | 0.862 | 26.3 (-278, 330) | 0.862 |
| INR vs IR | -57.6 (-311, 195) | 0.647 | -57.6 (-311, 195) | 0.647 | -86.9 (-335, 161) | 0.481 | -86.9 (-335, 161) | 0.481 |
| **Comparison at V8** |  |  |  |  |  |  |  |  |
| HIV+ vs HIV– | 0.0 (-17.7, 17.7) | 1.000 | 0.0 (-11.2, 11.2) | 1.000 | 0.0 (-18.6, 18.6) | 1.000 | 0.0 (-13.2, 13.2) | 1.000 |
| IR vs HIV– | 0.0 (-17.6, 17.6) | 1.000 | -0.5 (-13.6, 12.6) | 0.938 | 0.0 (-20.5, 20.5) | 1.000 | -0.5 (-16.7, 15.7) | 0.953 |
| INR vs HIV– | 0.0 (-25.1, 25.1) | 1.000 | -0.3 (-18.5, 17.9) | 0.971 | 0.0 (-27.9, 27.9) | 1.000 | -0.3 (-22.5, 21.9) | 0.979 |
| INR vs IR | 0.0 (-20.5, 20.5) | 1.000 | 0.2 (-15.2, 15.6) | 0.981 | 0.0 (-24.8, 24.8) | 1.000 | 0.2 (-17.7, 18.1) | 0.983 |
| **Comparison at V8b** |  |  |  |  |  |  |  |  |
| HIV+ vs HIV– | -282 (-1011, 448) | 0.440 | -282 (-1011, 448) | 0.440 | -72.1 (-717, 573) | 0.822 | -72.1 (-717, 573) | 0.822 |
| IR vs HIV– | -137 (-793, 519) | 0.674 | -137 (-793, 519) | 0.674 | 9.3 (-619, 638) | 0.976 | 9.3 (-619, 638) | 0.976 |
| INR vs HIV– | -680 (-1425, 65.5) | 0.073 | -680 (-1425, 65.5) | 0.073 | -450 (-1216, 316) | 0.240 | -450 (-1216, 316) | 0.240 |
| INR vs IR | -543 (-1087, 1.8) | 0.051 | -543 (-1087, 1.8) | 0.051 | -459 (-991, 72.0) | 0.088 | -459 (-991, 72.0) | 0.088 |
| **Comparison at V9** |  |  |  |  |  |  |  |  |
| HIV+ vs HIV– | 23.3 (-255, 301) | 0.867 | 23.3 (-255, 301) | 0.867 | 53.0 (-118, 224) | 0.533 | 53.0 (-118, 224) | 0.533 |
| IR vs HIV– | 107 (-338, 551) | 0.630 | 107 (-338, 551) | 0.630 | 111 (-121, 344) | 0.336 | 111 (-121, 344) | 0.336 |
| INR vs HIV– | 13.1 (-580, 607) | 0.965 | 13.1 (-580, 607) | 0.965 | -4.7 (-276, 267) | 0.972 | -4.7 (-276, 267) | 0.972 |
| INR vs IR | -93.8 (-713, 525) | 0.761 | -93.8 (-713, 525) | 0.761 | -116 (-464, 232) | 0.501 | -116 (-464, 232) | 0.501 |

**Supplemental table S12A**. Intergroup/subgroup comparisons for spike IgG levels in saliva (%AUC) at different time points. *P* values are based on Wilcoxon rank sum test.

|  | **HIV–** | **HIV+** | ***p*^a^** | **IRs** | **INRs** | ***p*^b^** | ***p*^c^** | ***p*^d^** |
| --- | --- | --- | --- | --- | --- | --- | --- | --- |
| **All samples** |  |  |  |  |  |  |  |  |
| V1, median (IQR) | 0.65 (0.25, 1.55) | 0.15 (0.00, 0.38) | 0.025 | 0.15 (0.00, 0.35) | 0.05 (0.00, 0.20) | 0.040 | 0.020 | 0.534 |
| n | 7 | 23 |  | 14 | 7 |  |  |  |
| V4, median (IQR) | 3.61 (1.25, 11.9) | 7.99 (2.88, 10.8) | 0.326 | 6.07 (2.88, 10.3) | 9.72 (8.34, 18.5) | 0.412 | 0.153 | 0.216 |
| n | 7 | 21 |  | 14 | 6 |  |  |  |
| V5, median (IQR) | 32.2 (17.9, 51.2) | 23.2 (8.32, 48.9) | 0.333 | 24.3 (10.7, 48.9) | 29.6 (5.76, 125) | 0.552 | 0.565 | 0.843 |
| n | 7 | 22 |  | 13 | 7 |  |  |  |
| V8, median (IQR) | 22.9 (6.16, 62.6) | 22.0 (9.47, 60.2) | 0.962 | 22.3 (8.59, 59.4) | 21.6 (13.5, 73.0) | 0.860 | 0.749 | 0.459 |
| n | 7 | 24 |  | 15 | 7 |  |  |  |
| V8b, median (IQR) | 157 (106, 181) | 119 (67.6, 163) | 0.178 | 111 (67.6, 168) | 119 (85.9, 161) | 0.161 | 0.475 | 0.881 |
| n | 6 | 23 |  | 14 | 7 |  |  |  |
| V9, median (IQR) | 109 (91.6, 120) | 48.1 (21.8, 98) | 0.145 | 48.0 (34.6, 129) | 65.8 (21.8, 88.2) | 0.458 | 0.055 | 0.934 |
| n | 6 | 22 |  | 14 | 6 |  |  |  |
| **Out-of-window data excluded** |  |  |  |  |  |  |  |  |
| V1, median (IQR) | 0.65 (0.25, 1.55) | 0.15 (0.00, 0.38) | 0.025 | 0.15 (0.00, 0.35) | 0.05 (0.00, 0.20) | 0.040 | 0.020 | 0.534 |
| n | 7 | 23 |  | 14 | 7 |  |  |  |
| V4, median (IQR) | 3.61 (1.25, 11.9) | 8.17 (3.73, 11.5) | 0.268 | 6.07 (2.88, 10.3) | 10.8 (8.69, 18.5) | 0.412 | 0.062 | 0.064 |
| n | 7 | 20 |  | 14 | 5 |  |  |  |
| V5, median (IQR) | 32.2 (17.9, 51.2) | 23.2 (8.32, 48.9) | 0.333 | 24.3 (10.7, 48.9) | 29.6 (5.76, 125) | 0.552 | 0.565 | 0.843 |
| n | 7 | 22 |  | 13 | 7 |  |  |  |
| V8, median (IQR) | 25.5 (6.16, 62.6) | 18.7 (8.59, 56.4) | 0.799 | 22.3 (8.59, 56.4) | 18.7 (13.5, 43.3) | 0.792 | 0.855 | 0.588 |
| n | 6 | 19 |  | 13 | 5 |  |  |  |
| V8b, median (IQR) | 157 (106, 181) | 119 (67.6, 163) | 0.178 | 111 (67.6, 168) | 119 (85.9, 161) | 0.161 | 0.475 | 0.881 |
| n | 6 | 23 |  | 14 | 7 |  |  |  |
| V9, median (IQR) | 109 (91.6, 120) | 48.1 (21.8, 97.7) | 0.145 | 48.0 (34.6, 129) | 65.8 (21.8, 88.2) | 0.458 | 0.055 | 0.934 |
| n | 6 | 22 |  | 14 | 6 |  |  |  |
| **N+/post-COVID-19 data excluded** |  |  |  |  |  |  |  |  |
| V1, median (IQR) | 0.65 (0.25, 1.55) | 0.15 (0.00, 0.38) | 0.025 | 0.15 (0.00, 0.35) | 0.05 (0.00, 0.20) | 0.040 | 0.020 | 0.534 |
| n | 7 | 23 |  | 14 | 7 |  |  |  |
| V4, median (IQR) | 3.61 (1.25, 11.9) | 7.99 (2.88, 10.8) | 0.326 | 6.07 (2.88, 10.3) | 9.72 (8.34, 18.5) | 0.412 | 0.153 | 0.216 |
| n | 7 | 21 |  | 14 | 6 |  |  |  |
| V5, median (IQR) | 32.2 (17.9, 51.2) | 23.2 (8.32, 48.9) | 0.333 | 24.3 (10.7, 48.9) | 29.6 (5.76, 125) | 0.552 | 0.565 | 0.843 |
| n | 7 | 22 |  | 13 | 7 |  |  |  |
| V8, median (IQR) | 22.9 (6.16, 62.6) | 21.6 (8.82, 60.9) | 0.980 | 27.0 (8.59, 59.4) | 21.6 (13.5, 73.0) | 0.881 | 0.749 | 0.412 |
| n | 7 | 23 |  | 14 | 7 |  |  |  |
| V8b, median (IQR) | 136 (106, 179) | 119 (67.6, 163) | 0.384 | 111 (67.6, 168) | 119 (85.9, 161) | 0.355 | 0.685 | 0.881 |
| n | 5 | 23 |  | 14 | 7 |  |  |  |
| V9, median (IQR) | 95.9 (56.2, 110) | 48.1 (21.8, 97.7) | 0.434 | 48.0 (34.6, 129) | 65.8 (21.8, 88.2) | 0.832 | 0.201 | 0.934 |
| n | 4 | 22 |  | 14 | 6 |  |  |  |
| **N+/post-COVID-19 and out-of-window data excluded** |  |  |  |  |  |  |  |  |
| V1, median (IQR) | 0.65 (0.25, 1.55) | 0.15 (0.00, 0.38) | 0.025 | 0.15 (0.00, 0.35) | 0.05 (0.00, 0.20) | 0.040 | 0.020 | 0.534 |
| n | 7 | 23 |  | 14 | 7 |  |  |  |
| V4, median (IQR) | 3.61 (1.25, 11.9) | 8.17 (3.73, 11.5) | 0.268 | 6.07 (2.88, 10.3) | 10.8 (8.69, 18.5) | 0.412 | 0.062 | 0.064 |
| n | 7 | 20 |  | 14 | 5 |  |  |  |
| V5, median (IQR) | 32.2 (17.9, 51.2) | 23.2 (8.32, 48.9) | 0.333 | 24.3 (10.7, 48.9) | 29.6 (5.76, 126) | 0.552 | 0.565 | 0.843 |
| n | 7 | 22 |  | 13 | 7 |  |  |  |
| V8, median (IQR) | 25.5 (6.16, 62.6) | 16.1 (8.59, 56.4) | 0.790 | 26.9 (7.74, 57.9) | 18.7 (13.5, 43.3) | 0.779 | 0.855 | 0.527 |
| n | 6 | 18 |  | 12 | 5 |  |  |  |
| V8b, median (IQR) | 136 (106, 179) | 119 (67.6, 163) | 0.384 | 111 (67.6, 168) | 119 (85.9, 161) | 0.355 | 0.685 | 0.881 |
| n | 5 | 23 |  | 14 | 7 |  |  |  |
| V9, median (IQR) | 95.9 (56.2, 110) | 48.1 (21.8, 97.7) | 0.434 | 48.0 (34.6, 129) | 65.8 (21.8, 88.2) | 0.832 | 0.201 | 0.934 |
| n | 4 | 22 |  | 14 | 6 |  |  |  |

^a^Comparison between HIV^–^ and HIV^+^.

^b^Comparison between HIV^–^ and IRs.

^c^Comparison between HIV^–^ and INRs.

^d^Comparison between IRs and INRs.

**Supplemental table S12B**. Intergroup/subgroup comparisons for spike IgA levels in saliva (%AUC) at different time points. *P* value is based on Wilcoxon rank sum test.

|  | **HIV–** | **HIV+** | ***p*^a^** | **IRs** | **INRs** | ***p*^b^** | ***p*^c^** | ***p*^d^** |
| --- | --- | --- | --- | --- | --- | --- | --- | --- |
| **All samples** |  |  |  |  |  |  |  |  |
| V1, median (IQR) | 17.1 (5.91, 21.1) | 1.42 (0.29, 6.20) | 0.011 | 3.17 (0.29, 8.69) | 1.42 (0.24, 2.93) | 0.086 | 0.006 | 0.549 |
| n | 7 | 23 |  | 14 | 7 |  |  |  |
| V4, median (IQR) | 11 (0.34, 14.5) | 5.52 (1.71, 11.8) | 1.000 | 6.88 (1.71, 19.9) | 2.95 (1.51, 10.7) | 0.794 | 0.668 | 0.458 |
| n | 7 | 21 |  | 14 | 6 |  |  |  |
| V5, median (IQR) | 17.9 (15.2, 38.1) | 6.52 (2.10, 22.5) | 0.251 | 7.03 (5.66, 37.9) | 2.73 (1.32, 17.5) | 0.552 | 0.224 | 0.250 |
| n | 7 | 22 |  | 13 | 7 |  |  |  |
| V8, median (IQR) | 10.3 (2.20, 34.0) | 5.62 (0.90, 14.9) | 0.237 | 7.37 (1.12, 19.1) | 1.56 (0.34, 1.81) | 0.622 | 0.035 | 0.053 |
| n | 7 | 24 |  | 15 | 7 |  |  |  |
| V8b, median (IQR) | 16.4 (10.6, 22.6) | 7.62 (2.29, 15.0) | 0.090 | 10.0 (5.62, 32.6) | 4.64 (1.42, 7.86) | 0.302 | 0.022 | 0.156 |
| n | 6 | 23 |  | 14 | 7 |  |  |  |
| V9, median (IQR) | 24.6 (17.3, 30.5) | 3.83 (0.49, 10.2) | 0.008 | 5.88 (1.71, 16.4) | 2.12 (0.20, 4.39) | 0.058 | 0.004 | 0.149 |
| n | 6 | 22 |  | 14 | 6 |  |  |  |
| **Out-of-window data excluded** |  |  |  |  |  |  |  |  |
| V1, median (IQR) | 17.1 (5.91, 21.1) | 1.42 (0.29, 6.20) | 0.011 | 3.17 (0.29, 8.69) | 1.42 (0.24, 2.93) | 0.086 | 0.006 | 0.549 |
| n | 7 | 23 |  | 14 | 7 |  |  |  |
| V4, median (IQR) | 11.0 (0.34, 14.5) | 6.10 (2.10, 13.9) | 0.890 | 6.88 (1.71, 19.9) | 3.08 (2.83, 10.7) | 0.794 | 0.935 | 0.781 |
| n | 7 | 20 |  | 14 | 5 |  |  |  |
| V5, median (IQR) | 17.9 (15.2, 38.1) | 6.52 (2.10, 22.5) | 0.251 | 7.03 (5.66, 37.9) | 2.73 (1.32, 17.5) | 0.552 | 0.224 | 0.250 |
| n | 7 | 22 |  | 13 | 7 |  |  |  |
| V8, median (IQR) | 9.01 (2.20, 34.0) | 5.57 (0.34, 17.5) | 0.279 | 7.37 (5.57, 19.1) | 1.07 (0.34, 1.56) | 0.693 | 0.068 | 0.043 |
| n | 6 | 19 |  | 13 | 5 |  |  |  |
| V8b, median (IQR) | 16.4 (10.6, 22.6) | 7.62 (2.29, 15.0) | 0.090 | 10.0 (5.62, 32.6) | 4.64 (1.42, 7.86) | 0.302 | 0.022 | 0.156 |
| n | 6 | 23 |  | 14 | 7 |  |  |  |
| V9, median (IQR) | 24.6 (17.3, 30.5) | 3.83 (0.49, 10.2) | 0.008 | 5.88 (1.71, 16.4) | 2.12 (0.20, 4.39) | 0.058 | 0.004 | 0.149 |
| n | 6 | 22 |  | 14 | 6 |  |  |  |
| **N+/post-COVID-19 data excluded** |  |  |  |  |  |  |  |  |
| V1, median (IQR) | 17.1 (5.91, 21.1) | 1.42 (0.29, 6.20) | 0.011 | 3.17 (0.29, 8.69) | 1.42 (0.24, 2.93) | 0.086 | 0.006 | 0.549 |
| n | 7 | 23 |  | 14 | 7 |  |  |  |
| V4, median (IQR) | 11.0 (0.34, 14.5) | 5.52 (1.71, 11.8) | 1.000 | 6.88 (1.71, 19.9) | 2.95 (1.51, 10.7) | 0.794 | 0.668 | 0.458 |
| n | 7 | 21 |  | 14 | 6 |  |  |  |
| V5, median (IQR) | 17.9 (15.2, 38.1) | 6.52 (2.10, 22.5) | 0.251 | 7.03 (5.66, 37.9) | 2.73 (1.32, 17.5) | 0.552 | 0.224 | 0.250 |
| n | 7 | 22 |  | 13 | 7 |  |  |  |
| V8, median (IQR) | 10.3 (2.20, 34.0) | 5.57 (0.73, 15.7) | 0.229 | 6.59 (1.12, 19.1) | 1.56 (0.34, 1.81) | 0.628 | 0.035 | 0.073 |
| n | 7 | 23 |  | 14 | 7 |  |  |  |
| V8b, median (IQR) | 14.2 (10.6, 18.6) | 7.62 (2.29, 15.0) | 0.208 | 10.0 (5.62, 32.6) | 4.64 (1.42, 7.86) | 0.547 | 0.042 | 0.156 |
| n | 5 | 23 |  | 14 | 7 |  |  |  |
| V9, median (IQR) | 20.5 (13.2, 24.6) | 3.83 (0.49, 10.2) | 0.039 | 5.88 (1.71, 16.4) | 2.12 (0.20, 4.39) | 0.167 | 0.011 | 0.149 |
| n | 4 | 22 |  | 14 | 6 |  |  |  |
| **N+/post-COVID-19 and out-of-window data excluded** |  |  |  |  |  |  |  |  |
| V1, median (IQR) | 17.1 (5.91, 21.1) | 1.42 (0.29, 6.20) | 0.011 | 3.17 (0.29, 8.69) | 1.42 (0.24, 2.93) | 0.086 | 0.006 | 0.549 |
| n | 7 | 23 |  | 14 | 7 |  |  |  |
| V4, median (IQR) | 11.0 (0.34, 14.5) | 6.10 (2.10, 13.9) | 0.890 | 6.88 (1.71, 19.9) | 3.08 (2.83, 10.7) | 0.794 | 0.935 | 0.781 |
| n | 7 | 20 |  | 14 | 5 |  |  |  |
| V5, median (IQR) | 17.9 (15.2, 38.1) | 6.52 (2.10, 22.5) | 0.251 | 7.03 (5.66, 37.9) | 2.73 (1.32, 17.5) | 0.552 | 0.224 | 0.250 |
| n | 7 | 22 |  | 13 | 7 |  |  |  |
| V8, median (IQR) | 9.01 (2.20, 34.0) | 3.69 (0.34, 17.5) | 0.257 | 6.59 (3.15, 19.4) | 1.07 (0.34, 1.56) | 0.673 | 0.068 | 0.058 |
| n | 6 | 18 |  | 12 | 5 |  |  |  |
| V8b, median (IQR) | 14.2 (10.6, 18.6) | 7.62 (2.29, 15.0) | 0.208 | 10.0 (5.62, 32.6) | 4.64 (1.42, 7.86) | 0.547 | 0.042 | 0.156 |
| n | 5 | 23 |  | 14 | 7 |  |  |  |
| V9, median (IQR) | 20.5 (13.2, 24.6) | 3.83 (0.49, 10.2) | 0.039 | 5.88 (1.71, 16.4) | 2.12 (0.20, 4.39) | 0.167 | 0.011 | 0.149 |
| n | 4 | 22 |  | 14 | 6 |  |  |  |

^a^Comparison between HIV^–^ and HIV^+^.

^b^Comparison between HIV^–^ and IRs.

^c^Comparison between HIV^–^ and INRs.

^d^Comparison between IRs and INRs.

**Supplemental table S13A**. Within-group/subgroup changes in spike IgG levels in saliva (%AUC) from baseline and between neighbouring time points. *P* values are based on quantile regression.

| **IgG spike** | **All data** | | **Out-of-window data excluded** | | **N+/post-COVID-19 data excluded** | | **N+/post-COVID-19 and out-of-window data excluded** | |
| --- | --- | --- | --- | --- | --- | --- | --- | --- |
|  | **Median change (95% CI)** | ***p*** | **Median change (95% CI)** | ***p*** | **Median change (95% CI)** | ***p*** | **Median change (95% CI)** | ***p*** |
| **V4 vs V1** |  |  |  |  |  |  |  |  |
| Within HIV– | 1.93 (-5.09, 8.95) | 0.578 | 1.93 (-5.10, 8.96) | 0.578 | 1.93 (-5.09, 8.95) | 0.578 | 1.93 (-5.10, 8.96) | 0.578 |
| Within HIV+ | 6.59 (2.46, 10.7) | 0.003 | 7.21 (3.23, 11.2) | 0.001 | 6.59 (2.46, 10.7) | 0.003 | 7.21 (3.23, 11.2) | 0.001 |
| Within IR | 4.73 (0.94, 8.53) | 0.017 | 4.73 (1.09, 8.38) | 0.013 | 4.73 (0.94, 8.53) | 0.017 | 4.73 (1.09, 8.38) | 0.013 |
| Within INR | 8.69 (-1.33, 18.7) | 0.086 | 10.7 (0.22, 21.1) | 0.046 | 8.69 (-1.33, 18.7) | 0.086 | 10.7 (0.22, 21.1) | 0.046 |
| **V5 vs V1** |  |  |  |  |  |  |  |  |
| Within HIV– | 31.4 (-7.87, 70.7) | 0.112 | 31.4 (-7.87, 70.7) | 0.112 | 31.4 (-7.87, 70.7) | 0.112 | 31.4 (-7.87, 70.7) | 0.112 |
| Within HIV+ | 21.9 (3.02, 40.7) | 0.025 | 21.9 (3.02, 40.7) | 0.025 | 21.9 (3.02, 40.7) | 0.025 | 21.9 (3.02, 40.7) | 0.025 |
| Within IR | 21.9 (2.28, 41.5) | 0.030 | 21.9 (2.28, 41.5) | 0.030 | 21.9 (2.28, 41.5) | 0.030 | 21.9 (2.28, 41.5) | 0.030 |
| Within INR | 29.6 (-34.7, 94.0) | 0.351 | 29.6 (-34.7, 94.0) | 0.351 | 29.6 (-34.7, 94.0) | 0.351 | 29.6 (-34.7, 94.0) | 0.351 |
| **V8 vs V1** |  |  |  |  |  |  |  |  |
| Within HIV– | 21.3 (-11.9, 54.6) | 0.200 | 16.9 (-28.6, 62.4) | 0.448 | 21.3 (-10.4, 53.0) | 0.179 | 16.9 (-27.1, 61.0) | 0.433 |
| Within HIV+ | 21.6 (-1.59, 44.9) | 0.067 | 13.5 (-6.83, 33.8) | 0.183 | 18.4 (-9.34, 46.1) | 0.185 | 13.5 (-12.1, 39.0) | 0.286 |
| Within IR | 10.1 (-19.6, 39.8) | 0.489 | 10.1 (-21.8, 42.0) | 0.516 | 10.1 (-24.9, 45.1) | 0.556 | 10.1 (-23.9, 44.1) | 0.541 |
| Within INR | 21.6 (-14.5, 57.8) | 0.229 | 18.4 (-22.9, 59.6) | 0.364 | 21.6 (-10.9, 54.2) | 0.182 | 18.4 (-25.3, 62.0) | 0.390 |
| **V8b vs V1** |  |  |  |  |  |  |  |  |
| Within HIV– | 134 (48.2, 220) | 0.004 | 134 (48.2, 220) | 0.004 | 134 (46.5, 222) | 0.004 | 134 (46.5, 222) | 0.004 |
| Within HIV+ | 109 (64.0, 154) | <0.001 | 109 (64.0, 154) | <0.001 | 109 (63.7, 155) | <0.001 | 109 (63.7, 154) | <0.001 |
| Within IR | 94.9 (31.8, 158) | 0.005 | 94.9 (31.8, 158) | 0.005 | 94.9 (33.8, 156) | 0.004 | 94.9 (33.8, 156) | 0.004 |
| Within INR | 119 (66.2, 171) | <0.001 | 119 (66.2, 171) | <0.001 | 119 (61.3, 176) | <0.001 | 119 (61.3, 176) | <0.001 |
| **V9 vs V1** |  |  |  |  |  |  |  |  |
| Within HIV– | 100 (42.2, 158) | 0.002 | 100 (42.2, 158) | 0.002 | 91.4 (16.7, 166) | 0.019 | 91.4 (16.7, 166) | 0.019 |
| Within HIV+ | 46.4 (25.7, 67.1) | <0.001 | 46.4 (25.7, 67.1) | <0.001 | 46.4 (27.8, 65.0) | <0.001 | 46.4 (27.8, 65.0) | <0.001 |
| Within IR | 46.4 (11.2, 81.6) | 0.012 | 46.4 (11.2, 81.6) | 0.012 | 46.4 (8.19, 84.6) | 0.020 | 46.4 (8.19, 84.6) | 0.020 |
| Within INR | 59.3 (5.58, 113) | 0.032 | 59.3 (5.58, 113) | 0.032 | 59.3 (4.44, 114) | 0.036 | 59.3 (4.44, 114) | 0.036 |
| **V5 vs V4** |  |  |  |  |  |  |  |  |
| Within HIV– | 28.6 (-18.5, 75.7) | 0.222 | 28.6 (0.90, 56.3) | 0.044 | 28.6 (-18.5, 75.7) | 0.222 | 28.6 (0.90, 56.3) | 0.044 |
| Within HIV+ | 15.3 (1.10, 29.5) | 0.036 | 15.3 (-2.28, 32.8) | 0.085 | 15.3 (1.10, 29.5) | 0.036 | 15.3 (-2.28, 32.8) | 0.085 |
| Within IR | 13.7 (-3.68, 30.1) | 0.117 | 13.7 (-2.69, 30.0) | 0.097 | 13.7 (-3.68, 31.0) | 0.117 | 13.7 (-2.69, 30.0) | 0.097 |
| Within INR | 21.3 (-53.7, 96.3) | 0.562 | 23.2 (-62.1, 108) | 0.578 | 21.3 (-53.7, 96.3) | 0.562 | 23.2 (-62.1, 108) | 0.578 |
| **V8 vs V5** |  |  |  |  |  |  |  |  |
| Within HIV– | -11.8 (-60.1, 36.6) | 0.621 | -9.55 (-66.4, 47.3) | 0.730 | -11.8 (-60.1, 36.6) | 0.621 | -9.55 (-66.4, 47.3) | 0.730 |
| Within HIV+ | -1.80 (-17.2, 13.6) | 0.812 | -13.1 (-24.9, -1.32) | 0.031 | -1.80 (-17.2, 13.6) | 0.812 | -13.1 (-24.9, -1.32) | 0.031 |
| Within IR | -6.1 (-27.1, 14.9) | 0.553 | -13.5 (-47.3, 20.3) | 0.415 | -6.11 (-27.1, 14.9) | 0.553 | -13.5 (-47.3, 20.4) | 0.415 |
| Within INR | 11.5 (-22.9, 45.9) | 0.497 | -13.1 (-115, 88.7) | 0.790 | 11.5 (-22.9, 45.9) | 0.497 | -13.1 (-115, 88.7) | 0.790 |
| **V8b vs V8** |  |  |  |  |  |  |  |  |
| Within HIV– | 88.4 (6.82, 170) | 0.035 | 88.4 (3.41, 173) | 0.042 | 88.4 (13.5, 163) | 0.023 | 51.1 (-24.3, 127) | 0.173 |
| Within HIV+ | 72.6 (26.6, 119) | 0.003 | 72.6 (17.4, 128) | 0.012 | 72.6 (25.5, 120) | 0.004 | 72.6 (15.9, 129) | 0.015 |
| Within IR | 72.6 (14.2, 131) | 0.017 | 76.8 (19.1, 135) | 0.012 | 72.6 (14.4, 131) | 0.017 | 76.8 (25.0, 129) | 0.006 |
| Within INR | 65.9 (-4.08, 136) | 0.064 | 65.9 (-24.2, 156) | 0.142 | 65.9 (-5.01, 137) | 0.067 | 65.9 (-19.9, 152) | 0.124 |
| **V9 vs V8b** |  |  |  |  |  |  |  |  |
| Within HIV– | -18.5 (-75.4, 38.5) | 0.511 | -18.5 (-75.4, 38.5) | 0.511 | -18.5 (-83.9, 47.0) | 0.566 | -18.5 (-83.9, 47.0) | 0.566 |
| Within HIV+ | -37.3 (-69.8, -4.73) | 0.026 | -37.3 (-69.8, -4.73) | 0.026 | -37.3 (-68.9, -5.63) | 0.023 | -37.3 (-68.9, -5.63) | 0.023 |
| Within IR | -31.7 (-65.1, 1.74) | 0.062 | -31.7 (-65.1, 1.74) | 0.062 | -31.7 (-64.2, 0.82) | 0.056 | -31.7 (-64.2, 0.82) | 0.056 |
| Within INR | -49.6 (-109, 9.73) | 0.097 | -49.6 (-109, 9.73) | 0.097 | -49.6 (-106, 6.33) | 0.079 | -49.6 (-106, 6.33) | 0.079 |

**Supplemental table S13B**. Within-group/subgroup changes in spike IgA levels in saliva (%AUC) from baseline and between neighbouring time points. *P* value is based on quantile regression.

| **IgA spike** | **All data** | | **Out-of-window data excluded** | | **N+/post-COVID-19 data excluded** | | **N+/post-COVID-19 and out-of-window data excluded** | |
| --- | --- | --- | --- | --- | --- | --- | --- | --- |
|  | **Median change (95% CI)** | ***p*** | **Median change (95% CI)** | ***p*** | **Median change (95% CI)** | ***p*** | **Median change (95% CI)** | ***p*** |
| **V4 vs V1** |  |  |  |  |  |  |  |  |
| Within HIV– | -5.37 (-12.6, 1.89) | 0.141 | -5.37 (-12.5, 1.79) | 0.135 | -5.37 (-12.6, 1.89) | 0.141 | -5.37 (-12.5, 1.79) | 0.135 |
| Within HIV+ | 1.71 (-0.57, 3.99) | 0.135 | 1.71 (-0.43, 3.85) | 0.113 | 1.71 (-0.57, 3.99) | 0.135 | 1.71 (-0.43, 3.85) | 0.113 |
| Within IR | 1.27 (-2.64, 5.18) | 0.509 | 1.27 (-1.76, 4.30) | 0.395 | 1.27 (-2.64, 5.18) | 0.509 | 1.27 (-1.76, 4.30) | 0.395 |
| Within INR | 1.71 (-3.46, 6.88) | 0.502 | 3.08 (-2.03, 8.18) | 0.225 | 1.71 (-3.46, 6.88) | 0.502 | 3.08 (-2.03, 8.18) | 0.225 |
| **V5 vs V1** |  |  |  |  |  |  |  |  |
| Within HIV– | 5.18 (-19.0, 29.3) | 0.663 | 5.18 (-19.0, 29.3) | 0.663 | 5.18 (-19.0, 29.3) | 0.663 | 5.18 (-19.0, 29.3) | 0.663 |
| Within HIV+ | 1.32 (-3.57, 6.20) | 0.584 | 1.32 (-3.57, 6.20) | 0.584 | 1.32 (-3.57, 6.20) | 0.584 | 1.32 (-3.57, 6.20) | 0.584 |
| Within IR | 3.37 (-5.08, 11.8) | 0.418 | 3.37 (-5.08, 11.8) | 0.418 | 3.37 (-5.08, 11.8) | 0.418 | 3.37 (-5.08, 11.8) | 0.418 |
| Within INR | 1.32 (-27.0, 29.6) | 0.924 | 1.32 (-27.0, 29.6) | 0.924 | 1.32 (-27.0, 29.6) | 0.924 | 1.32 (-27.0, 29.6) | 0.924 |
| **V8 vs V1** |  |  |  |  |  |  |  |  |
| Within HIV– | -0.15 (-10.8, 10.5) | 0.978 | 0.29 (-11.1, 11.7) | 0.958 | -0.15 (-10.7, 10.4) | 0.978 | 0.29 (-10.6, 11.2) | 0.956 |
| Within HIV+ | 0.20 (-0.92, 1.31) | 0.721 | 0.15 (-1.29, 1.59) | 0.835 | 0.15 (-1.25, 1.55) | 0.832 | 0.15 (-1.61, 1.91) | 0.864 |
| Within IR | 0.54 (-3.53, 4.60) | 0.788 | 0.54 (-6.06, 7.13) | 0.867 | 0.54 (-4.44, 5.52) | 0.826 | 0.54 (-11.4, 12.5) | 0.926 |
| Within INR | -0.20 (-1.74, 1.35) | 0.797 | -0.20 (-2.13, 1.74) | 0.836 | -0.20 (-1.57, 1.18) | 0.773 | -0.20 (-2.22, 1.83) | 0.842 |
| **V8b vs V1** |  |  |  |  |  |  |  |  |
| Within HIV– | 0.88 (-21.9, 23.7) | 0.937 | 0.88 (-21.9, 23.7) | 0.937 | 0.88 (-14.5, 16.3) | 0.907 | 0.88 (-14.5, 16.3) | 0.907 |
| Within HIV+ | 2.49 (-1.65, 6.63) | 0.227 | 2.49 (-1.65, 6.63) | 0.227 | 2.49 (-1.71, 6.69) | 0.233 | 2.49 (-1.71, 6.69) | 0.233 |
| Within IR | 1.95 (-3.92, 7.83) | 0.498 | 1.95 (-3.92, 7.83) | 0.498 | 1.95 (-5.43, 9.33) | 0.589 | 1.95 (-5.43, 9.33) | 0.589 |
| Within INR | 3.22 (-1.41, 7.85) | 0.163 | 3.22 (-1.41, 7.85) | 0.163 | 3.22 (-1.73, 8.18) | 0.191 | 3.22 (-1.73, 8.18) | 0.191 |
| **V9 vs V1** |  |  |  |  |  |  |  |  |
| Within HIV– | 2.64 (-11.5, 16.8) | 0.704 | 2.64 (-11.5, 16.8) | 0.704 | -0.34 (-2.11, 1.43) | 0.694 | -0.34 (-2.11, 1.43) | 0.694 |
| Within HIV+ | 0.44 (-1.98, 2.86) | 0.711 | 0.44 (-1.98, 2.86) | 0.711 | 0.44 (-2.14, 3.02) | 0.728 | 0.44 (-2.14, 3.02) | 0.728 |
| Within IR | 0.49 (-5.48, 6.46) | 0.867 | 0.49 (-5.48, 6.46) | 0.867 | 0.49 (-6.35, 7.33) | 0.883 | 0.49 (-6.35, 7.33) | 0.883 |
| Within INR | 0.44 (-1.85, 2.73) | 0.695 | 0.44 (-1.85, 2.73) | 0.695 | 0.44 (-1.59, 2.47) | 0.656 | 0.44 (-1.59, 2.47) | 0.656 |
| **V5 vs V4** |  |  |  |  |  |  |  |  |
| Within HIV– | 14.8 (-15.3, 44.9) | 0.319 | 14.8 (-9.51, 39.2) | 0.220 | 14.8 (-15.3, 44.9) | 0.319 | 14.8 (-9.51, 39.2) | 0.220 |
| Within HIV+ | 4.44 (0.04, 8.84) | 0.048 | 4.44 (-5.03, 13.9) | 0.342 | 4.44 (0.04, 8.84) | 0.048 | 4.44 (-5.03, 13.9) | 0.342 |
| Within IR | 4.44 (-10.5, 19.3) | 0.542 | 4.44 (-5.87, 14.8) | 0.380 | 4.44 (-10.5, 19.3) | 0.542 | 4.44 (-5.87, 14.8) | 0.380 |
| Within INR | 1.22 (-31.0, 33.5) | 0.938 | 6.79 (-36.0, 49.6) | 0.745 | 1.22 (-31.0, 33.5) | 0.938 | 6.79 (-36.0, 49.6) | 0.745 |
| **V8 vs V5** |  |  |  |  |  |  |  |  |
| Within HIV– | -7.08 (-45.6, 31.5) | 0.709 | -4.88 (-48.2, 38.4) | 0.817 | -7.08 (-45.6, 31.5) | 0.709 | -4.88 (-48.2, 38.4) | 0.817 |
| Within HIV+ | -1.37 (-8.99, 6.26) | 0.716 | -1.46 (-17.0, 14.1) | 0.846 | -1.37 (-8.99, 6.26) | 0.716 | -1.46 (-17.0, 14.1) | 0.846 |
| Within IR | -3.37 (-39.5, 32.8) | 0.849 | -5.96 (-68.6, 56.6) | 0.844 | -3.37 (-39.5, 32.8) | 0.849 | -5.96 (-68.6, 56.6) | 0.844 |
| Within INR | -1.17 (-22.1, 19.7) | 0.909 | -1.17 (-97.3, 95.0) | 0.980 | -1.17 (-22.1, 19.7) | 0.909 | -1.17 (-97.3, 95.0) | 0.980 |
| **V8b vs V8** |  |  |  |  |  |  |  |  |
| Within HIV– | 2.93 (-31.7, 37.5) | 0.863 | 2.93 (-34.4, 40.2) | 0.872 | 2.93 (-25.2, 31.1) | 0.832 | 2.93 (-23.5, 29.4) | 0.820 |
| Within HIV+ | 1.61 (-2.22, 5.44) | 0.395 | 3.08 (-2.02, 8.17) | 0.223 | 1.61 (-2.32, 5.54) | 0.407 | 3.08 (-1.53, 7.68) | 0.179 |
| Within IR | 1.12 (-6.05, 8.30) | 0.750 | 1.61 (-8.12, 11.3) | 0.733 | 1.12 (-5.97, 8.22) | 0.746 | 1.61 (-6.74, 9.97) | 0.690 |
| Within INR | 3.08 (-2.29, 8.44) | 0.248 | 3.08 (-2.21, 8.37) | 0.238 | 3.08 (-2.47, 8.62) | 0.263 | 3.08 (-1.98, 8.13) | 0.217 |
| **V9 vs V8b** |  |  |  |  |  |  |  |  |
| Within HIV– | 1.13 (-17.4, 19.7) | 0.901 | 1.13 (-17.4, 19.7) | 0.901 | 1.13 (-13.3, 15.6) | 0.873 | 1.13 (-13.3, 15.6) | 0.873 |
| Within HIV+ | -3.91 (-6.82, -1.00) | 0.010 | -3.91 (-6.82, -1.00) | 0.010 | -3.91 (-6.63, -1.18) | 0.007 | -3.91 (-6.63, -1.18) | 0.007 |
| Within IR | -3.03 (-7.74, 1.68) | 0.196 | -3.03 (-7.74, 1.68) | 0.196 | -3.03 (-8.59, 2.54) | 0.271 | -3.03 (-8.59, 2.54) | 0.271 |
| Within INR | -4.64 (-11.2, 1.89) | 0.155 | -4.64 (-11.2, 1.89) | 0.155 | -4.64 (-11.3, 2.01) | 0.161 | -4.64 (-11.3, 2.01) | 0.161 |

**Supplemental table S14**. SARS-CoV-2 anti-spike responses in T cells at 24 weeks (V8) and 48 weeks (V9) post-D1 relative to the baseline (V1), by ELISpot and Wilcoxon rank sum test. Formal comparisons involving the LLV and LTNP subgroups were not performed as the sample size was too small.

|  | **HIV–** | **HIV+** | ***p*^a^** | **IRs** | **INRs** | ***p*^b^** | ***p*^c^** | ***p*^d^** |
| --- | --- | --- | --- | --- | --- | --- | --- | --- |
| **IFN-γ spike** |  |  |  |  |  |  |  |  |
| V1, median (IQR) | 0.0 (0.0, 17.5) | 0.0 (0.0, 4.4) | 0.839 | 1.3 (0.0, 7.5) | 0.0 (0.0, 0.0) | 1.000 | 0.400 | 0.146 |
| n | 6 | 28 |  | 18 | 6 |  |  |  |
| V8, median (IQR) | 11.3 (3.8, 12.5) | 12.5 (0.0, 35.0) | 0.526 | 25.0 (10.0, 67.5) | 0.0 (0.0, 12.5) | 0.082 | 0.204 | 0.020 |
| n | 8 | 29 |  | 19 | 6 |  |  |  |
| V9, median (IQR) | 17.5 (6.3, 23.8) | 35.0 (10.0, 70.0) | 0.166 | 55.0 (25.0, 133) | 2.5 (0.0, 5.0) | 0.002 | 0.057 | 0.001 |
| n | 8 | 29 |  | 19 | 6 |  |  |  |
| **IL-2 spike** |  |  |  |  |  |  |  |  |
| V1, median (IQR) | 1.9 (0.0, 2.5) | 0.0 (0.0, 2.5) | 0.571 | 1.3 (0.0, 5.0) | 0.0 (0.0, 2.5) | 0.943 | 0.491 | 0.558 |
| n | 6 | 28 |  | 18 | 6 |  |  |  |
| V8, median (IQR) | 41.3 (23.8, 50.0) | 92.5 (15.0, 158) | 0.196 | 123 (85.0, 268) | 10.6 (0.0, 20.8) | 0.009 | 0.070 | 0.016 |
| n | 8 | 29 |  | 19 | 6 |  |  |  |
| V9, median (IQR) | 43.8 (20.0, 60.0) | 103 (40.0, 190) | 0.027 | 135 (100, 360) | 12.5 (0.0, 37.5) | <0.001 | 0.119 | 0.001 |
| n | 8 | 29 |  | 19 | 6 |  |  |  |
| **Dual spike** |  |  |  |  |  |  |  |  |
| V1, median (IQR) | 0.0 (0.0, 0.0) | 0.0 (0.0, 0.0) | 0.409 | 0.0 (0.0, 0.0) | 0.0 (0.0, 0.0) | 0.296 | 1.000 | 0.296 |
| n | 6 | 28 |  | 18 | 6 |  |  |  |
| V8, median (IQR) | 3.8 (2.5, 6.3) | 2.5 (0.0, 7.5) | 0.790 | 7.5 (2.5, 12.5) | 0.0 (0.0, 7.5) | 0.389 | 0.230 | 0.078 |
| n | 8 | 29 |  | 19 | 6 |  |  |  |
| V9, median (IQR) | 3.8 (2.5, 5.0) | 7.5 (2.5, 22.5) | 0.168 | 15.0 (7.5, 40.0) | 0.0 (0.0, 2.5) | 0.001 | 0.021 | <0.001 |
| n | 8 | 29 |  | 19 | 6 |  |  |  |

^a^Comparisons between HIV^–^ and HIV^+^.

^b^Comparisons between HIV^–^ and IRs.

^c^Comparisons between HIV^–^ and INRs.

^d^Comparisons between IRs and INRs.

**Supplemental table S15**. Within-group median changes between study visits for SARS-CoV-2 spike responses in T cells (ELISpot) following COVID-19 vaccination, by quantile regression.

| **IFN-γ spike** | **All data** | | **Out-of-window data excluded** | |
| --- | --- | --- | --- | --- |
|  | **Median change (95% CI)** | ***p*** | **Median change (95% CI)** | ***p*** |
| **V8 vs V1** |  |  |  |  |
| Within HIV– | 7.50 (-8.99, 24.0) | 0.361 | 7.50 (-9.23, 24.2) | 0.368 |
| Within HIV+ | 10.0 (1.66, 18.3) | 0.020 | 12.5 (4.02, 21.0) | 0.005 |
| Within IR | 15.0 (-1.91, 31.9) | 0.080 | 15.0 (-1.91, 31.9) | 0.080 |
| Within INR | 0.00 (-7.52, 7.52) | 1.000 | 0.00 (-7.52, 7.52) | 1.000 |
| **V9 vs V8** |  |  |  |  |
| Within HIV– | 5.00 (-3.31, 13.3) | 0.230 | 5.00 (-6.64, 16.6) | 0.389 |
| Within HIV+ | 22.5 (1.32, 43.7) | 0.038 | 22.5 (1.58, 43.4) | 0.036 |
| Within IR | 35.0 (14.6, 55.4) | 0.001 | 35.0 (13.1, 56.9) | 0.003 |
| Within INR | 0.00 (-12.1, 12.1) | 1.000 | 0.00 (-12.4, 12.4) | 1.000 |
| **IL-2 spike** | **All data** | | **Out-of-window data excluded** | |
|  | **Median change (95% CI)** | ***p*** | **Median change (95% CI)** | ***p*** |
| **V8 vs V1** |  |  |  |  |
| Within HIV– | 40.0 (26.4, 53.6) | <0.001 | 40.0 (25.5, 54.5) | <0.001 |
| Within HIV+ | 87.5 (28.4, 147) | 0.005 | 87.5 (26.1, 149) | 0.007 |
| Within IR | 103 (55.7, 149) | <0.001 | 103 (55.7, 149) | <0.001 |
| Within INR | 7.50 (-61.4, 76.4) | 0.825 | 7.50 (-61.4, 76.4) | 0.825 |
| **V9 vs V8** |  |  |  |  |
| Within HIV– | 2.50 (-8.50, 13.5) | 0.647 | -2.50 (-22.2, 17.2) | 0.798 |
| Within HIV+ | 12.5 (-10.7, 35.9) | 0.281 | 12.5 (-13.8, 38.8) | 0.341 |
| Within IR | 12.5 (-42.5, 67.5) | 0.646 | 12.5 (-45.9, 70.9) | 0.665 |
| Within INR | 12.5 (-105, 130) | 0.830 | 12.5 (-111, 136) | 0.837 |
| **Dual spike** | **All data** | | **Out-of-window data excluded** | |
|  | **Median change (95% CI)** | ***p*** | **Median change (95% CI)** | ***p*** |
| **V8 vs V1** |  |  |  |  |
| Within HIV– | 5.00 (1.57, 8.43) | 0.006 | 5.00 (1.21, 8.79) | 0.011 |
| Within HIV+ | 2.50 (-1.56, 6.56) | 0.219 | 2.50 (-1.43, 6.43) | 0.204 |
| Within IR | 5.00 (0.81, 9.19) | 0.021 | 5.00 (0.81, 9.19) | 0.021 |
| Within INR | 0.00 (-4.41, 4.41) | 1.000 | 0.00 (-4.41, 4.41) | 1.000 |
| **V9 vs V8** |  |  |  |  |
| Within HIV– | 0.00 (-3.07, 3.07) | 1.000 | -2.50 (-5.91, 0.91) | 0.146 |
| Within HIV+ | 2.50 (-5.10, 10.1) | 0.509 | 2.50 (-5.10, 10.1) | 0.508 |
| Within IR | 12.5 (4.07, 20.9) | 0.005 | 12.5 (3.54, 21.5) | 0.008 |
| Within INR | 0.00 (-5.82, 5.82) | 1.000 | 0.00 (-5.99, 5.99) | 1.000 |

**Supplemental table S16**. Changes in the intact HIV reservoir relative to the baseline (V1) following COVID-19 vaccination, by IPDA and quantile regression analysis.

| **All HIV+** | | **V1** | | | **V8** | **V9** |
| --- | --- | --- | --- | --- | --- | --- |
| **Intact proviruses per 10^6^ CD4+ T cells** | |  | | |  |  |
| n | | 45 | | | 44 | 41 |
| Median (IQR) | | 90.9 (32.5, 203) | | | 122 (22.4, 222) | 95.0 (33.0, 217) |
| Range | | (0.0, 2954) | | | (0.0, 549) | (0.0, 2460) |
| **Change** | |  | | |  |  |
| n | | - | | | 44 | 41 |
| Median (IQR) | | - | | | 1.2 (-18.5, 34.1) | -1.0 (-40.0, 21.7) |
| Range | |  | | | (-245, 169) | (-494, 308) |
| *p* | | - | | | 0.920 | 0.952 |
| **IRs** | | **V1** | | | **V8** | **V9** |
| **Intact proviruses per 10^6^ CD4+ T cells** | |  | | |  |  |
| n | | 29 | | | 29 | 27 |
| Median (IQR) | | 84.6 (16.1, 158) | | | 83.0 (18.2, 161) | 76.7 (16.5, 141) |
| Range | | (0.0, 495) | | | (0.0, 300) | (0.0, 419) |
| **Change** | |  | | |  |  |
| n | | - | | | 29 | 27 |
| Median (IQR) | | - | | | 1.3 (-14.6, 25.0) | 0.0 (-40.0, 21.3) |
| Range | |  | | | (-245, 92.0) | (-156, 118) |
| *p* | | - | | | 0.887 | 1.000 |
| **INRs** | | **V1** | | | **V8** | **V9** |
| **Intact proviruses per 10^6^ CD4+ T cells** | |  | | |  |  |
| n | | 12 | | | 11 | 10 |
| Median (IQR) | | 141 (60.4, 378) | | | 139 (34.0, 391) | 166 (39.7, 322) |
| Range | | (7.1, 2954) | | | (6.9, 549) | (20.2, 2460) |
| **Change** | |  | | |  |  |
| n | | - | | | 11 | 10 |
| Median (IQR) | | - | | | 0.0 (-35.8, 89.0) | -33.6 (-54.0, 13.1) |
| Range | |  | | | (-127, 169) | (-494, 308) |
| *p* | | - | | | 1.000 | 0.136 |

**Supplemental table S17**. Changes in the number of samples with detectable HIV viral load (above 40 copies/mL) – for all PWH, IRs, and INRs.

|  | **Detectable VL, n (%)** | | |
| --- | --- | --- | --- |
|  | **All HIV+** | **IRs** | **INRs** |
| V1 | 3/68 (4.4) | 0/42 (0.0) | 1/20 (5.0) |
| V2 | 1/44 (2.3) | 1/29 (3.4) | 0/11 (0.0) |
| V3 | 2/44 (4.5) | 0/29 (0.0) | 2/11 (18.2) |
| V4 | 2/45 (4.4) | 2/29 (6.9) | 0/12 (0.0) |
| V4a | 3/35 (8.6) | 1/22 (4.5) | 2/10 (20.0) |
| V5 | 1/39 (2.6) | 0/23 (0.0) | 1/12 (8.3) |
| V6 | 3/40 (7.5) | 2/24 (8.3) | 1/12 (8.3) |
| V7 | 5/43 (11.6) | 4/27 (14.8) | 0/12 (0.0) |
| V8 | 4/66 (6.1) | 1/41 (2.4) | 1/19 (5.3) |
| V8a | 3/42 (7.1) | 1/28 (3.6) | 1/9 (11.1) |
| V8b | 1/56 (1.8) | 0/31 (0.0) | 0/19 (0.0) |
| V8c | 2/33 (6.1) | 0/16 (0.0) | 2/15 (13.3) |
| V9 | 5/62 (8.1) | 1/39 (2.6) | 1/17 (5.9) |

**Supplemental table S18**. Changes in T cell gag and nef responses relative to the baseline following COVID-19 vaccination in PWH, by ELISpot and quantile regression analysis.

|  | **All HIV+** | | | **IRs** | | | **INRs** | | |
| --- | --- | --- | --- | --- | --- | --- | --- | --- | --- |
| **Variable** | **V1** | **V8** | **V9** | **V1** | **V8** | **V9** | **V1** | **V8** | **V9** |
| **IFN-γ Gag** |  |  |  |  |  |  |  |  |  |
| n | 28 | 29 | 29 | 18 | 19 | 19 | 6 | 6 | 6 |
| Median (IQR) | 205 (77.5, 448) | 185 (45.0, 413) | 190 (45.0, 673) | 268 (82.5, 435) | 203 (80.0, 413) | 255 (90.0, 595) | 72.5 (10.0, 165) | 65.3 (5.0, 135) | 48.8 (17.5, 153) |
| Range | (0.0, 2070) | (0.0, 1700) | (0.0, 2345) | (10.0, 538) | (7.5, 1145) | (0.0, 1538) | (0.0, 218) | (0.0, 223) | (0.0, 673) |
| **Change in IFN-γ gag** |  |  |  |  |  |  |  |  |  |
| n | - | 28 | 28 | - | 18 | 18 | - | 6 | 6 |
| Median (IQR) | - | -2.5 (-45.0, 25.0) | 40.0 (-28.8, 261) | - | 6.3 (-37.5, 32.5) | 50.0 (-2.5, 260) | - | -20.0 (-47.5, 13.0) | -13.8 (-65.0, 27.5) |
| Range | - | (-370, 633) | (-193, 1025) | - | (-175, 633) | (-193, 1025) | - | (-82.5, 113) | (-95.0, 563) |
| *p* | - | 0.776 | 0.456 | - | 0.784 | 0.221 | - | 0.718 | 0.863 |
| **IFN-γ nef** |  |  |  |  |  |  |  |  |  |
| n | 28 | 29 | 29 | 18 | 19 | 19 | 6 | 6 | 6 |
| Median (IQR) | 33.8 (10.0, 118) | 35.0 (5.0, 143) | 32.5 (3.8, 128) | 38.8 (25.0, 115) | 50.0 (20.0, 168) | 42.5 (17.5, 165) | 5.0 (0.0, 20.0) | 2.5 (0.0, 27.5) | 2.5 (0.0, 67.5) |
| Range | (0.0, 1015) | (0.0, 1565) | (0.0, 1523) | (0.0, 1015) | (0.0, 1565) | (0.0, 1523) | (0.0, 318) | (0.0, 111) | (0.0, 108) |
| **Change in IFN-γ nef** |  |  |  |  |  |  |  |  |  |
| n | - | 28 | 28 | - | 18 | 18 | - | 6 | 6 |
| Median (IQR) | - | 0.0 (-13.8, 18.8) | 1.3 (-13.8, 48.8) | - | 12.5 (-12.5, 52.5) | 3.8 (-15.0, 50.0) | - | -1.3 (-2.5, 0.0) | -1.3 (-5.0, 2.5) |
| Range | - | (-207, 550) | (-418, 508) | - | (-205, 550) | (-418, 508) | - | (-207, 7.5) | (-210, 47.5) |
| *p* | - | 1.000 | 1.000 | - | 0.381 | 0.859 | - | 1.000 | 1.000 |
| **IL-2 gag** |  |  |  |  |  |  |  |  |  |
| n | 28 | 29 | 29 | 18 | 19 | 19 | 6 | 6 | 6 |
| Median (IQR) | 115 (36.3, 221) | 105 (41.6, 178) | 133 (40.0, 265) | 119 (65.0, 198) | 110 (52.5, 178) | 133 (62.5, 370) | 7.5 (0.0, 57.5) | 44.6 (0.0, 105) | 25.0 (2.5, 203) |
| Range | (0.0, 583) | (0.0, 740) | (0.0, 863) | (10.0, 363) | (0.0, 740) | (22.5, 680) | (0.0, 255) | (0.0, 234) | (0.0, 230) |
| **Change in IL-2 gag** |  |  |  |  |  |  |  |  |  |
| n | - | 28 | 28 | - | 18 | 18 | - | 6 | 6 |
| Median (IQR) | - | 1.3 (-24.3, 40.8) | 36.3 (-3.8, 129) | - | -3.8 (-27.5, 40.0) | 41.3 (-2.5, 113) | - | 20.8 (-15.0, 47.5) | 5.0 (-12.5, 40.0) |
| Range | - | (-87.5, 460) | (-92.5, 430) | - | (-57.5, 460) | (-92.5, 430) | - | (-21.0, 47.5) | (-25.0, 145) |
| *p* | - | 1.000 | 0.050 | - | 0.852 | 0.097 | - | 1.000 | 1.000 |
| **IL-2 nef** |  |  |  |  |  |  |  |  |  |
| n | 28 | 29 | 29 | 18 | 19 | 19 | 6 | 6 | 6 |
| Median (IQR) | 16.3 (0.0, 55.0) | 12.5 (1.3, 60.0) | 25.0 (0.0, 72.5) | 25.0 (5.0, 62.5) | 22.5 (5.0, 65.0) | 30.0 (10.0, 75.0) | 1.3 (0.0, 2.5) | 2.5 (0.0, 10.0) | 0.0 (0.0, 2.5) |
| Range | (0.0, 443) | (0.0, 515) | (0.0, 450) | (0.0, 215) | (0.0, 515) | (0.0, 450) | (0.0, 443) | (0.0, 327) | (0.0, 288) |
| **Change in IL-2 nef** |  |  |  |  |  |  |  |  |  |
| n | - | 28 | 28 | - | 18 | 18 | - | 6 | 6 |
| Median (IQR) | - | 1.3 (-6.3, 7.5) | 0.0 (-1.9, 32.5) | - | 5.0 (-1.3, 10.0) | 2.5 (-2.5, 42.5) | - | 0.0 (-2.5, 2.5) | 0.0 (-2.5, 0.0) |
| Range | - | (-116, 300) | (-155, 235) | - | (-27.5, 300) | (-13.8, 235) | - | (-116, 10.0) | (-155, 0.0) |
| *p* | - | 0.486 | 1.000 | - | 0.116 | 0.786 | - | 1.000 | 1.000 |
| **Dual gag** |  |  |  |  |  |  |  |  |  |
| n | 28 | 29 | 29 | 18 | 19 | 19 | 6 | 6 | 6 |
| Median (IQR) | 37.5 (10.0, 67.5) | 40.0 (5.0, 62.5) | 47.5 (5.0, 105) | 42.5 (12.5, 75.0) | 40.0 (12.5, 62.5) | 47.5 (7.5, 105) | 8.1 (0.0, 25.0) | 6.3 (2.5, 50.0) | 3.8 (0.0, 52.5) |
| Range | (0.0, 393) | (0.0, 298) | (0.0, 443) | (0.0, 135) | (0.0, 278) | (0.0, 368) | (0.0, 50.0) | (0.0, 54.0) | (0.0, 97.5) |
| **Change in Dual gag** |  |  |  |  |  |  |  |  |  |
| n | - | 28 | 28 | - | 18 | 18 | - | 6 | 6 |
| Median (IQR) | - | 2.5 (-8.8, 8.8) | 10.0 (-3.8, 65.0) | - | 0.0 (-10.0, 12.5) | 16.3 (-5.0, 65.0) | - | 3.2 (0.0, 4.0) | 1.3 (-6.3, 7.5) |
| Range | - | (-95.0, 143) | (-30.0, 233) | - | (-27.5, 143) | (-30.0, 233) | - | (-7.5, 25.0) | (-10.0, 72.5) |
| *p* | - | 0.407 | 0.556 | - | 1.000 | 0.387 | - | 0.508 | 1.000 |
| **Dual nef** |  |  |  |  |  |  |  |  |  |
| n | 28 | 29 | 29 | 18 | 19 | 19 | 6 | 6 | 6 |
| Median (IQR) | 7.5 (0.0, 26.3) | 7.5 (0.0, 17.5) | 2.5 (0.0, 25.0) | 7.5 (2.5, 25.0) | 10.0 (2.5, 25.0) | 5.0 (2.5, 25.0) | 0.0 (0.0, 7.5) | 0.0 (0.0, 0.0) | 0.0 (0.0, 5.0) |
| Range | (0.0, 153) | (0.0, 288) | (0.0, 283) | (0.0, 153) | (0.0, 288) | (0.0, 283) | (0.0, 138) | (0.0, 57.0) | (0.0, 50.0) |
| **Change in Dual nef** |  |  |  |  |  |  |  |  |  |
| n | - | 28 | 28 | - | 18 | 18 | - | 6 | 6 |
| Median (IQR) | - | 0.0 (-5.0, 3.8) | 0.0 (-3.8, 11.3) | - | 2.5 (0.0, 5.0) | 1.3 (-5.0, 12.5) | - | 0.0 (-7.5, 0.0) | 0.0 (-2.5, 0.0) |
| Range | - | (-80.5, 135) | (-87.5, 130) | - | (-20.0, 135) | (-25.0, 130) | - | (-80.5, 0.0) | (-87.5, 0.0) |
| *p* | - | 1.000 | 1.000 | - | 0.215 | 1.000 | - | 1.000 | 1.000 |

**Supplemental table S19**. Changes in CD4-related clinical parameters in PWH relative to the baseline (mixed effects linear regression with participant-specific random intercept).

| **Variable** | | **V1^a^** | **V2** | | | **V3** | **V4** | | | **V4a** | **V5** | **V6** |
| --- | --- | --- | --- | --- | --- | --- | --- | --- | --- | --- | --- | --- |
| **CD4+ T cell count, cells/μL** | |  |  | | |  |  | | |  |  |  |
| n | | 68 | 44 | | | 44 | 44 | | | 35 | 39 | 40 |
| Median (IQR) | | 527 (364, 665) | 597 (415, 810) | | | 603 (407, 776) | 569 (417, 760) | | | 529 (408, 686) | 507 (390, 664) | 543 (377, 748) |
| Range | | (74.0, 1784) | (106, 1817) | | | (115, 1590) | (93.0, 1846) | | | (266, 1432) | (133, 1518) | (113, 1627) |
| **Change in CD4+ T cell count** | |  |  | | |  |  | | |  |  |  |
| n | | - | 44 | | | 44 | 44 | | | 35 | 39 | 40 |
| Median (IQR) | | - | 34.5 (-19.5, 114) | | | 41.5 (-34.0, 120.5) | 15.0 (-22.0, 74.0) | | | 6.0 (-58.0, 88.0) | 5.0 (-87.0, 50.0) | -5.5 (-54.0, 48.5) |
| Range | | - | (-280, 386) | | | (-323, 323) | (-340, 287) | | | (-319, 404) | (-266, 455) | (-217, 305) |
| **CD4+ T cell %** | |  |  | | |  |  | | |  |  |  |
| n | | 68 | 44 | | | 44 | 44 | | | 35 | 39 | 40 |
| Median (IQR) | | 30.0 (22.0, 38.5) | 32.0 (22.5, 38.4) | | | 30.3 (22.2, 39.0) | 30.7 (21.5, 39.4) | | | 28.2 (22.5, 36.0) | 27.0 (21.8, 36.3) | 31.2 (21.0, 38.7) |
| Range | | (12.9, 62.3) | (14.0, 61.4) | | | (13.5, 60.6) | (11.7, 60.1) | | | (12.7, 59.1) | (12.8, 59.2) | (13.6, 58.6) |
| **Change in CD4+ T cell %** | |  |  | | |  |  | | |  |  |  |
| n | | - | 44 | | | 44 | 44 | | | 35 | 39 | 40 |
| Median (IQR) | | - | 0.1 (-1.6, 1.3) | | | -0.1 (-2.0, 1.5) | 0.1 (-2.1, 1.7) | | | 0.1 (-1.6, 1.3) | 0.0 (-2.1, 1.8) | -0.4 (-1.6, 2.3) |
| Range | | - | (-8.2, 6.7) | | | (-6.1, 5.4) | (-6.0, 5.8) | | | (-9.0, 5.8) | (-7.0, 4.6) | (-9.4, 5.6) |
| **CD4+/CD8+ T cell ratio** | |  |  | | |  |  | | |  |  |  |
| n | | 68 | 44 | | | 44 | 44 | | | 35 | 39 | 40 |
| Median (IQR) | | 0.9 (0.5, 1.2) | 0.9 (0.6, 1.2) | | | 0.9 (0.5, 1.3) | 0.9 (0.5, 1.3) | | | 0.7 (0.5, 1.2) | 0.7 (0.4, 1.2) | 0.8 (0.5, 1.2) |
| Range | | (0.2, 6.1) | (0.3, 5.5) | | | (0.2, 5.3) | (0.2, 5.4) | | | (0.3, 2.3) | (0.2, 5.3) | (0.2, 5.3) |
| **Change in CD4+/CD8+ T cell ratio** | |  |  | | |  |  | | |  |  |  |
| n | | - | 44 | | | 44 | 44 | | | 35 | 39 | 40 |
| Median (IQR) | | - | 0.0 (-0.1, 0.1) | | | 0.0 (0.0, 0.1) | 0.0 (-0.1, 0.1) | | | 0.0 (0.0, 0.1) | 0.0 (-0.1, 0.1) | 0.0 (-0.1, 0.1) |
| Range | | - | (-0.6, 0.3) | | | (-0.8, 0.2) | (-0.7, 0.8) | | | (-0.3, 0.3) | (-0.8, 0.2) | (-0.8, 0.3) |
| **Variable** | | **V1^a^** | **V7** | | | **V8** | **V8a** | | | **V8b** | **V8c** | **V9** |
| **CD4+ T cell count, cells/μL** | |  |  | | |  |  | | |  |  |  |
| n | | 68 | 43 | | | 66 | 42 | | | 56 | 33 | 63 |
| Median (IQR) | | 527 (364, 665) | 563 (394, 727) | | | 537 (356, 677) | 615 (411, 776) | | | 498 (318, 715) | 492 (356, 658) | 566 (401, 680) |
| Range | | (74.0, 1784) | (95.0, 1806) | | | (119, 1863) | (204, 1837) | | | (120, 1480) | (142, 1037) | (128, 1980) |
| **Change in CD4+ T cell count** | |  |  | | |  |  | | |  |  |  |
| n | | - | 43 | | | 66 | 42 | | | 56 | 33 | 63 |
| Median (IQR) | | - | 12.0 (-35.0, 55.0) | | | -12.5 (-78.0, 71.0) | 22.0 (-45.0, 71.0) | | | -19.0 (-74.5, 55.5) | -12.0 (-51.0, 39.0) | 12.0 (-52.0, 67.0) |
| Range | | - | (-305, 284) | | | (-252, 262) | (-232, 402) | | | (-304, 509) | (-482, 287) | (-527, 315) |
| **CD4+ T cell %** | |  |  | | |  |  | | |  |  |  |
| n | | 68 | 43 | | | 66 | 42 | | | 56 | 33 | 63 |
| Median (IQR) | | 30.0 (22.0, 38.5) | 30.1 (21.9, 38.2) | | | 30.3 (23.4, 39.9) | 32.0 (23.2, 38.5) | | | 31.1 (23.1, 39.3) | 28.3 (20.9, 37.2) | 31.1 (22.9, 40.4) |
| Range | | (12.9, 62.3) | (12.9, 61.5) | | | (13.4, 57.7) | (14.4, 61.2) | | | (13.0, 59.8) | (12.6, 54.6) | (10.7, 57.7) |
| **Change in CD4+ T cell %** | |  |  | | |  |  | | |  |  |  |
| n | | - | 43 | | | 66 | 42 | | | 56 | 33 | 63 |
| Median (IQR) | | - | 0.3 (-2.2, 1.4) | | | 0.0 (-1.8, 2.3) | -0.4 (-2.0, 1.7) | | | 0.4 (-1.8, 2.5) | 0.0 (-1.4, 1.7) | -0.1 (-1.1, 1.6) |
| Range | | - | (-6.7, 6.2) | | | (-6.4, 10.4) | (-10.4, 4.8) | | | (-7.2, 16.4) | (-7.5, 7.7) | (-7.5, 14.5) |
| **CD4+/CD8+ T cell ratio** | |  |  | | |  |  | | |  |  |  |
| n | | 68 | 43 | | | 66 | 42 | | | 56 | 33 | 63 |
| Median (IQR) | | 0.9 (0.5, 1.2) | 0.8 (0.5, 1.2) | | | 0.8 (0.5, 1.2) | 0.9 (0.6, 1.2) | | | 0.8 (0.5, 1.2) | 0.8 (0.5, 1.1) | 0.9 (0.6, 1.2) |
| Range | | (0.2, 6.1) | (0.2, 5.8) | | | (0.2, 5.7) | (0.2, 5.8) | | | (0.2, 5.2) | (0.2, 2.2) | (0.2, 5.3) |
| **Change in CD4+/CD8+ T cell ratio** | |  |  | | |  |  | | |  |  |  |
| n | | - | 43 | | | 66 | 42 | | | 56 | 33 | 63 |
| Median (IQR) | | - | 0.0 (-0.1, 0.1) | | | 0.0 (-0.1, 0.1) | 0.0 (-0.1, 0.1) | | | 0.0 (-0.0, 0.1) | 0.0 (0.0, 0.1) | 0.0 (-0.1, 0.1) |
| Range | | - | (-0.4, 0.2) | | | (-0.4, 0.3) | (-0.5, 0.3) | | | (-0.9, 0.9) | (-0.3, 0.5) | (-0.8, 0.6) |

^a^If baseline data were not available, data at screening were used.

**Supplemental table S20**. Estimated within-group/subgroup mean change from baseline (V1) in three clinical parameters, by regression analysis.

| **All HIV+** | | **CD4+ T cell count, cells/μL** | | | **CD4+ T cell %** | | | | **CD4+/CD8+ T cell ratio** | | | |
| --- | --- | --- | --- | --- | --- | --- | --- | --- | --- | --- | --- | --- |
|  |  | **Estimated mean change (95% CI)** | | ***p*** | **Estimated mean change (95% CI)** | | | ***p*** | **Estimated mean change (95% CI)** | | | ***p*** |
| **V2** | | 47.9 (12.7, 83.1) | | 0.008 | -0.1 (-0.9, 0.7) | | | 0.804 | -0.01 (-0.05, 0.04) | | | 0.792 |
| **V3** | | 24.5 (-10.7, 59.7) | | 0.172 | -0.1 (-1.0, 0.7) | | | 0.779 | 0.00 (-0.04, 0.05) | | | 0.952 |
| **V4** | | 26.1 (-9.1, 61.3) | | 0.145 | -0.0 (-0.8, 0.8) | | | 0.981 | 0.01 (-0.03, 0.06) | | | 0.586 |
| **V4a** | | 21.4 (-16.6, 59.4) | | 0.269 | -0.2 (-1.1, 0.8) | | | 0.739 | 0.00 (-0.05, 0.05) | | | 0.911 |
| **V5** | | 4.1 (-32.5, 40.7) | | 0.826 | -0.2 (-1.0, 0.7) | | | 0.722 | -0.01 (-0.05, 0.04) | | | 0.763 |
| **V6** | | 7.8 (-28.5, 44.1) | | 0.674 | -0.2 (-1.0, 0.7) | | | 0.684 | -0.02 (-0.06, 0.03) | | | 0.413 |
| **V7** | | 2.0 (-33.5, 37.4) | | 0.914 | -0.3 (-1.1, 0.6) | | | 0.539 | -0.02 (-0.07, 0.02) | | | 0.322 |
| **V8** | | -6.7 (-37.4, 24.1) | | 0.670 | 0.2 (-0.6, 0.9) | | | 0.675 | -0.01 (-0.05, 0.03) | | | 0.540 |
| **V8a** | | 22.5 (-13.1, 58.1) | | 0.216 | -0.0 (-0.9, 0.8) | | | 0.914 | -0.02 (-0.06, 0.03) | | | 0.479 |
| **V8b** | | -10.5 (-43.0, 21.9) | | 0.524 | 0.2 (-0.6, 0.9) | | | 0.665 | 0.01 (-0.03, 0.05) | | | 0.702 |
| **V8c** | | -11.6 (-50.3, 27.1) | | 0.556 | -0.2 (-1.1, 0.7) | | | 0.697 | 0.00 (-0.05, 0.05) | | | 0.953 |
| **V9** | | 6.0 (-25.2, 37.2) | | 0.705 | -0.0 (-0.7, 0.7) | | | 0.988 | -0.00 (-0.04, 0.03) | | | 0.806 |
| **IRs** | **CD4+ T cell count, cells/μL** | | | | **CD4+ T cell %** | | | | **CD4+/CD8+ T cell ratio** | | | |
|  | **Estimated mean change (95% CI)** | | | ***p*** | **Estimated mean change (95% CI)** | | | ***p*** | **Estimated mean change (95% CI)** | | | ***p*** |
| **V2** | 73.0 (28.5, 117.4) | | | 0.001 | -0.4 (-1.4, 0.7) | | | 0.505 | -0.03 (-0.08, 0.03) | | | 0.378 |
| **V3** | 28.4 (-16.0, 72.8) | | | 0.210 | -0.4 (-1.4, 0.7) | | | 0.509 | -0.01 (-0.07, 0.05) | | | 0.683 |
| **V4** | 32.8 (-12.1, 77.8) | | | 0.152 | -0.1 (-1.1, 1.0) | | | 0.863 | 0.00 (-0.06, 0.06) | | | 0.922 |
| **V4a** | 12.9 (-35.8, 61.5) | | | 0.603 | -0.6 (-1.7, 0.5) | | | 0.309 | -0.02 (-0.08, 0.04) | | | 0.567 |
| **V5** | -5.8 (-53.7, 42.1) | | | 0.812 | -0.6 (-1.7, 0.6) | | | 0.317 | -0.03 (-0.09, 0.03) | | | 0.370 |
| **V6** | -1.8 (-49.1, 45.4) | | | 0.939 | -0.5 (-1.6, 0.6) | | | 0.338 | -0.04 (-0.10, 0.02) | | | 0.158 |
| **V7** | -7.2 (-52.7, 38.3) | | | 0.755 | -0.9 (-1.9, 0.2) | | | 0.108 | -0.05 (-0.11, 0.01) | | | 0.079 |
| **V8** | 0.3 (-39.3, 39.8) | | | 0.990 | 0.2 (-0.8, 1.1) | | | 0.741 | -0.03 (-0.08, 0.02) | | | 0.310 |
| **V8a** | 38.6 (-6.2, 83.5) | | | 0.091 | -0.4 (-1.4, 0.7) | | | 0.467 | -0.04 (-0.10, 0.02) | | | 0.149 |
| **V8b** | -10.2 (-53.6, 33.2) | | | 0.645 | -0.5 (-1.5, 0.6) | | | 0.368 | -0.02 (-0.08, 0.04) | | | 0.477 |
| **V8c** | -36.1 (-90.7, 18.4) | | | 0.194 | -0.6 (-1.8, 0.7) | | | 0.397 | -0.03 (-0.10, 0.04) | | | 0.442 |
| **V9** | 6.4 (-33.7, 46.6) | | | 0.753 | -0.1 (-1.0, 0.8) | | | 0.830 | -0.02 (-0.07, 0.03) | | | 0.394 |
| **INRs** | | **CD4+ T cell count, cells/μL** | | | **CD4+ T cell %** | | | | **CD4+/CD8+ T cell ratio** | | | |
|  |  | **Estimated mean change (95% CI)** | | ***p*** | **Estimated mean change (95% CI)** | | | ***p*** | **Estimated mean change (95% CI)** | | | ***p*** |
| **V2** | | 3.7 (-65.5, 72.9) | | 0.917 | 0.2 (-1.4, 1.9) | | | 0.762 | 0.02 (-0.07, 0.10) | | | 0.730 |
| **V3** | | 17.2 (-52.0, 86.4) | | 0.625 | 0.7 (-0.9, 2.3) | | | 0.415 | 0.02 (-0.07, 0.11) | | | 0.704 |
| **V4** | | 34.2 (-33.0, 101.4) | | 0.318 | 0.5 (-1.0, 2.1) | | | 0.497 | 0.02 (-0.06, 0.11) | | | 0.580 |
| **V4a** | | 37.8 (-33.6, 109.3) | | 0.299 | 0.8 (-0.8, 2.5) | | | 0.333 | 0.03 (-0.07, 0.12) | | | 0.589 |
| **V5** | | 32.8 (-34.4, 100.1) | | 0.337 | 0.9 (-0.7, 2.4) | | | 0.288 | 0.02 (-0.07, 0.10) | | | 0.715 |
| **V6** | | 20.8 (-46.5, 88.0) | | 0.544 | 0.6 (-1.0, 2.1) | | | 0.491 | 0.01 (-0.08, 0.09) | | | 0.860 |
| **V7** | | 12.3 (-55.0, 79.5) | | 0.720 | 0.9 (-0.6, 2.5) | | | 0.244 | 0.01 (-0.08, 0.09) | | | 0.860 |
| **V8** | | -14.3 (-72.0, 43.4) | | 0.627 | 0.7 (-0.6, 2.1) | | | 0.302 | 0.01 (-0.06, 0.09) | | | 0.699 |
| **V8a** | | 8.5 (-65.5, 82.6) | | 0.821 | 1.2 (-0.5, 2.9) | | | 0.175 | 0.05 (-0.05, 0.14) | | | 0.339 |
| **V8b** | | 8.3 (-49.3, 66.0) | | 0.776 | 1.7 (0.4, 3.1) | | | 0.013 | 0.05 (-0.02, 0.13) | | | 0.164 |
| **V8c** | | 12.5 (-49.5, 74.5) | | 0.692 | 0.6 (-0.8, 2.1) | | | 0.412 | 0.04 (-0.04, 0.12) | | | 0.358 |
| **V9** | | 20.3 (-38.3, 79.0) | | 0.496 | 0.5 (-0.9, 1.8) | | | 0.512 | 0.03 (-0.04, 0.11) | | | 0.425 |

**Supplemental table S21**. Within-group/subgroup changes in CD4^+^ T cell parameters between neighbouring time points in PWH based on mixed effects linear regression.

| **All HIV+** | **CD4+ T cell count, cells/μL** | | **CD4+ T cell %** | | **CD4+/CD8+ T cell ratio** | |
| --- | --- | --- | --- | --- | --- | --- |
|  | **Estimated mean change (95% CI)** | ***p*** | **Estimated mean change (95% CI)** | ***p*** | **Estimated mean change (95% CI)** | ***p*** |
| V2 vs V1 | 47.9 (12.7, 83.1) | 0.008 | -0.1 (-0.9, 0.7) | 0.805 | -0.01 (-0.05, 0.04) | 0.793 |
| V3 vs V2 | -23.6 (-61.5, 14.4) | 0.223 | -0.0 (-0.9, 0.9) | 0.973 | 0.01 (-0.04, 0.06) | 0.765 |
| V4 vs V3 | 1.6 (-36.3, 39.6) | 0.933 | 0.1 (-0.8, 1.0) | 0.812 | 0.01 (-0.04, 0.06) | 0.653 |
| V4a vs V4 | -4.8 (-45.3, 35.8) | 0.817 | -0.1 (-1.1, 0.8) | 0.771 | -0.01 (-0.06, 0.04) | 0.713 |
| V5 vs V4a | -17.3 (-58.9, 24.4) | 0.416 | -0.0 (-1.0, 1.0) | 0.992 | -0.01 (-0.06, 0.04) | 0.714 |
| V6 vs V5 | 3.7 (-36.5, 43.9) | 0.856 | -0.0 (-1.0, 0.9) | 0.966 | -0.01 (-0.06, 0.04) | 0.641 |
| V7 vs V6 | -5.8 (-44.9, 33.3) | 0.771 | -0.1 (-1.0, 0.8) | 0.857 | -0.00 (-0.05, 0.05) | 0.891 |
| V8 vs V7 | -8.5 (-44.2, 27.2) | 0.640 | 0.4 (-0.4, 1.3) | 0.330 | 0.01 (-0.03, 0.06) | 0.648 |
| V8a vs V8 | 29.2 (-6.6, 65.0) | 0.110 | -0.2 (-1.1, 0.7) | 0.641 | -0.00 (-0.05, 0.04) | 0.859 |
| V8b vs V8a | -33.1 (-70.0, 3.7) | 0.078 | 0.2 (-0.7, 1.1) | 0.629 | 0.02 (-0.02, 0.07) | 0.308 |
| V8c vs V8b | -1.1 (-40.8, 38.6) | 0.956 | -0.4 (-1.3, 0.6) | 0.463 | -0.01 (-0.06, 0.04) | 0.798 |
| V9 vs V8c | 19.0 (-20.3, 58.3) | 0.342 | 0.2 (-0.7, 1.1) | 0.692 | -0.01 (-0.06, 0.04) | 0.801 |
| **IRs** | **CD4+ T cell count, cells/μL** | | **CD4+ T cell %** | | **CD4+/CD8+ T cell ratio** | |
|  | **Estimated mean change (95% CI)** | ***p*** | **Estimated mean change (95% CI)** | ***p*** | **Estimated mean change (95% CI)** | ***p*** |
| V2 vs V1 | 73.0 (28.5, 117.4) | 0.001 | -0.4 (-1.4, 0.7) | 0.505 | -0.03 (-0.08, 0.03) | 0.379 |
| V3 vs V2 | -44.6 (-91.8, 2.7) | 0.064 | 0.0 (-1.1, 1.1) | 0.995 | 0.01 (-0.05, 0.07) | 0.657 |
| V4 vs V3 | 4.4 (-43.3, 52.1) | 0.856 | 0.3 (-0.9, 1.4) | 0.651 | 0.01 (-0.05, 0.08) | 0.636 |
| V4a vs V4 | -20.0 (-71.6, 31.7) | 0.448 | -0.5 (-1.7, 0.7) | 0.420 | -0.02 (-0.09, 0.05) | 0.533 |
| V5 vs V4a | -18.7 (-72.7, 35.3) | 0.497 | 0.0 (-1.2, 1.3) | 0.978 | -0.01 (-0.08, 0.06) | 0.779 |
| V6 vs V5 | 4.0 (-48.9, 56.8) | 0.882 | 0.0 (-1.2, 1.3) | 0.959 | -0.02 (-0.08, 0.05) | 0.651 |
| V7 vs V6 | -5.4 (-56.0, 45.2) | 0.835 | -0.3 (-1.5, 0.9) | 0.582 | -0.01 (-0.07, 0.06) | 0.796 |
| V8 vs V7 | 7.5 (-38.2, 53.1) | 0.748 | 1.0 (-0.0, 2.1) | 0.060 | 0.03 (-0.03, 0.09) | 0.385 |
| V8a vs V8 | 38.4 (-6.7, 83.4) | 0.095 | -0.5 (-1.6, 0.5) | 0.310 | -0.02 (-0.07, 0.04) | 0.584 |
| V8b vs V8a | -48.8 (-96.2, -1.4) | 0.044 | -0.1 (-1.2, 1.0) | 0.892 | 0.02 (-0.04, 0.08) | 0.474 |
| V8c vs V8b | -25.9 (-82.6, 30.7) | 0.368 | -0.1 (-1.4, 1.2) | 0.899 | -0.01 (-0.08, 0.07) | 0.845 |
| V9 vs V8c | 42.5 (-12.5, 97.6) | 0.129 | 0.4 (-0.8, 1.7) | 0.494 | 0.01 (-0.07, 0.08) | 0.889 |
| **INRs** | **CD4+ T cell count, cells/μL** | | **CD4+ T cell %** | | **CD4+/CD8+ T cell ratio** | |
|  | **Estimated mean change (95% CI)** | ***p*** | **Estimated mean change (95% CI)** | ***p*** | **Estimated mean change (95% CI)** | ***p*** |
| V2 vs V1 | 3.8 (-65.4, 73.0) | 0.915 | 0.3 (-1.4, 1.9) | 0.762 | 0.02 (-0.07, 0.11) | 0.730 |
| V3 vs V2 | 12.9 (-64.2, 89.9) | 0.743 | 0.4 (-1.4, 2.2) | 0.647 | 0.00 (-0.10, 0.10) | 0.977 |
| V4 vs V3 | 17.0 (-58.2, 92.2) | 0.657 | -0.1 (-1.9, 1.6) | 0.886 | 0.01 (-0.09, 0.10) | 0.885 |
| V4a vs V4 | 3.5 (-73.8, 80.8) | 0.929 | 0.3 (-1.5, 2.1) | 0.761 | 0.00 (-0.10, 0.10) | 0.986 |
| V5 vs V4a | -4.8 (-82.2, 72.5) | 0.902 | 0.0 (-1.8, 1.8) | 0.977 | -0.01 (-0.11, 0.09) | 0.856 |
| V6 vs V5 | -12.1 (-85.5, 61.3) | 0.747 | -0.3 (-2.0, 1.4) | 0.732 | -0.01 (-0.10, 0.09) | 0.863 |
| V7 vs V6 | -8.5 (-81.9, 64.9) | 0.820 | 0.4 (-1.3, 2.1) | 0.662 | 0.00 (-0.09, 0.09) | 1.000 |
| V8 vs V7 | -26.0 (-94.2, 42.1) | 0.453 | -0.2 (-1.8, 1.4) | 0.783 | 0.01 (-0.08, 0.09) | 0.878 |
| V8a vs V8 | 23.1 (-51.8, 97.9) | 0.545 | 0.5 (-1.3, 2.2) | 0.583 | 0.03 (-0.06, 0.13) | 0.517 |
| V8b vs V8a | -0.4 (-75.3, 74.4) | 0.991 | 0.5 (-1.2, 2.3) | 0.561 | 0.01 (-0.09, 0.10) | 0.899 |
| V8c vs V8b | 4.1 (-58.5, 66.6) | 0.899 | -1.1 (-2.6, 0.4) | 0.137 | -0.02 (-0.10, 0.07) | 0.709 |
| V9 vs V8c | 12.6 (-51.8, 77.1) | 0.700 | -0.1 (-1.6, 1.4) | 0.865 | -0.01 (-0.09, 0.08) | 0.886 |

**Supplemental table S22**. Longitudinal model results for IgG kinetics. Values are sorted by HIV status and are shown as the median value across all respective individuals, and standard deviation (SD) across individual fits. RSE – relative standard error that represents the fitting uncertainty amongst the population of parameters, *μA_spike_* (*μA_RBD_*) – model-predicted spike (RBD) IgG production rate, *γA_spike_* (*γA_RBD_*) – model-predicted spike (RBD) IgG decay rate.

|  |  | **Dose 1 and 2** | | | **Dose 3** | | |
| --- | --- | --- | --- | --- | --- | --- | --- |
|  |  | **HIV^–^** | **PWH** | RSE (%) | **HIV^–^** | **PWH** | RSE (%) |
|  |  | Median (SD) | Median (SD) |  | Median (SD) | Median (SD) |  |
| Spike IgG | *μA_spike_* | 3.17 (0.08) | 3.18 (0.14) | 27.1 | 0.625 (0.003) | 0.643 (0.004) | 28.7 |
|  | *γA_spike_* | 0.024 (0.001) | 0.024 (0.001) | 24.8 | 0.0086 (0.001) | 0.0087 (0.0003) | 28.4 |
| RBD IgG | *μA_RBD_* | 2.4 (0.1) | 2.4 (0.1) | 26 | 0.81 (0.02) | 0.82 (0.02) | 28 |
|  | *γA_RBD_* | 0.0216 (0.002) | 0.0215 (0.0002) | 13.5 | 0.011 (0.002) | 0.011 (0.002) | 28 |

**Supplemental table S23**. Longitudinal model results for cytokine kinetics. Values are sorted by HIV status and are shown as fit-estimated median value across all respective individuals, and standard deviation (SD) across individual fits (in brackets). RSE – relative standard error representing the fitting uncertainty amongst the population of parameters. *T_D_* – estimated doubling time from the mean longitudinal response for each time point given, where the average time of D2 and D3 post-D1 is used for the doubling time estimate, *μ_F_* – model-predicted IFN-γ production rate, *μ_I_* – model-predicted IL-2 production rate.

|  |  | **HIV^–^** | **All PWH** | **IR** | **INR** | **RSE (%)** |
| --- | --- | --- | --- | --- | --- | --- |
| **IFN-γ** | Production rate, *μF* (days^–1^) | 0.023 (0.016) | 0.041 (0.12) | 0.12 (0.13) | 0.010 (0.033) | 36.4 |
|  | *T_D_* following D1 (days) | 162 | 58 | 47 | 150 | – |
|  | *T_D_* following D2 (days) | 208 | 109 | 101 | 196 | – |
|  | *T_D_* following D3 (days) | 498 | 389 | 381 | 477 | – |
| **IL-2** | Production rate, *μI* (days^–1^) | 0.12 (0.05) | 0.23 (0.18) | 0.27 (0.19) | 0.060 (0.05) | 21.0 |
|  | *T_D_* following D1 (days) | 45 | 47 | 48 | 46 | – |
|  | *T_D_* following D2 (days) | 102 | 166 | 217 | 83 | – |
|  | *T_D_* following D3 (days) | 384 | 849 | 1,230 | 258 | – |

**Supplemental table S24**. Multi-colour flow cytometry panel used for B cell staining.

| **#** | **Antibody or tetramer** | **Fluorochrome** |
| --- | --- | --- |
| 1 | Mouse anti-human CD45 | BUV805 |
| 2 | Mouse anti-human CD19 | BV650 |
| 3 | Mouse anti-human CD20 | APC-H7 |
| 4 | Mouse anti-human CD10 | BV510 |
| 5 | Mouse anti-human IgG | PE-Cy7 |
| 6 | Mouse anti-human CD11c | BUV395 |
| 7 | Mouse anti-human CD3 | BV570 |
| 8 | Mouse anti-human IgD | BV605 |
| 9 | Mouse anti-human IgM | BV711 |
| 10 | Mouse anti-human CD27 | BV785 |
| 11 | Mouse anti-human CD21 | PE/Dazzle594 |
| 12 | Mouse anti-human CD38 | APC/Fire810 |
| 13 | Mouse anti-human CD71 | Alexa Fluor 700 |
| 14 | Mouse anti-human IgA | VioBlue |
| 15 | SARS-CoV-2 S1 wild type (WT) | PE |
| 16 | SARS-CoV-2 S2P (WT) | PE-Cy5.5 |
| 17 | SARS-CoV-2 RBD (WT) | BV421 |
| 18 | SARS-CoV-2 NTD (WT) | AF488 |
| 19 | Zombie NIR fixable Live/Dead dye | |

**Supplemental table S25**. Sequences of IPDA primers and probes.

| **ID** | **Oligo** | **Sequence** | **HXB2 Coordinates** | **Dye** |
| --- | --- | --- | --- | --- |
| **IPDA**  **Ψ** | Forward primer | CAGGACTCGGCTTGCTGAAG | 692 → 711 | – |
|  | Reverse primer | GCACCCATCTCTCTCCTTCTAGC | 775 → 797 | – |
|  | MGB probe | TTTTGGCGTACTCACCAGT–MGBNFQ | 758 → 740 | FAM |
| **IPDA**  **Env** | Forward primer | AGTGGTGCAGAGAGAAAAAAGAGC | 7736 → 7759 | – |
|  | Reverse primer | GTCTGGCCTGTACCGTCAGC | 7832 → 7851 | – |
|  | MGB probe | CCTTGGGTTCTTGGGA–MGBNFQ | 7781 → 7796 | VIC |
|  | Hypermutant MGB probe | CCTTAGGTTCTTAGGAGC–MGBNFQ | 7781 → 7798 | – |
| **Alt**  **Ψ** | Forward primer | TCTCGACGCAGGACTCG | 684 → 700 | – |
|  | Reverse primer | TACTGACGCTCTCGCACC | 793 → 810 | – |
|  | Probe | CTCTCTCCTTCTAGCCTC–MGBNFQ | 772 → 789 | FAM |
| **RPP30** | Forward primer | GATTTGGACCTGCGAGCG | 474 → 491,  9559 → 9576 | – |
|  | Reverse primer | GCGGCTGTCTCCACAAGT | 6250 → 6267 | – |
|  | MGB probe | CTGACCTGAAGGCTCT–MGBNFQ | 1160 → 1175 | VIC |
| **RPP30 Shear** | Forward primer | CCATTTGCTGCTCCTTGGG | Chr 10, 7882-7900 | – |
|  | Reverse primer | CATGCAAAGGAGGAAGCCG | Chr 10, 7951-7969 | – |
|  | MGB probe | AAGGAGCAAGGTTCTATTGTAG–MGBNFQ | Chr 10, 7906-7927 | FAM |

**Supplemental table S26**. Longitudinal modelling of random effects and relative standard error (RSE) of random effects for the population parameters from each fit. *μ_F_* (*μ_I_*) – model-predicted IFN-γ (IL-2) production rates, *μ_RBD+_* (*μ_NTD+_*) – model-predicted RBD (NTD) B cell priming rate, *μA_spike_* (*μA_RBD_*) – model-predicted spike (RBD) IgG production rate, *γA_spike_* (*γA_RBD_*) – model-predicted spike (RBD) IgG decay rate, AIC – Akaike information criterion, BIC – Bayesian information criterion.

|  | | | **Random effects** | **RSE (%)** |
| --- | --- | --- | --- | --- |
| **Single-stage fit** | **Doses 1-3** | *μF* | 1.69 | 20.1 |
|  |  | *μI* | 0.87 | 23.1 |
|  |  | *μRBD+* | 0.4 | 47.8 |
|  |  | *μNTD+* | 0.52 | 48.5 |
|  |  | AIC | 1974 | |
|  |  | BIC | 2097 | |
| **Two-stage fit** | **Doses 1-2** | *μA_spike_* | 0.13 | 98 |
|  |  | *γA_spike_* | 0.22 | 67 |
|  |  | *μA_RBD_* | 0.12 | 48 |
|  |  | *γA_RBD_* | 0.13 | 68.6 |
|  |  | AIC | 981 | |
|  |  | BIC | 1099 | |
|  | **Dose 3** | *μA_spike_* | 0.15 | 57.7 |
|  |  | *γA_spike_* | 0.15 | 151 |
|  |  | *μA_RBD_* | 0.13 | 89.7 |
|  |  | *γA_RBD_* | 0.27 | 124 |
|  |  | AIC | 1060 | |
|  |  | BIC | 1185 | |

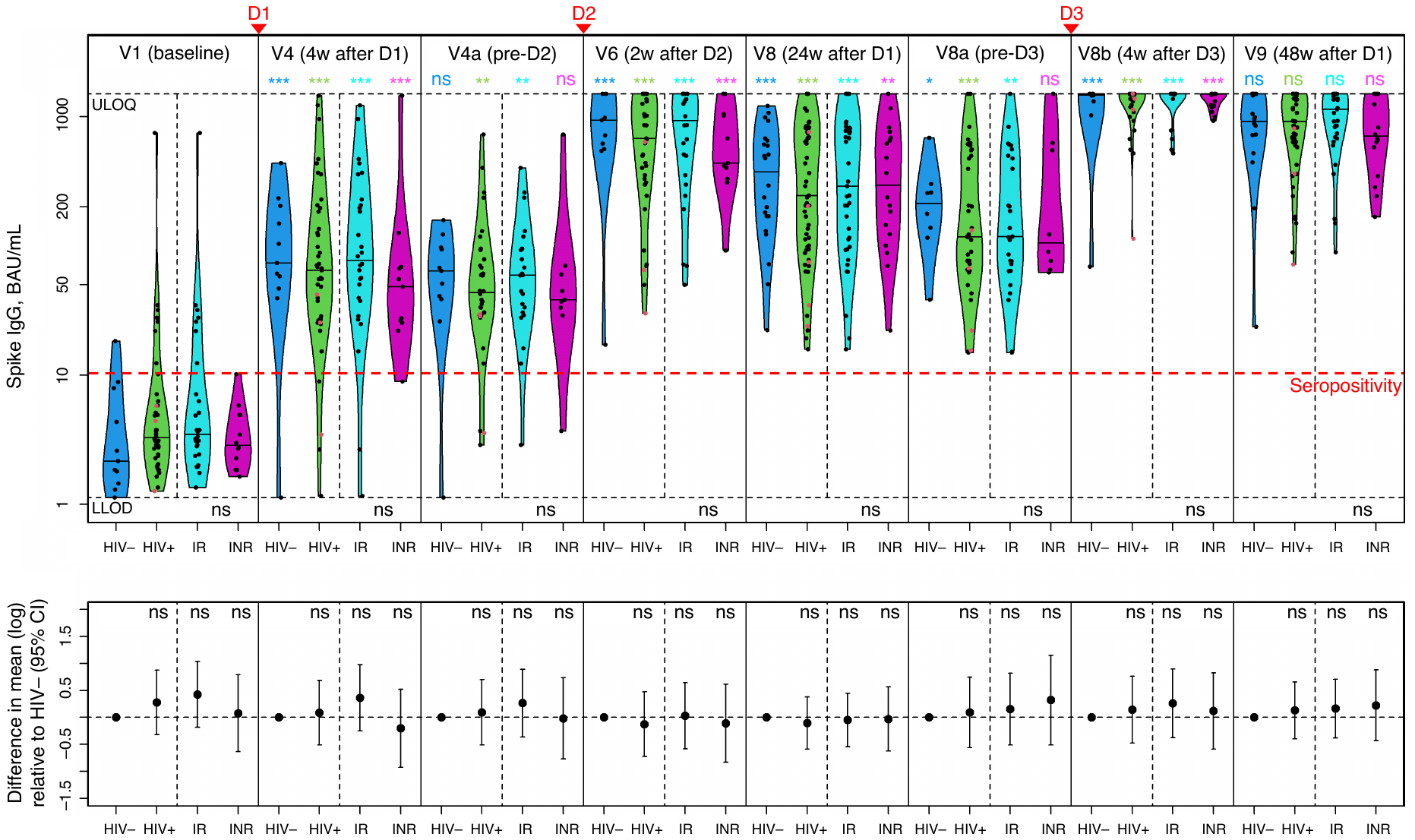

**Supplemental Figure S1**. **Three doses of COVID-19 vaccines elicit equally high levels of serum anti-spike IgG that increase with each dose.** The top panel shows a violin plot with medians for IgG concentrations (BAU/mL) in HIV^–^ (blue), total PWH (green), IRs (cyan), INRs (purple); N^+^/post-COVID-19 samples are excluded. The bottom panel shows adjusted log*_e_* mean differences between PWH and HIV^–^ individuals, based on mixed effects linear regression. *P* values for log*_e_* mean differences between HIV^+^ and HIV^–^ samples at each timepoint are shown at the top of the bottom panel: *p*<0.001 (***), *p*<0.01 (**), *p*<0.05 (*), *p*≥0.05 (ns = ‘not significant’). *P* values for within-group/subgroup changes between neighbouring timepoints are shown above their respective violin plots and are colour-coded accordingly. *P* values for IR vs INR differences are shown at the bottom of the top panel. The lower limit of detection (LLOD) is 1.13 BAU/mL; the upper limit of quantification (ULOQ): 1,501 BAU/mL; the seropositivity threshold: 11.3 BAU/mL. LLV participants are shown as red dots. Vaccination timepoints (D1, D2, D3) are indicated with red arrowheads.

**Supplemental Figure S2. Correlation between different datasets related to SARS-CoV-2 neutralization.** **(A-B)** Scatterplots showing Spearman correlation between live SARS-CoV-2 50% neutralization titers (NT50) and anti-RBD **(A)** or anti-NTD **(B)** B cells. See also Fig. 3A,B. **(C)** Spearman correlation between snELISA data and live SARS-CoV-2 NT50. See also Fig. 3A,C.

**Supplemental Figure S3**. **Post-vaccination increase in HIV viral load ‘blips’ in PWH.** Longitudinal changes in the percentage of samples with detectable HIV viral load (above 40 copies/mL) – for total PWH (top), IRs (middle) and INRs (bottom). The timing of administration of each COVID-19 vaccine dose (D1, D2, D3) is indicated with red arrowheads.

**Supplemental Figure S4**. **HIV gag/nef-specific T cells do not change in frequency following COVID-19 vaccination.** HIV-1 gag (left) and nef (right) responses from T cells at 24 weeks (V8) and 48 weeks (V9) post-D1 relative to the baseline, by the ELISpot assay and quantile regression analysis. Responses in PWH who were neither N^+^ or post-COVID-19, nor had a D4 prior to V9 (n=37) were measured as the number of cells that secrete IFN-γ (top), IL-2 (middle) or both cytokines (‘Dual’, bottom) following stimulation with HIV-1 gag or nef peptide pools and expressed as spot-forming cells (SFC) per 10^6^ PBMC; horizontal bars indicate the median. LLV participants are shown in red. *P* values are not shown because none of the changes were significant. The timing of administration of each COVID-19 vaccine dose (D1-2, D3) is indicated with red arrowheads.

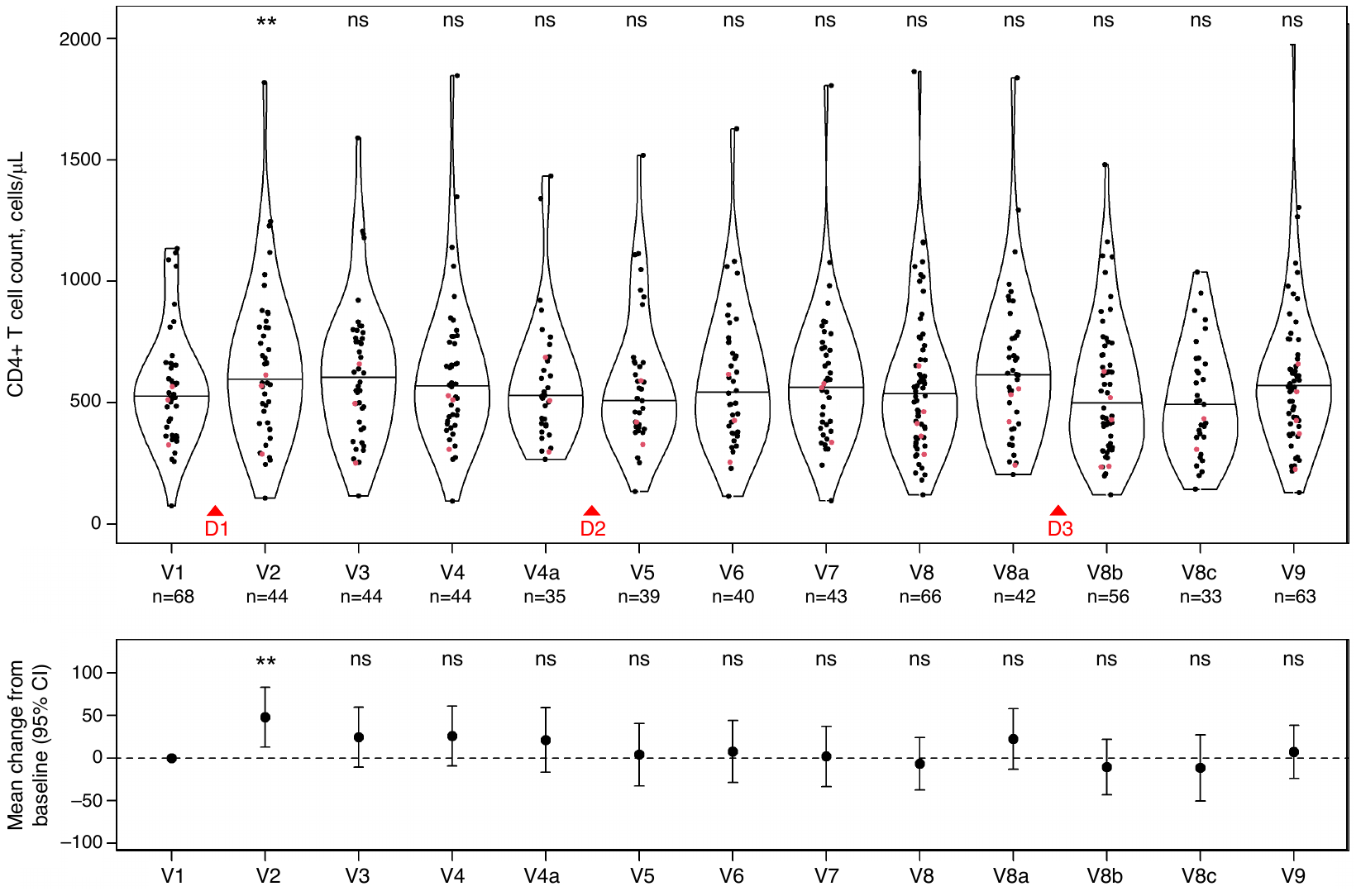

**Supplemental Figure S5**. **CD4^+^ T cell count in PWH increased transiently after the first vaccine dose.** Longitudinal changes in CD4^+^ T cell count (cells/μL) in PWH based on mixed effects linear regression, with horizontal bars indicating the mean. The red dots represent LLV participants. The *p* values for changes between neighbouring time points are shown at the top: *p*<0.01 (**), *p*≥0.05 (ns = ‘not significant’). The bottom panel shows mean changes relative to the baseline, with *p* values for differences between each time point and the baseline. The timing of administration of each COVID-19 vaccine dose (D1, D2, D3) is indicated with red arrowheads.

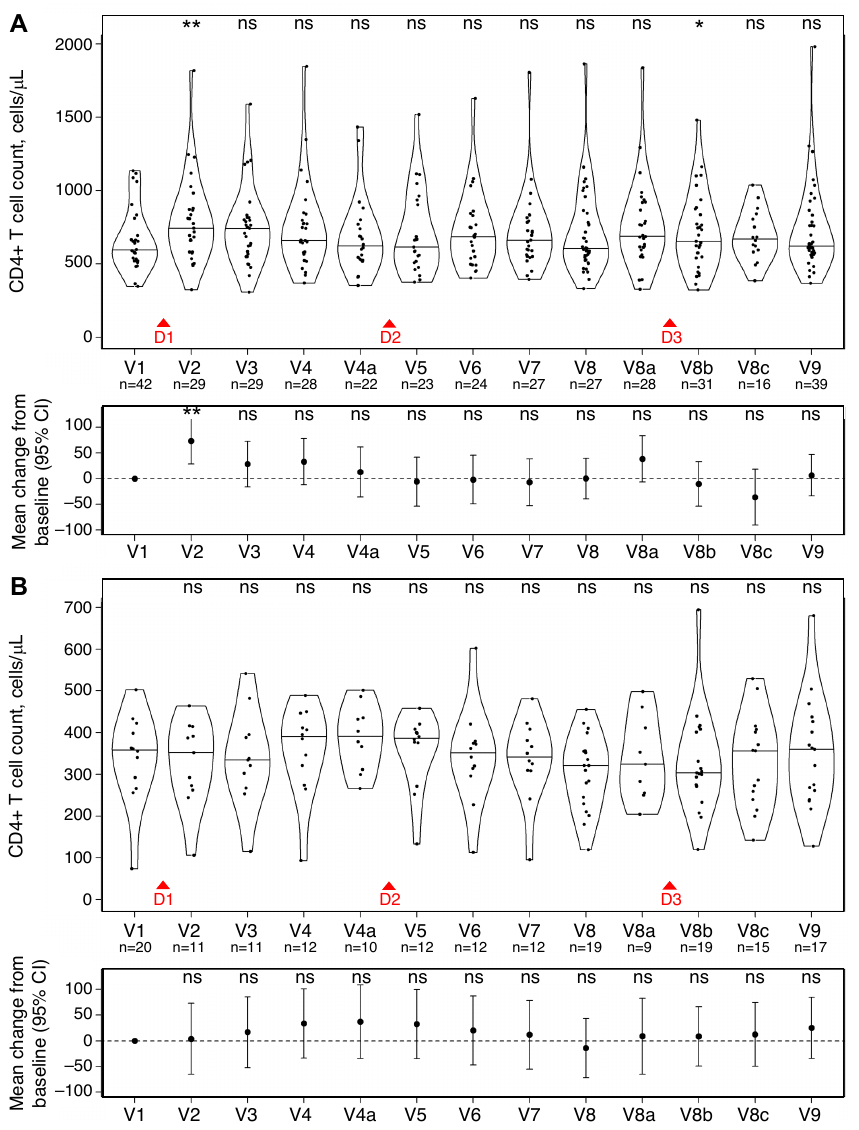

**Supplemental Figure S6**. **CD4^+^ T cell count increased transiently after the first vaccine dose in IRs but not in INRs.** Longitudinal changes in CD4^+^ T cell count (cells/μL) in PWH who are IR **(A)** and INR **(B)** based on mixed effects linear regression, with horizontal bars indicating the mean. The red dots represent LLV participants. The *p* values for changes between neighbouring time points are shown at the top: *p*<0.01 (**), *p*≥0.05 (ns = ‘not significant’). The two bottom panels show mean changes relative to the baseline, with *p* values for differences between each time point and the baseline. The timing of administration of each COVID-19 vaccine dose (D1, D2, D3) is indicated with red arrowheads.

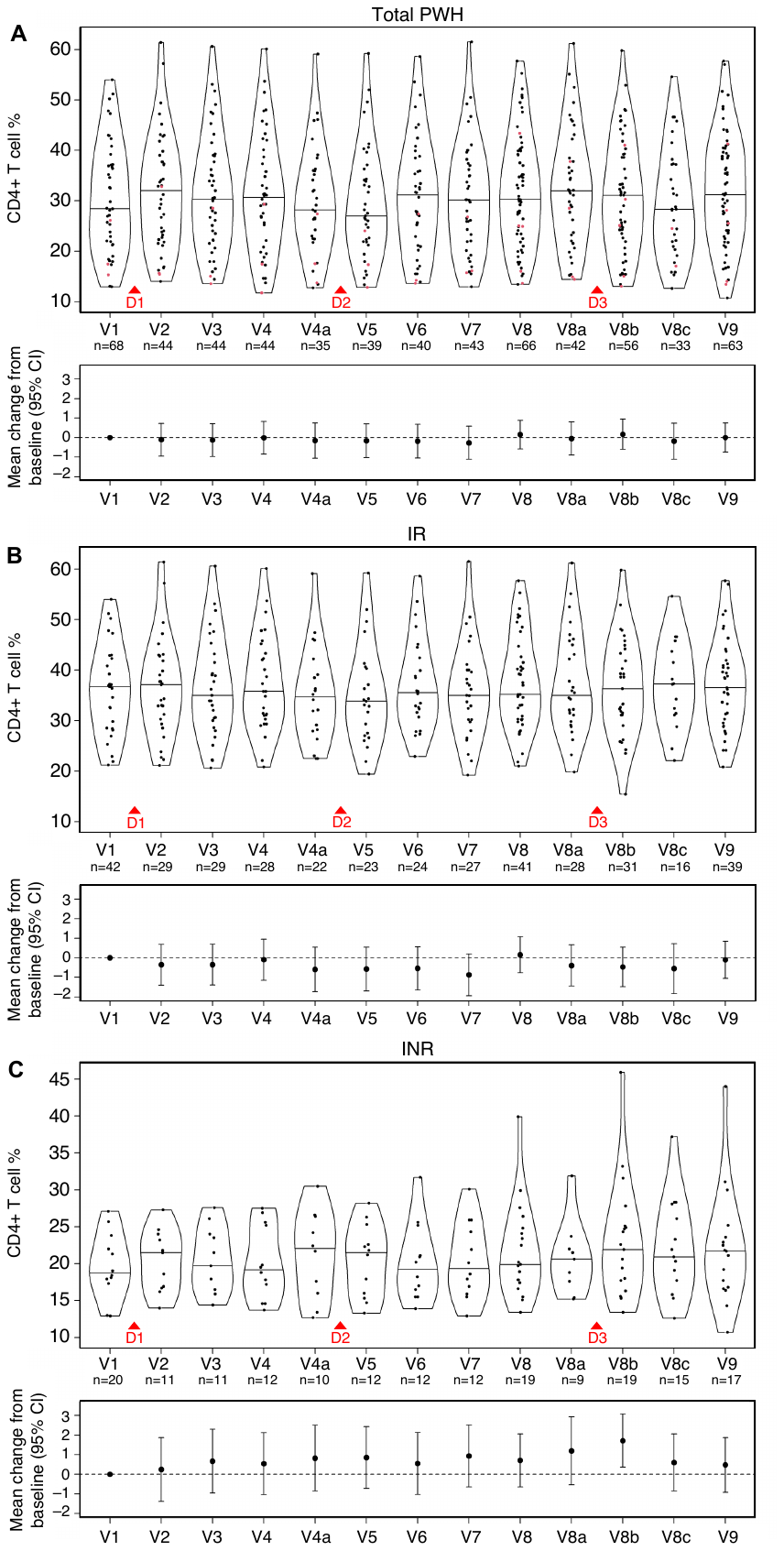

**Supplemental Figure S7**. **CD4^+^ T-cell percentage did not change in PWH following COVID-19 vaccination.** Longitudinal changes in CD4^+^ T-cell percentage in total PWH **(A)**, IRs **(B)** and INRs **(C)** based on mixed effects linear regression, with red dots representing LLV participants. Horizontal bars indicate the mean. Each bottom panel shows mean changes relative to the baseline. *P* values are not shown because none of the changes were significant. The timing of administration of each COVID-19 vaccine dose (D1, D2, D3) is indicated with red arrowheads.

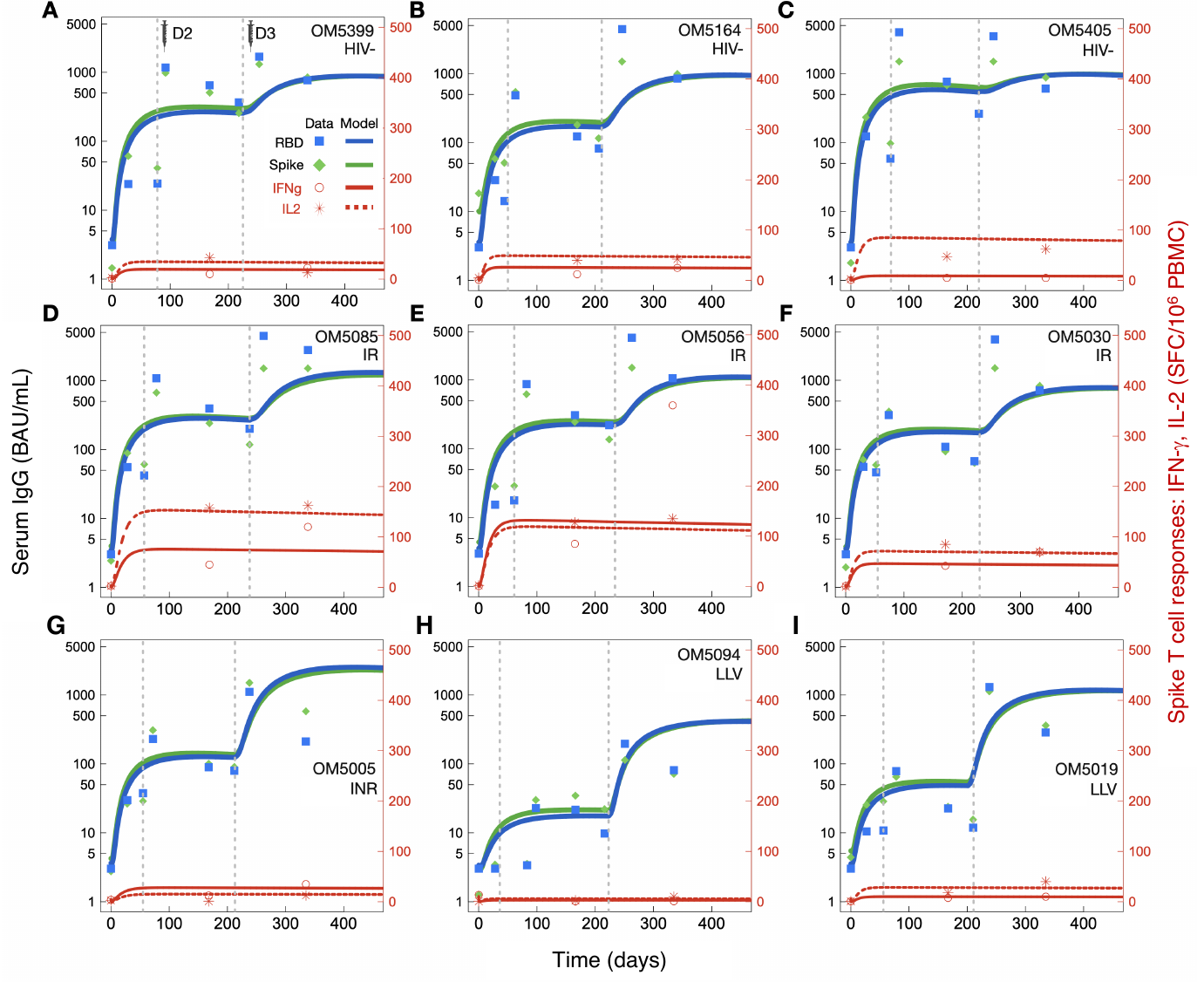

**Supplemental Figure S8**. **Individual longitudinal Eq. 1 model fit examples.** Left-hand Y-axes show serum IgG measures (BAU/mL), right-hand (red) axes are ELISpot measures for IFN-γ and IL-2 anti-spike T cell responses in SFC/10^6^ PBMC units. Grey vertical dashed lines show the timings of D2 and D3. Also see Fig. 7. **(A-C)** HIV^–^ individual fits. **(D-F)** IR individual fits. **(G)** An INR fit. **(H-I)** LLV individual fits.

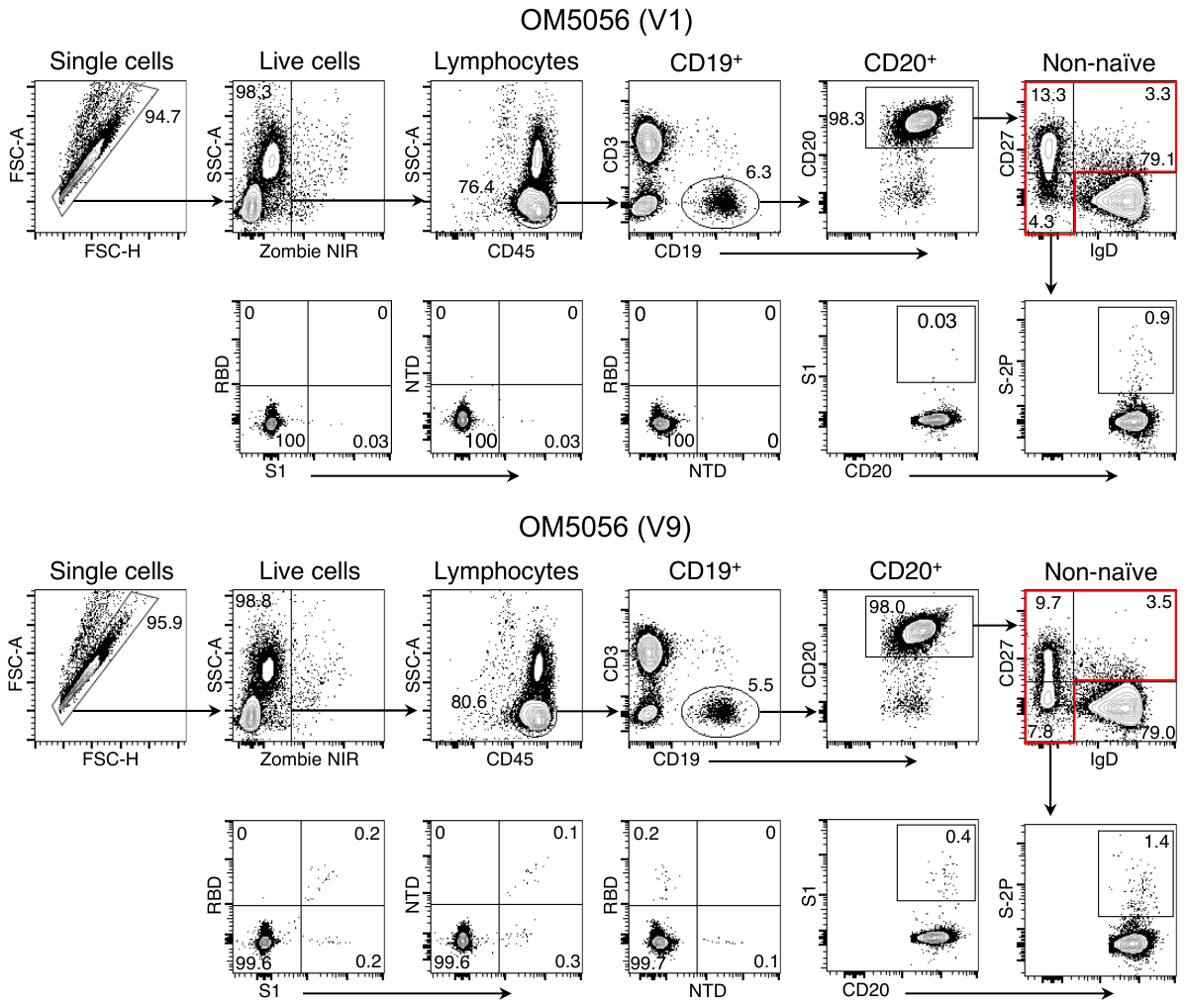

**Supplemental Figure S9.** Example spectral flow cytometry plots for OM5056 (V1 and V9) showing our gating strategy for the identification of SARS-CoV-2 RBD/NTD-specific B cells.

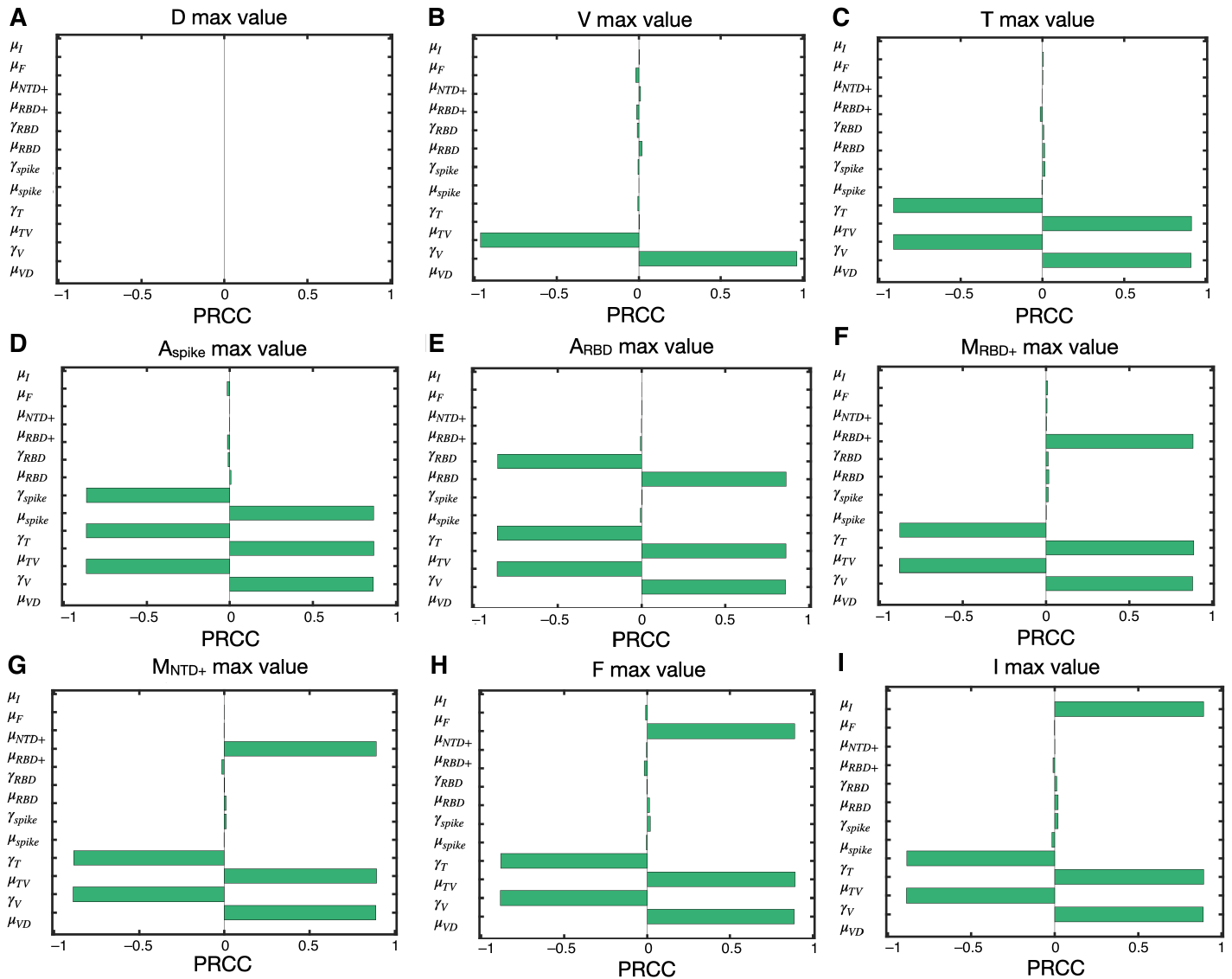

**Supplemental Figure S10. Partial rank correlation coefficient (PRCC) sensitivity analysis with Latin hypercube sampling (LHS) for model parameters.** The maximum value of each state variable is selected as the model output. PRCC values close to the maximum value of 1.0 indicate that the model output is highly sensitive to variation in that parameter, with values greater than 0.5 considered significant.

**SUPPLEMENTAL** **REFERENCES**

1.	Korosec, C.S., Farhang-Sardroodi, S., Dick, D.W., Gholami, S., Ghaemi, M.S., Moyles, I.R., Craig, M., Ooi, H.K., and Heffernan, J.M. (2022). Long-term durability of immune responses to the BNT162b2 and mRNA-1273 vaccines based on dosage, age and sex. Sci Rep *12*, 21232. 10.1038/s41598-022-25134-0.

2.	Moyles, I.R., Korosec, C.S., and Heffernan, J.M. (2022). Determination of significant immunological timescales from mRNA-LNP-based vaccines in humans. medRxiv, 2022.2007.2025.22278031. 10.1101/2022.07.25.22278031.

3.	Farhang-Sardroodi, S., Korosec, C.S., Gholami, S., Craig, M., Moyles, I.R., Ghaemi, M.S., Ooi, H.K., and Heffernan, J.M. (2021). Analysis of Host Immunological Response of Adenovirus-Based COVID-19 Vaccines. Vaccines *9*, 861.

4.	McHeyzer-Williams, L.J., and McHeyzer-Williams, M.G. (2005). Antigen-specific memory B cell development. Annu Rev Immunol *23*, 487-513. 10.1146/annurev.immunol.23.021704.115732.

5.	Sher, A., Niederer, S.A., Mirams, G.R., Kirpichnikova, A., Allen, R., Pathmanathan, P., Gavaghan, D.J., van der Graaf, P.H., and Noble, D. (2022). A Quantitative Systems Pharmacology Perspective on the Importance of Parameter Identifiability. Bull Math Biol *84*, 39. 10.1007/s11538-021-00982-5.

6.	McKay, M.D., Beckman, R.J., and Conover, W.J. (1979). A Comparison of Three Methods for Selecting Values of Input Variables in the Analysis of Output from a Computer Code. Technometrics *21*, 239-245. 10.2307/1268522.

7.	Wu, J., Dhingra, R., Gambhir, M., and Remais, J.V. (2013). Sensitivity analysis of infectious disease models: methods, advances and their application. J R Soc Interface *10*, 20121018. 10.1098/rsif.2012.1018.
